# Impact of Post-Translational Modification on MHC Peptide Binding and TCR Engagement

**DOI:** 10.1101/2023.03.02.530810

**Authors:** Joey J. Kelly, Nathaniel Bloodworth, Qianqian Shao, Jeffery Shabanowitz, Donald Hunt, Jens Meiler, Marcos M. Pires

## Abstract

The human major histocompatibility complex (MHC) plays a crucial role in the presentation of peptidic fragments from proteins; these peptides can be derived from self-proteins or from non-human antigens, such as those produced by viruses or bacteria. To prevent cytotoxicity against healthy cells, thymocytes expressing T cell receptors (TCRs) that recognize self-peptides are removed from circulation in a process called negative selection. However, post-translational modifications (PTMs) are largely excluded from negative selection; this feature opens the door to the possibility that PTMs directly contribute to the development of autoreactive T cells and subsequent autoimmune diseases. Despite it being well-established that PTMs are prevalent in peptides presented on MHCs, the exact mechanisms by which PTMs influence the antigen presentation machinery remains poorly understood. In our work, we introduce chemical modifications mirroring PTMs onto peptides to systematically investigate their impact on MHC binding and TCR recognition. Our findings reveal the numerous ways PTMs alter antigen presentation, which could have implications for tumor neoantigen presentation.

## Introduction

To maintain homeostasis, the human immune system must efficiently recognize and destroy cells that have accumulated genetic mutations or have been invaded by pathogenic microorganisms.^1, 2^ Self-identification to the immune system is a primary mechanism that is deployed by human cells to flag the presence of non-self-proteins or proteins produced by genetic lesions.^3^ To this end, presentation of peptidic fragments from non-self-proteins *via* the major histocompatibility complex (MHC) to surveying immune cells serves as a key system to recognize diseased cells (**Figure 1**).^4^ MHC is present in the membrane of every nucleated cell and is responsible for presenting both antigenic and endogenous protein fragments to the extracellular space. Binding of peptide-MHC (pMHC) complexes by T cells, which are equipped with T cell receptors (TCRs)^5^, initiates an immune cell response; the nature of the response depends on the type of the T cell (primarily CD4+ and CD8+ cells).^6^

**Figure 1.**
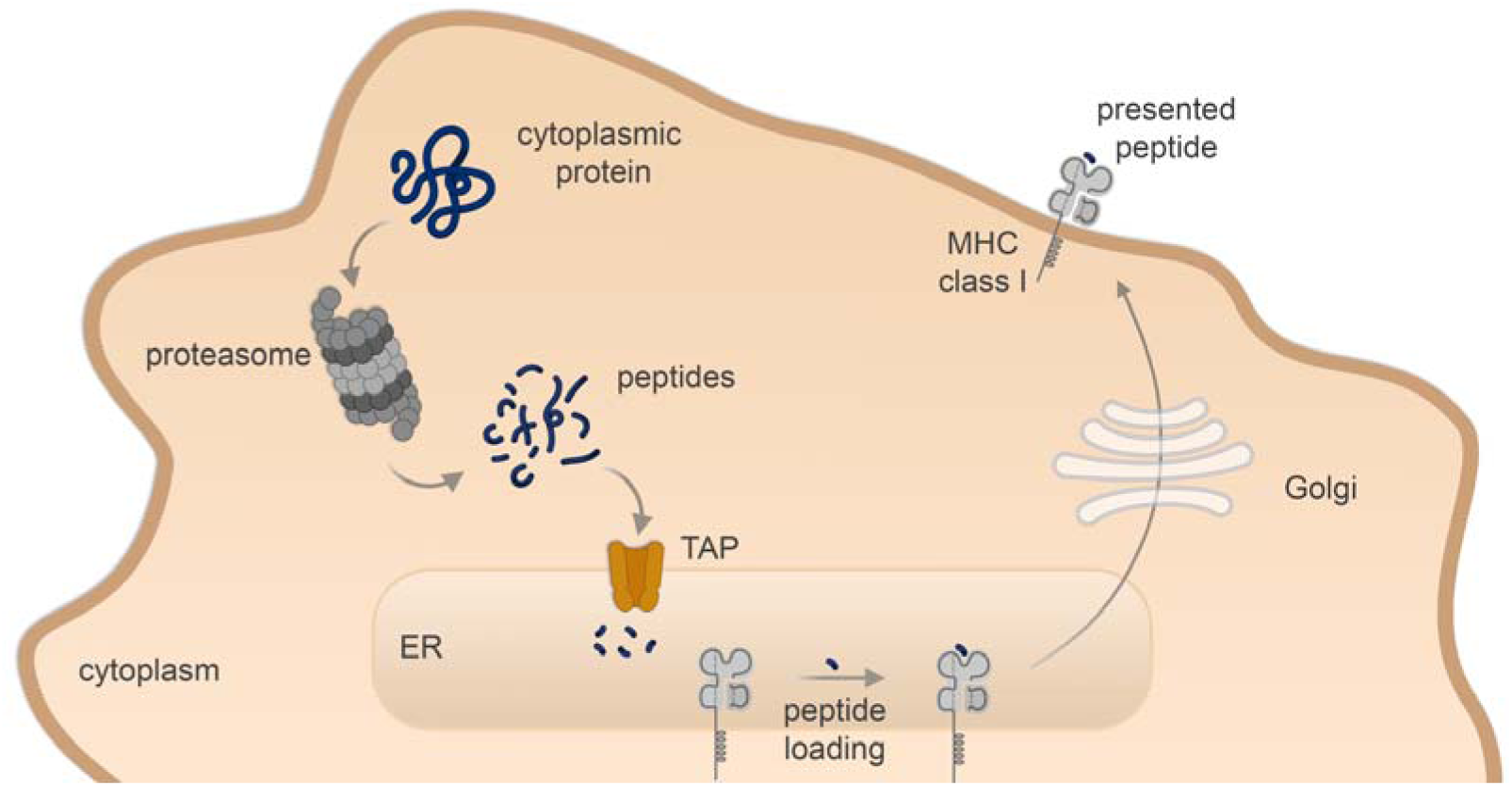
Schematic representation of the process involving the proteolytic processing of cytosolic protein into peptides. Subsequent steps lead to the loading of the peptides onto MHC molecules that are then transported for presentation on the cell surface.

Critically, the binding of pMHC by TCRs must operate with a high level of selectivity since the recognition of self-peptides in healthy cells can result in cellular injury.^7^ Through negative selection, thymocytes expressing TCRs that bind tightly to self-peptides are removed from circulation.^5, 8^ In theory, changes to the primary sequence of a protein could yield autoreactive pMHC as long as that primary sequence was not part of the negative selection process. Some primary sequence changes can be permanent (e.g., amino acid change), while others can be transient, such as post-translational modifications (PTMs). In healthy states, enzymatic PTMs are installed by specific sets of enzymes to modulate protein activity, localization, and interaction of the protein with other cellular components.^9^ Because PTMs are covalent modifications, most are stable enough to persist through proteasomal processing; therefore, the resulting modified peptide fragments can potentially be presented in the context of MHC for immune surveillance.^10^

Remarkably, human proteins containing PTMs were shown to be excluded from negative selection, despite the potential that pMHC with PTM-modified peptides could yield autoreactive cells.^11^ It was hypothesized that the reason was the relatively low abundance of PTMs in healthy and young adults in the prime phase of negative selection, compared to the unmodified parent proteins. Past adolescence, thymus involution has the potential to contribute to age-related autoimmune diseases.^12, 13^ At the same time, a myriad of factors (balance of PTM writers and erasers in diseased states, cellular stresses, or aging) can dramatically increase the prevalence of PTMs.^14^ Combined, the increased levels of pMHC complexes bearing PTMs means that a higher level of potentially autoreactive cells are present.^15–21^ In other words, peptides with PTM modifications could potentially be interpreted by the immune system as nonself, and, therefore, induce a pathogenic autoimmune response.^22–24^ A conclusive example of this phenomenon was recently described in which cysteine carboxyethylation by a cellular metabolite resulted in a pathogenic neoantigen presented on pMHC that resulted in autoreactive T cell responses.^25^

A wide range of PTMs have been previously found in pMHCs.^26–34^ As a prominent example, increased conversion of arginine to citrulline in the myelin sheath has been shown to lead to the development of self-reactive T cells that exacerbate the progression of autoimmune diseases such as Rheumatoid Arthritis (RA) and Multiple Sclerosis (MS).^35–39^ Additionally, there is mounting evidence that PTMs on peptides within pMHC complexes are prevalent in tumors.^40–43^ A recent report revealed that PTMs can shape the antigenic landscape.^43^ Despite the importance of PTMs to potentially generate autoreactive peptides, there are many aspects of PTM biology that remain poorly defined in this field. A primary barrier to better understanding the impact of PTMs on pMHC complexes has been the complexity in assigning the role of PTMs in the context of the many steps leading to MHC presentation. These roles can include changes in protein half-live, altered proteolytic processing to presentable peptides, or binding affinity of individual peptides to MHC molecules. Each of these factors on their own has the potential to impact levels and types of peptides displayed on MHCs; of which, binding affinity is a primary factor in defining the repertoire of peptides within pMHCs.^44^, given the number of possible peptides that can theoretically be loaded onto MHCs by the transporter associated with antigen processing (TAP).^44^ Herein, we sought to systematically isolate the impact of PTMs on the ability of the peptide to bind to the MHC molecules and be recognized by cognate T cells.

## Results and Discussion

In general, PTMs can be challenging to detect in the context of complex biological systems. Additionally, the levels of PTMs are dynamic and can shift in response to cellular queues.^45^ Traditional cellular assays (e.g., Western blots) do not usually establish the ratio of PTM proteins relative to their unmodified counterparts.^46^ These challenges become amplified when analyzing PTMs on peptides from pMHC complexes.^47^ Extraction of peptides from isolated MHCs involves difficult separation steps and downstream mass spectrometry analysis.^48^ On the other hand, a^48^ A top-down approach of treating cells with PTM-modified proteins also poses a challenge because exogenous proteins are not readily processed to yield peptides that are presented by MHC I molecules.^49^ Even if the peptides get processed, the ultimate pMHC presentation entails a combination of factors including peptide half-life^50^, uptake efficiency, processing by proteases, or peptide binding affinity to MHC molecules. Therefore, identifying how various naturally occurring PTMs impact peptide binding to MHCs and recognition by TCRs has not been carried out in a systematic way.

To overcome these challenges, we centered our approach on the RMA-S cell line that lacks the TAP importer.^51, 52^ TAP is essential for delivering peptides from the cytosol to the endoplasmic reticulum. Once in the endoplasmic reticulum, high affinity binding of peptides to MHC leads to stable associations of this complex and robust presentation on the cell surface. In contrast, TAP deficient cells have inherently low levels of MHC class I expression on the cell surface. Lowering the temperature (22-26 °C) allows for proper expression of MHC class I that are loaded with low affinity peptides or are empty (**Figure S1**). Cell treatment with peptides that bind MHC with high affinity results in pMHC complexes with thermostability and high surface presentation. Effectively, peptides in the extracellular space displace low binding peptides or associate with empty MHC molecules. Upon the elevation of the temperature to 37 °C, low affinity binders dissociate from the pMHC complexes leading to internalization and degradation of the associated MHC molecules. In contrast, supplementing a high-affinity MHC binding peptide to the culture medium at 26 °C enables the stabilization of pMHC on the surface of the cell when the temperature is raised to 37 °C.

The quantity of MHC on the cell surface correlates with the binding affinity of the peptide to the MHC molecule, which can be quantified *via* detection with a fluorescently labeled anti-MHC antibody.^53^ Given these features of this well-established cell line, we anticipated that the RMA-S cells would serve the role of revealing the impact of PTMs on peptides that bind to MHC with high affinity.^54^ Moreover, there is considerable precedence for using RMA-S cell lines for isolating the affinity of a peptide for MHC class I in contexts other than PTMs.^55–59^ To start, we sought to identify a peptide from SARS-CoV-2 spike protein. We used NetMHCPan4.0, an algorithm that estimates the propensity of peptides to bind MHC, to select a peptide sequence (ESIVRFPNI, referred to as **sarsWT**) for stabilization of the specific MHC molecule (H-2K^b^) on the surface of RMA-S cells.

The peptide was synthesized using standard solid-phase peptide synthesis techniques. The culture medium of the RMA-S cells was supplemented with **sarsWT** (**Figure S2A**) at 26 °C to enable association of the peptide with surface bound MHCs. Following **sarsWT** incubation, the temperature was raised to 37 °C. Subsequently, the cells were incubated with APC conjugated anti-mouse H-2K^b^ antibody and analyzed by flow cytometry where fluorescence is expected to correspond with the amount of **sarsWT** bound to the MHC. The peptide SNFVSAGI (**cntPEP**) was used as a negative control since it has been reported to not interact appreciably with H-2K^b^.^60^ Various concentrations of **sarsWT** were used to optimize the RMA-S stabilization assay with the goal of increasing the number of pMHC molecules formed on the surface of the cells (**Figure S2**). We found an increased fluorescence signal corresponding to increasing concentrations of **sarsWT,** indicating that the peptide bound and stabilized the MHC on the surface of the cell. The amount of stable pMHC complexes that formed at the surface also saturated at 50 μM of peptide and showed an EC_50_ of 1.7 μM. These results confirmed the suitability of RMA-S to report on the stabilization of the pMHC complex by **sarsWT**.

Another parameter to consider is the time in which the MHC binding peptide has to associate with the MHC to reach MHC saturation. To evaluate this time frame, the workflow described above was conducted; however, the peptide incubation time at 37 °C was varied from 2 to 6 h (**Figure S3**). The results showed that the fluorescent signal increased with increased incubation periods. This is likely due to accumulation of stabilized pMHC complexes in the presence of excess high-affinity peptide. Nevertheless, a 6 h incubation at 37 °C provided the highest signal-to-noise ratio and was used for downstream assays.

With an optimized protocol in hand, we then synthesized a panel of peptides containing PTMs to investigate their potential impact on MHC binding (**Figure 2A**). We envisioned that modifying residues on **sarsWT** would be representative of how the selected modifications may impact binding of different MHC peptides. The specific PTMs chosen were a proline to hydroxyproline^61, 62^, arginine to citrulline^63^, and *N*-terminus acetylation^64^ modifications, as they are naturally occurring PTMs. By performing the RMA-S stabilization assay, we found that the citrullination and hydroxy-proline modifications had no significant effect on MHC binding, but the *N*-terminal acetylation modification completely disrupted MHC binding (**Figure 2B**). Gratifyingly, there is precedence for *N*-acetylated peptides to be displaced in MHC class I.^65, 66^ These results demonstrate that naturally occurring PTMs can alter pMHC complex formation and influence the antigen presentation pathway depending on the interface of the peptide and the MHC molecule (**Figure 3**).

**Figure 2.**
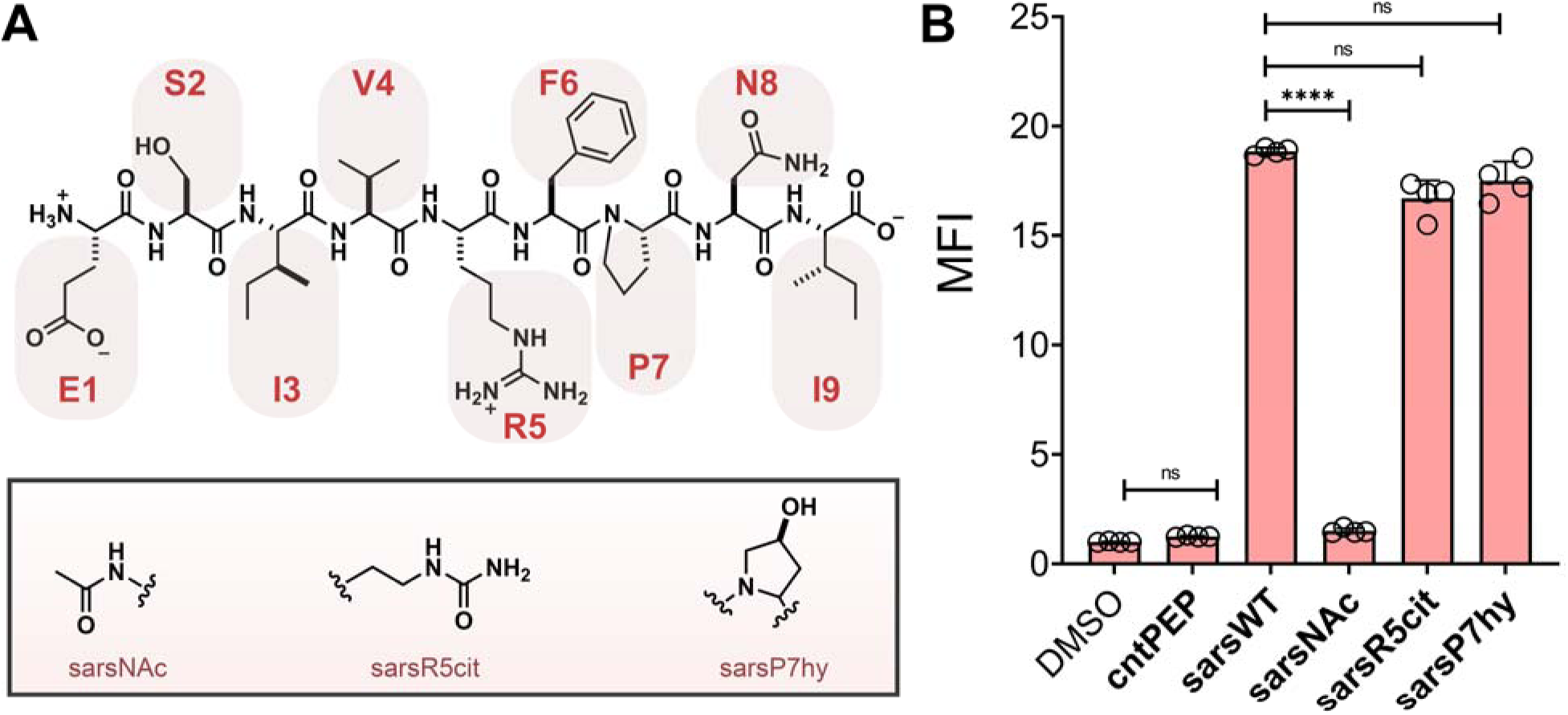
(A) Chemical structures of **sarsWT** and the PTM modified variants. (B) Flow cytometry analysis of RMA-S cells treated with specific peptide (20 μM) detected by APC conjugated anti-mouse H-2K^b^ antibody. Data are represented as mean ± SD (n= 3). P-values were determined by a two-tailed *t*-test (* p < 0.05, ** p < 0.01, *** p < 0.001, **** p < 0.0001, ns = not significant).

**Figure 3.**
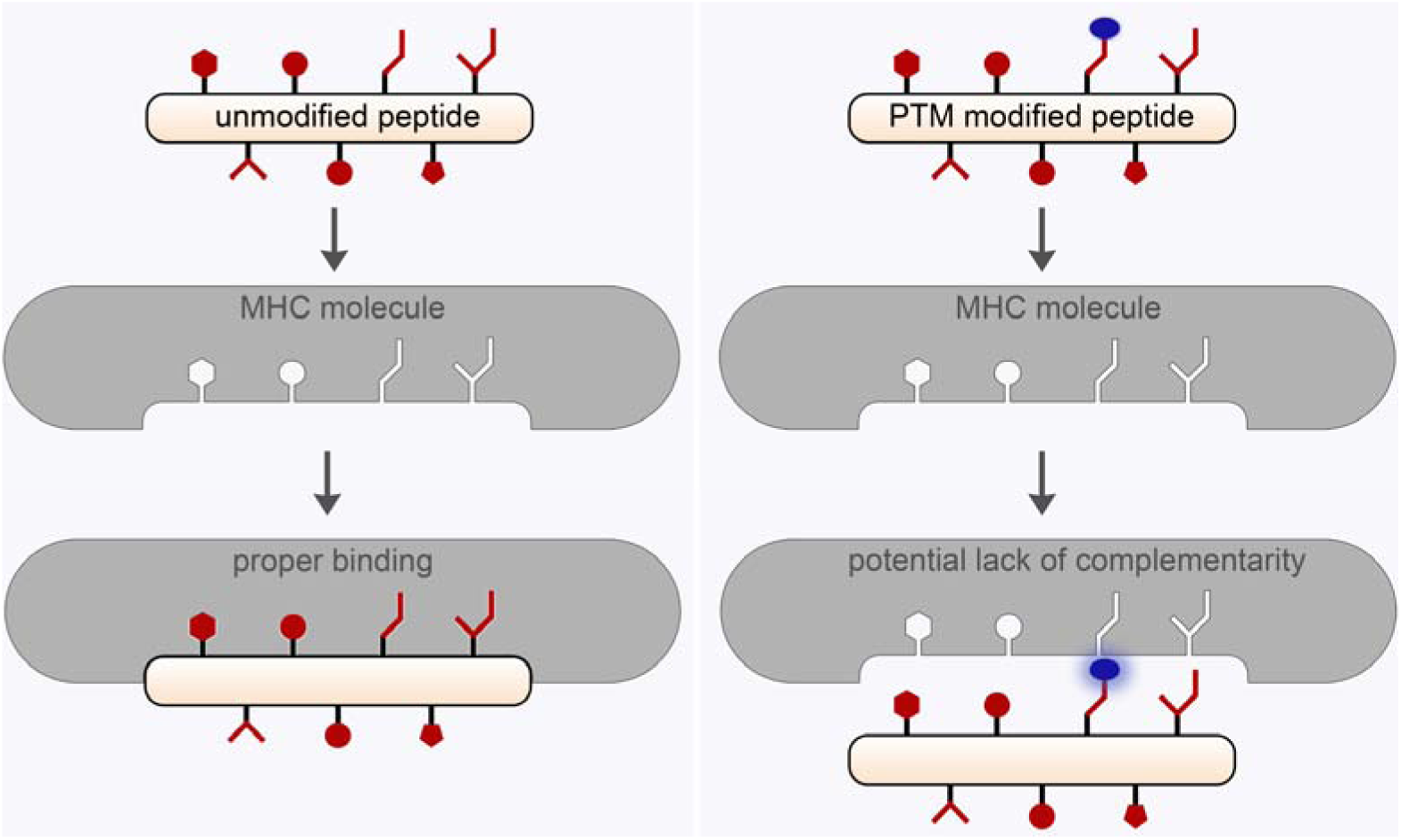
Schematic representation of how the orientation of the PTM within the presented peptide could negatively impact binding to MHC molecules (*right*) compared to the unmodified peptide (*left*).

We then shifted our focus to the peptide epitope SIINFEKL (**ovaWT**) from the model antigen ovalbumin (OVA) to interrogate how other PTMs impact MHC binding as well as TCR recognition.^67^ To this end, we synthesized a new library of OVA peptides containing PTMs on its serine and lysine residues and performed a concentration scan to assess MHC binding using the RMA-S stabilization assay (**Figure 4A**). Interestingly, there was a significant decrease in MHC binding when serine was phosphorylated despite predictions that this site does not contribute significantly to binding.^68^ Our results highlight the potential deficiency of current models to predict how PTMs could impact peptide binding to MHC molecules given how PTM products can be structurally distinct relative to the sidechains of the canonical amino acids.

**Figure 4.**
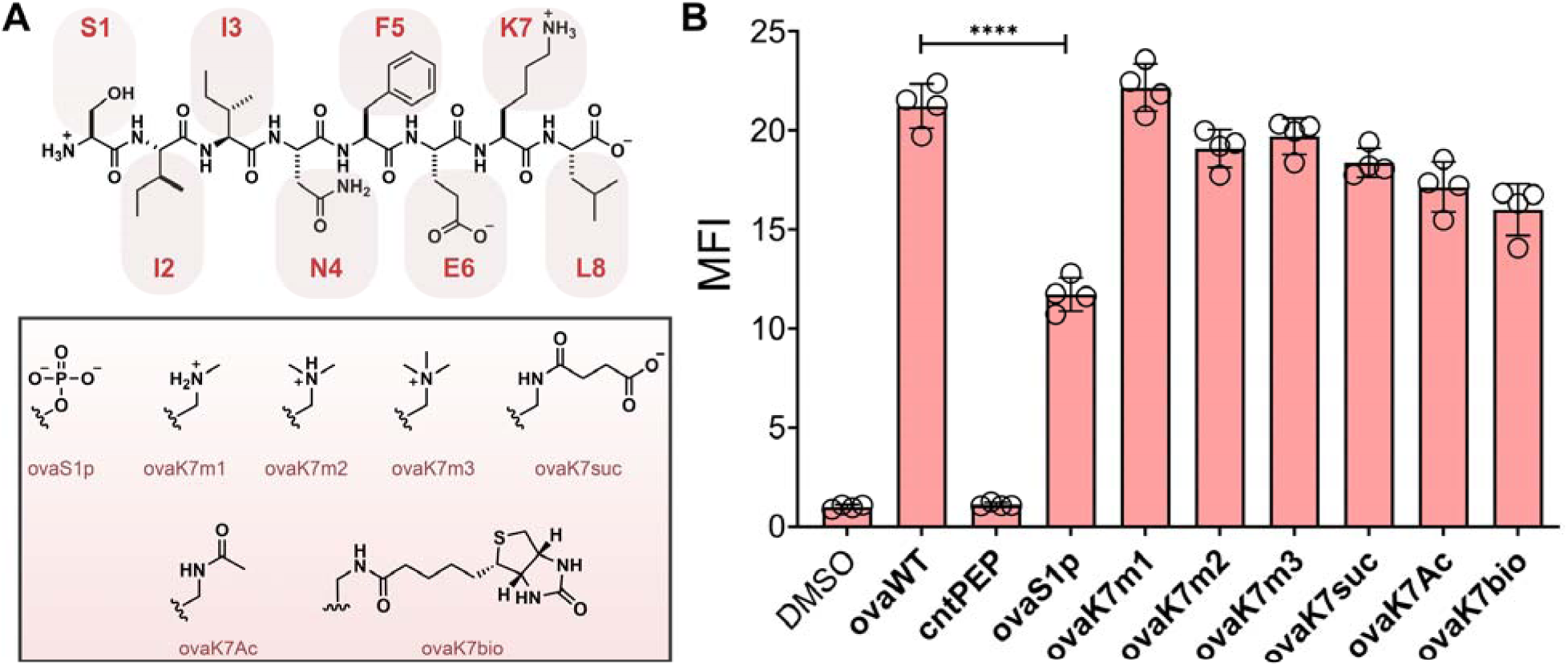
(A) Chemical structures of **ovaWT** and the PTM modified variants. (B) Flow cytometry analysis of RMA-S cells treated with specific peptide (20 μM) detected by APC conjugated anti-mouse H-2K^b^ antibody. Data are represented as mean ± SD (n= 3). P-values were determined by a two-tailed *t*-test (* p < 0.05, ** p < 0.01, *** p < 0.001, **** p < 0.0001, ns = not significant).

The sidechain of lysine on **ovaWT** was explored next. Lysine was decorated with different types of PTMs that included three different charge states. Previously, the same lysine on SIINFEKL had been altered with various groups including fluorophores and caging groups.^69–71^ Our data showed considerable accommodation of structural alterations to the lysine sidechain (**Figure 4B**). With larger modifications such as biotin, binding levels are altered but the change is not considerable. These findings can be explained by the solvent exposed nature of lysine 7 on the pMHC complex and the length of the lysine sidechain, which may reduce structurally unfavorable interactions with the MHC molecule (**Figure 3**).^72^ Critically, for **ovaWT** the anchor residues that are more important for the overall binding affinity to MHC molecules (Phe5 and Leu8) are not amendable to conventional PTMs.^73^ While PTMs on solvent exposed residues could potentially retain binding to MHCs, their overall impact may be more strongly reflected in the pMHC engagement with TCRs. TCRs recognize T cell exposed motifs of the bound peptide within pMHCs.^74, 75^ Therefore, PTMs at these positions may have a greater impact on T cell activation.

To assess the impact of **ovaWT** PTMs on TCR recognition, we utilized the B3Z T cell hybridoma cell line that contains an OVA specific TCR along with a NFAT-LacZ reporter gene that encodes for β-galactosidase on an IL-2 inducible promoter.^76^ Upon B3Z recognition of the OVA pMHC complex on RMA-S cells, IL-2 production promotes β-galactosidase expression, which can hydrolyze chlorophenol red-β-D-galactopyranoside (CPRG), and the resulting color change is representative of activation levels. Our results showed that the PTMs have a much more significant effect on TCR recognition than on peptide binding to MHC (**Figure 5A**). While relatively modest disruption of TCR recognition was seen for increasing the degree of methylation on the lysine residue, near complete disruption of TCR recognition was seen for all PTMs that imparted a change in charge of either the serine or lysine residue. Additionally, the reduction in TCR recognition for both the phosphoserine and biotinylated OVA cannot be fully explained solely by their decrease in MHC binding. Instead, the PTMs that are well accommodated for MHC binding could be displayed away from the binding cleft and alter engagement with TCRs on cognate T cells (**Figure 5B**).

**Figure 5.**
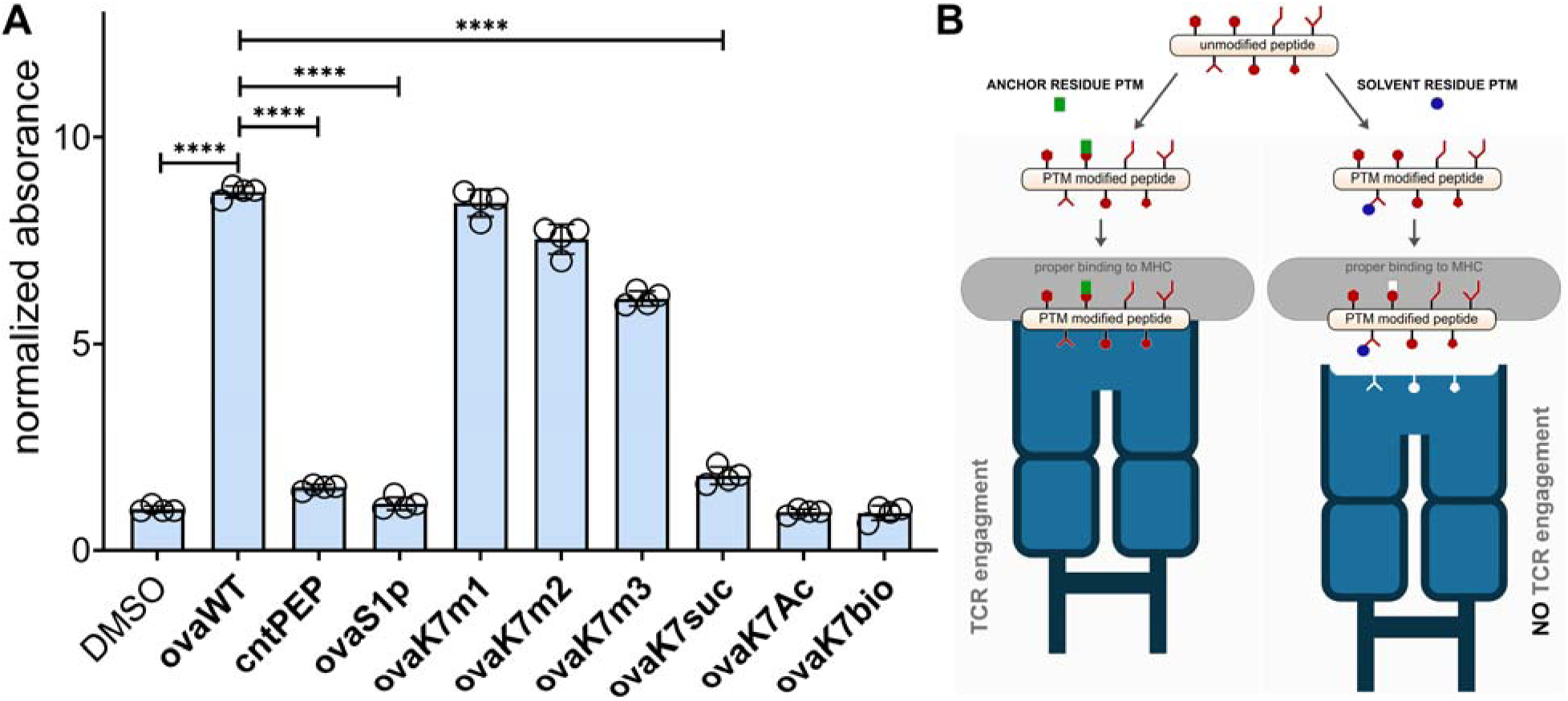
(A) RMA-S cells were incubated with peptide and B3Z T cells overnight at an effector to target ratio of 1:1. β-galactosidase expression was then measured *via* the colorimetric reagent CPRG on a plate reader at 570 nm.

To confirm the relevance of these findings in a broader context, we also performed the T cell activation assay on DC2.4 dendritic cells, which also express H-2K^b^. Unlike the RMA-S cells, peptides were incubated with DC2.4 cells at 37 °C and were expected to load onto MHC molecules if they display sufficient affinity towards H-2K^b^. Satisfyingly, the pattern of T cells activation with PTM modified **ovaWT** peptides showed a similar profile with DC2.4 cells as RMA-S cells (**Figure S4**). Our results confirmed that the impact of PTMs on T cell activation may be consistent across multiple cell types. Finally, we sought to investigate whether we could assess PTMs on a peptide that has potential pathological implications. To this end, a number of autoimmune diseases, including RA and MS, have been described to involve the citrullination of arginine.^77^ Citrullination is carried out by peptidylarginine deiminases (PADs) and higher levels of citrullinated proteins have been found in older mice relative to young mice.^78^ A shift in higher levels of citrullination past the full scope of negative selection could provide a pathway for autoreactivity. Using a peptide originating from a known PAD substrate myelin basic protein (RTAHYGSL, **mbpWT)** as the baseline (**Figure 6A**), we found that there was a statistically significant increase in peptide binding to MHC molecules upon citrullination. This result is consistent with the possibility that PTMs can impact presentation of peptides in the context of disease-linked peptides.

**Figure 6.**
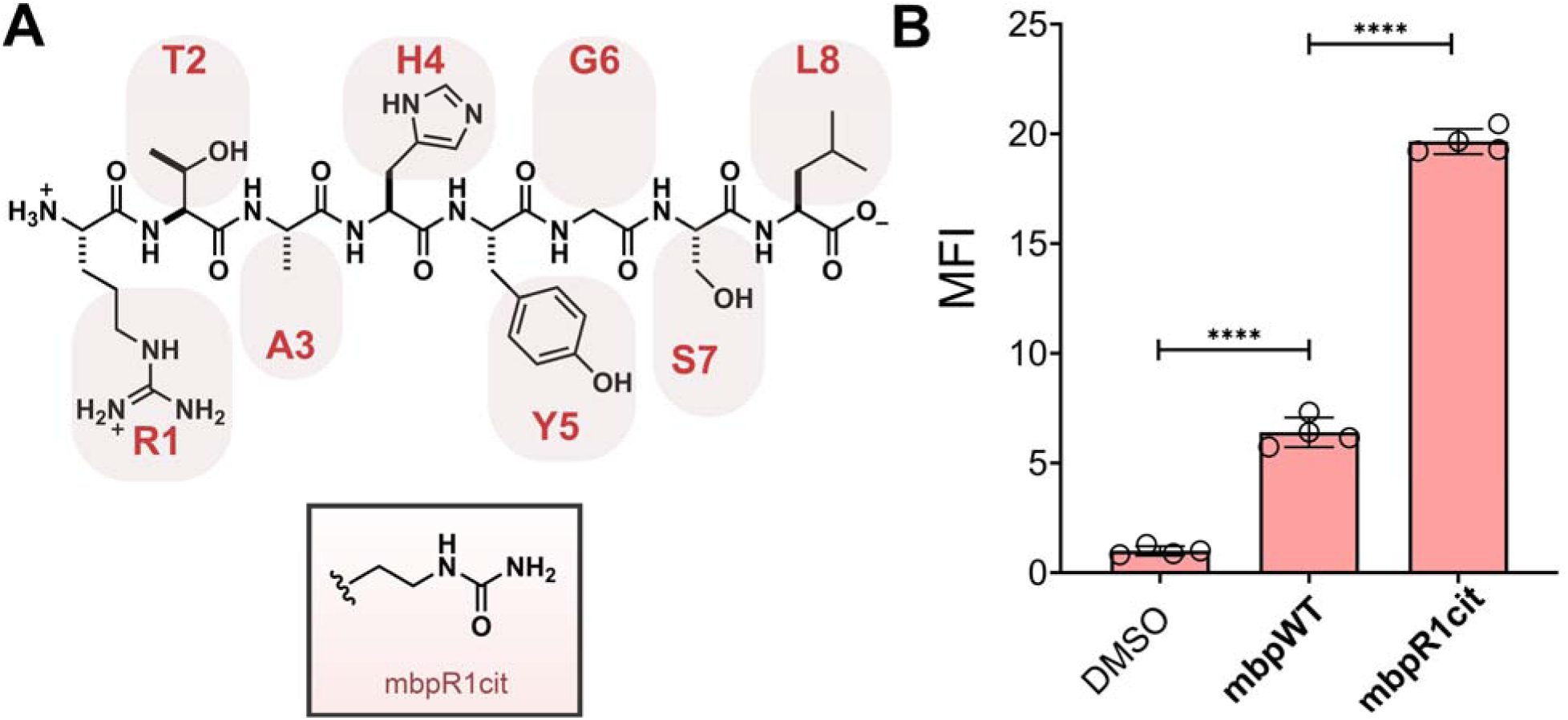
(A) Chemical structures of **mbpWT** and the PTM modified variant. (B) Flow cytometry analysis of RMA-S cells treated with specific peptide (20 μM) detected by APC conjugated anti-mouse H-2Kb antibody. Data are represented as mean ± SD (n= 3). P-values were determined by a two-tailed t-test (* p < 0.05, ** p < 0.01, *** p < 0.001, **** p < 0.0001, ns = not significant).

To gain additional insight into how PTM s might alter peptide binding affinity to H-2K^b^, we simulated peptide binding to H-2K^b^ using ROSETTA and FlexPepDock. FlexPepDock has been previously benchmarked against MHC-I bound peptides and is capable of generating models with sub-angstrom accuracy.^79, 80^ One advantage of FlexPepDock over contemporary machine learning methods is its ability to incorporate non-canonical amino acids and PTM residues by generating custom ROSETTA parameters files; this enables FlexPepDock to generate accurate models of MHC-I bound epitopes containing such residues.^81, 82^ Furthermore, the binding energy metrics calculated by this application (the reweighted_sc score term) can serve a surrogate for peptide binding affinity to MHC-I.^83^ We simulated the binding of the SARS-CoV-2-derived peptide to H-2K^b^ and generated results recapitulating experimentally derived binding affinity changes (**Figure 7**). ROSETTA correctly predicts decreased binding affinity for the top-scoring decoys after N-terminal acetylation (corresponding to an increase in the average reweighted_sc term; ROSETTA energy metrics are inversely proportional to binding affinity). ROSETTA predicts either no change or a modest increase in binding affinity for the other two PTMs assessed (**Figure 7A**). When the top-scoring model for each peptide is visually inspected, we noted significant alterations in the peptide backbone structure for the acetylated N-terminus peptide variant largely confined to the H-2K^b^ N-terminus binding pocket (**Figure 7B**), a region that is largely responsible for dictating epitope binding specificity and that cannot easily accommodate large conformational changes.^9^ Conversely, R5 citrullinated and P7 hydroxylated variants are located outside the H-2K^b^ binding pockets and regions tolerant of larger conformational changes (**Figure 7C**).

**Figure 7.**
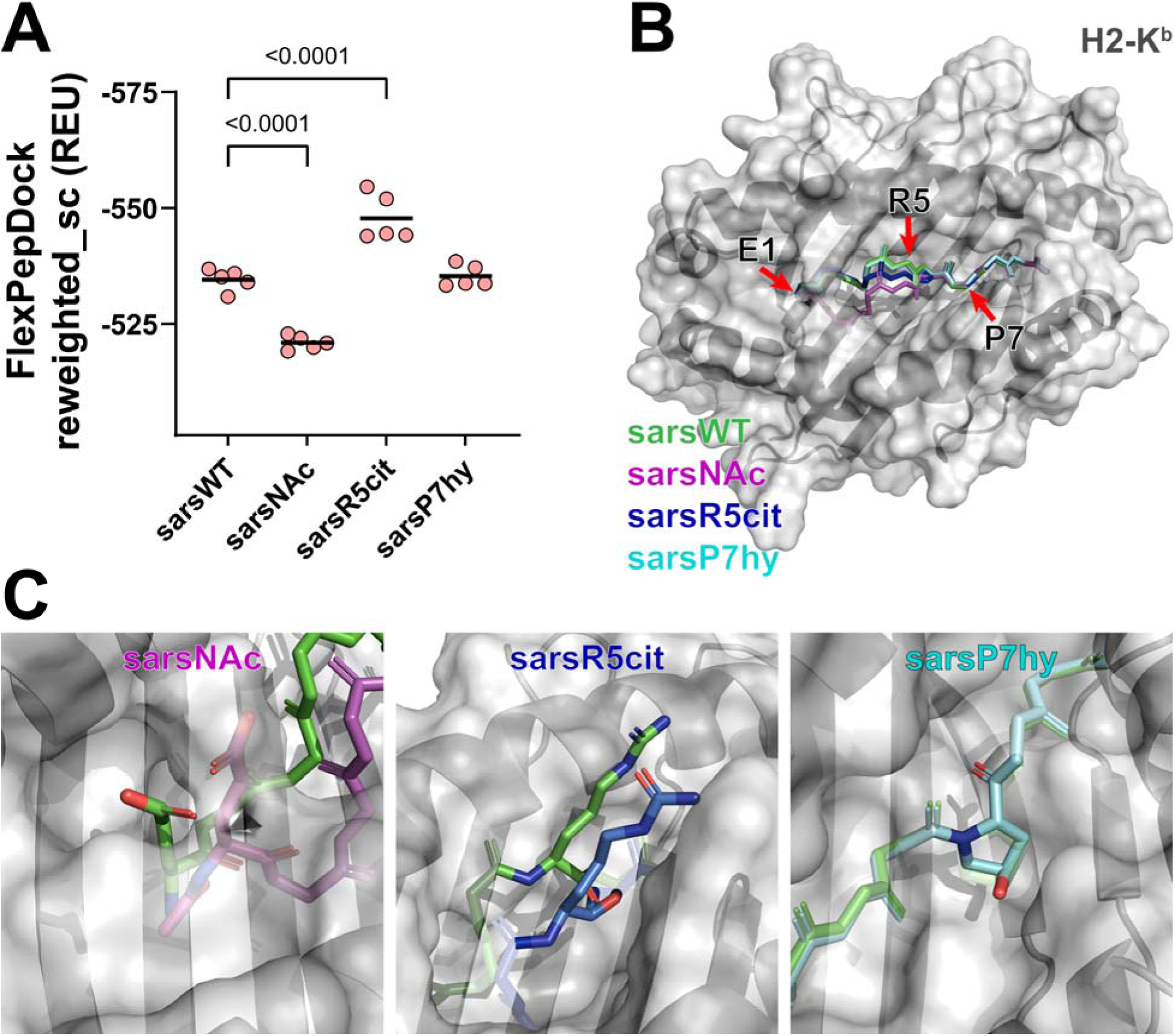
Modeling the sars peptide with and without PTMs using FlexPepDock. (A) N-terminal acetylation results in higher reweighted_sc values (and thus lower likelihood of binding) compared to the wild-type peptide, consistent with experimental results (one-way ANOVA and Holm-Sidak post-hoc; individual values correspond to the top 0.5% of models generated, and bars represent mean score term values). (B) Top peptide models generated for each epitope positioned in the H-2Kb binding cleft. (C) Side chain modifications compared to the unmodified epitope for the SARS-CoV-2 peptide.

We repeated this analysis on the ovalbumin-derived peptide and each PTM (**Figure 8A**). Phosphorylation of the N-terminus serine generates structures with higher (and therefore less favorable) score terms, again likely due to disruption of the *N*-terminus peptide-binding pocket (**Figure 8B**). Peptide models with the remaining PTMs scored either equivalently or slightly better than their wild-type counterparts. While largely consistent with experimental data, ROSETTA predicts modest increases in peptide binding favorability that are not observed experimentally. This may be a result of biases inherent in ROSETTA’s scoring weights that have not been fully optimized for unusual or non-canonical amino acids such as those modeled here. Alternatively, the assay used for assessing binding affinity may be insufficiently sensitive to detect small changes above certain affinity thresholds. Modifications at the lysine in position 7 alter T cell activation *in vitro* as illustrated in Figure 5. *In silico*, these altered residues do not interact appreciably with peptide binding pockets but instead occur in the region displayed by MHC-I to patrolling T cells (**Figure 8C**). Methylated and acetylated lysine alters residue charge and hydrophobicity; succinylation adds a negative charge (as opposed to lysine’s normally positively charged side chain); and biotinylation results in the addition of a large aliphatic heteropolycyclic side chain very different in size and character from the native lysine residue.

**Figure 8.**
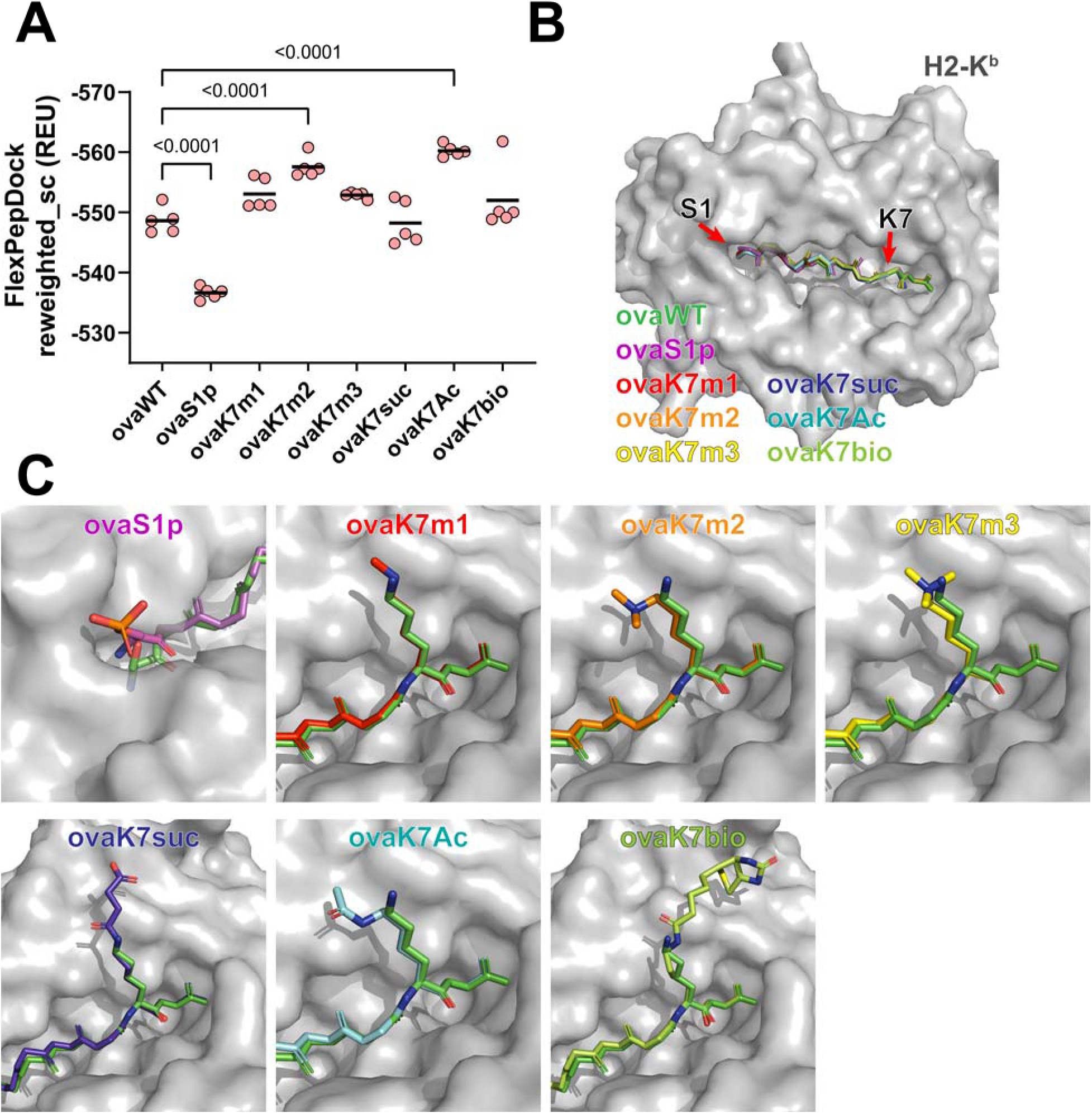
Modeling the ovalbumin-derived peptide with and without PTMs using FlexPepDock. (A) Phosphorylation of the N-terminal serine residue generates models with higher average reweighted_sc terms, similar to N-terminal acetylation of the SARS-CoV-2 peptide (one-way ANOVA and Holm-Sidak post-hoc; individual values correspond to the top 0.5% of models generated, and bars represent mean score term values). (B) Superimposed peptide backbones for the top peptide models generated for each PTM. (C) Detailed side-chain configurations for the top-scoring models generated by FlexPepDock.

Finally, we performed a similar analysis with the **mbp** peptide with and without R1 citrullination and generated *in silico* results that recapitulated experimental findings (**Figure 6**). Citrullination at R1 significantly increases the reweighted_sc metrics for the models generated by ROSETTA (**Figure S5A**). While this modification occurs at the N-terminus binding pocket, it does not appreciably alter peptide backbone configuration (**Figure S5A**, in contrast to the effects of N-terminal acetylation of the **sars** peptide). It does, however, alter side chain hydrophobicity and charge. Given that the citrullinated variant is charge-neutral (compared to arginine’s 1+ charge) and given H-2Kb’s preference for hydrophobic or charge-neutral residues at binding pockets, it is reasonable to expect this modification may enhance peptide binding affinity.

Increasingly, there are efforts dedicated to developing therapeutic agents that harness a patient’s immune system against cancerous lesions.^84–86^ A prominent example involves the modulation of the programmed death-1 (PD-1) system and its cognate programmed death-ligand 1 (PD-L1). Cancers can often hijack this set of proteins to maintain growth while suppressing the patient immune response against transformed cells. Immune checkpoint inhibitors that disrupt the association of PD-1 and PD-L1 have shown potent anti-cancer activity by a number of mechanisms but primarily *via* increased T cell engagement.^87, 88^ Neoantigens arising from genetic lesions likely provide a pool of antigens that can be presented by cancer MHCs on the cell surface during checkpoint therapy. Importantly, our results suggest that therapeutic agents that increase the presentation of peptides through an altered balance of PTM marks could potentially complement the targeting antigenic pool.

Considering the observed roles of PTMs in MHC binding and recognition by TCRs, we propose that the balance of PTM addition by “writers” and PTM removal by “erasers” can be central in immunological health and in diseased states. This imbalance could be triggered by age or manufactured by therapeutic interventions. For example, proteins could naturally exist in high abundance of non-acetylated states on lysine side chains past the peak period of negative selection of T cells. After cancer development, there is an increased level of acetylation, but those marks need to be removed by “erasers” called Lysine Deacetylases (KDAC) to reduce an anticancer immunological response against the cancerous cells. Upon the administration of KDAC inhibitors, a larger pool of acetylated peptide would be presented. As our data showed, the acetylation of lysine that interacts with TCRs would escape negative selection and provide an anticancer immunological response. Such ‘cryptic’ antigens could potentially be driving the pharmacological effect of some PTM modulators in cancer patients. To this end, there are a high number of ongoing studies evaluating the combination of KDAC inhibitors and PD-1/PDL-1 inhibitors across many types of difficult-to-treat tumors.^89–95^ We are currently working towards using our platform to demonstrate how KDAC inhibitors directly lead to an immunopeptidome that reveal cryptic antigens able to enhance anti-tumor immunological activity.

## Conclusion

In conclusion, the data presented here demonstrate that naturally occurring PTMs can drastically impact the antigen presentation pathway by altering either affinity towards MHC molecules or TCR recognition by cognate T cells. In our assays, we showed that some PTMs can disrupt peptide binding to MHC, but primarily on residues that directly engage with MHCs. Conversely, PTMs on residues whose sidechains are positioned away from the MHC binding face are more sensitive to TCR recognition due to the altered binding modality with TCRs. To the best of our knowledge, our study is the first systematic analysis of the impact that PTMs have on MHC binding in a whole cell context and how PTMs can also alter T cell activation using live target and effector cells. Provided that negative selection is age-dependent and becomes increasingly diminished past adolescence, it is evident that PTMs could be causal events that result in autoimmune diseases.

## References

1. Neefjes, J., Jongsma, M. L., Paul, P., Bakke, O. Towards a systems understanding of MHC class I and MHC class II antigen presentation. Nat Rev Immunol. 2011, 11, 823–836.

2. Roche, P. A., Furuta, K. The ins and outs of MHC class II-mediated antigen processing and presentation. Nat Rev Immunol. 2015, 15, 203–216.

3. Pishesha, N., Harmand, T. J., Ploegh, H. L. A guide to antigen processing and presentation. Nat Rev Immunol. 2022, 22, 751–764.

4. Neefjes, J., Jongsma, M. L. M., Paul, P., Bakke, O. Towards a systems understanding of MHC class I and MHC class II antigen presentation. Nat Rev Immunol. 2011, 11, 823–836.

5. Huppa, J. B., Davis, M. M. T cell-antigen recognition and the immunological synapse. Nat Rev Immunol. 2003, 3, 973–983.

6. Gorentla, B. K., Zhong, X. P. T cell Receptor Signal Transduction in T lymphocytes. J Clin Cell Immunol. 2012, 2012, 5.

7. Jameson, S. C. Maintaining the norm: T cell homeostasis. Nat Rev Immunol. 2002, 2, 547–556.

8. Klein, L., Kyewski, B., Allen, P. M., Hogquist, K. A. Positive and negative selection of the T cell repertoire: what thymocytes see (and don’t see). Nat Rev Immunol. 2014, 14, 377–391.

9. Conibear, A. C. Deciphering protein post-translational modifications using chemical biology tools. Nat Rev Chem. 2020, 4, 674–695.

10. Sandalova, T., Sala, B. M., Achour, A. Structural aspects of chemical modifications in the MHC-restricted immunopeptidome; Implications for immune recognition. Front Chem. 2022, 10.

11. Raposo, B., Merky, P., Lundqvist, C., Yamada, H., Urbonaviciute, V., Niaudet, C., Viljanen, J., Kihlberg, J., Kyewski, B., Ekwall, O., Holmdahl, R., Backlund, J. T cells specific for post-translational modifications escape intrathymic tolerance induction. Nat Commun. 2018, 9, 353.

12. Srinivasan, J., Lancaster, J. N., Singarapu, N., Hale, L. P., Ehrlich, L. I. R., Richie, E. R. Age-Related Changes in Thymic Central Tolerance. Front Immunol. 2021, 12, 676236.

13. Lancaster, J. N., Keatinge-Clay, D. E., Srinivasan, J., Li, Y., Selden, H. J., Nam, S., Richie, E. R., Ehrlich, L. I. R. Central tolerance is impaired in the middle-aged thymic environment. Aging Cell. 2022, 21, e13624.

14. Li, W., Li, F. F., Zhang, X., Lin, H. K., Xu, C. Insights into the post-translational modification and its emerging role in shaping the tumor microenvironment (vol 7, 31, 2022). Signal Transduct Tar. 2022, 7.

15. Michaelsson, E., Malmstrom, V., Reis, S., Engstrom, A., Burkhardt, H., Holmdahl, R. T cell Recognition of Carbohydrates on Type-Ii Collagen. J Exp Med. 1994, 180, 745–749.

16. Sidney, J., Vela, J. L., Friedrich, D., Kolla, R., von Herrath, M., Wesley, J. D., Sette, A. Low HLA binding of diabetes-associated CD8+T cell epitopes is increased by post translational modifications. Bmc Immunol. 2018, 19.

17. McAdam, S. N., Fleckenstein, B., Rasmussen, I. B., Schmid, D. G., Sandlie, I., Bogen, B., Viner, N. J., Sollid, L. M. T cell recognition of the dominant I-A(k)-restricted hen egg lysozyme epitope: Critical role for asparagine deamidation. J Exp Med. 2001, 193, 1239–1246.

18. Mamula, M. J., Gee, R. J., Elliott, J. I., Sette, A., Southwood, S., Jones, P. J., Blier, P. R. Isoaspartyl post-translational modification triggers autoimmune responses to self-proteins. J Biol Chem. 1999, 274, 22321–22327.

19. Kim, J. K., Mastronardi, F. G., Wood, D. D., Lubman, D. M., Zand, R., Moscarello, M. A. Multiple sclerosis: an important role for post-translational modifications of myelin basic protein in pathogenesis. Mol Cell Proteomics. 2003, 2, 453–462.

20. Raposo, B., Merky, P., Lundqvist, C., Yamada, H., Urbonaviciute, V., Niaudet, C., Viljanen, J., Kihlberg, J., Kyewski, B., Ekwall, O., Holmdahl, R., Backlund, J. T cells specific for post-translational modifications escape intrathymic tolerance induction. Nat Commun. 2018, 9.

21. Doyle, H. A., Mamula, M. J. Autoantigenesis: the evolution of protein modifications in autoimmune disease. Curr Opin Immunol. 2012, 24, 112–118.

22. Clement, C. C., Moncrieffe, H., Lele, A., Janow, G., Becerra, A., Bauli, F., Saad, F. A., Perino, G., Montagna, C., Cobelli, N., Hardin, J., Stern, L. J., Ilowite, N., Porcelli, S. A., Santambrogio, L. Autoimmune response to transthyretin in juvenile idiopathic arthritis. JCI Insight. 2016, 1.

23. Clement, C. C., Nanaware, P. P., Yamazaki, T., Negroni, M. P., Ramesh, K., Morozova, K., Thangaswamy, S., Graves, A., Kim, H. J., Li, T. W., Vigano, M., Soni, R. K., Gadina, M., Tse, H. Y., Galluzzi, L., Roche, P. A., Denzin, L. K., Stern, L. J., Santambrogio, L. Pleiotropic consequences of metabolic stress for the major histocompatibility complex class II molecule antigen processing and presentation machinery. Immunity. 2021, 54, 721–736 e710.

24. Clement, C. C., Osan, J., Buque, A., Nanaware, P. P., Chang, Y. C., Perino, G., Shetty, M., Yamazaki, T., Tsai, W. L., Urbanska, A. M., Calvo-Calle, J. M., Ramsamooj, S., Ramsamooj, S., Vergani, D., Mieli-Vergani, G., Terziroli Beretta-Piccoli, B., Gadina, M., Montagna, C., Goncalves, M. D., Sallusto, F., Galluzzi, L., Soni, R. K., Stern, L. J., Santambrogio, L. PDIA3 epitope-driven immune autoreactivity contributes to hepatic damage in type 2 diabetes. Sci Immunol. 2022, 7, eabl3795.

25. Zhai, Y., Chen, L., Zhao, Q., Zheng, Z. H., Chen, Z. N., Bian, H., Yang, X., Lu, H. Y., Lin, P., Chen, X., Chen, R., Sun, H. Y., Fan, L. N., Zhang, K., Wang, B., Sun, X. X., Feng, Z., Zhu, Y. M., Zhou, J. S., Chen, S. R., Zhang, T., Chen, S. Y., Chen, J. J., Zhang, K., Wang, Y., Chang, Y., Zhang, R., Zhang, B., Wang, L. J., Li, X. M., He, Q., Yang, X. M., Nan, G., Xie, R. H., Yang, L., Yang, J. H., Zhu, P. Cysteine carboxyethylation generates neoantigens to induce HLA-restricted autoimmunity. Science. 2023, 379, eabg2482.

26. Petersen, J., Purcell, A. W., Rossjohn, J. Post-translationally modified T cell epitopes: immune recognition and immunotherapy. J Mol Med (Berl). 2009, 87, 1045–1051.

27. Marcilla, M., Alpizar, A., Lombardia, M., Ramos-Fernandez, A., Ramos, M., Albar, J. P. Increased diversity of the HLA-B40 ligandome by the presentation of peptides phosphorylated at their main anchor residue. Mol Cell Proteomics. 2014, 13, 462–474.

28. Bassani-Sternberg, M., Braunlein, E., Klar, R., Engleitner, T., Sinitcyn, P., Audehm, S., Straub, M., Weber, J., Slotta-Huspenina, J., Specht, K., Martignoni, M. E., Werner, A., Hein, R., D, H. B., Peschel, C., Rad, R., Cox, J., Mann, M., Krackhardt, A. M. Direct identification of clinically relevant neoepitopes presented on native human melanoma tissue by mass spectrometry. Nat Commun. 2016, 7, 13404.

29. Alpizar, A., Marino, F., Ramos-Fernandez, A., Lombardia, M., Jeko, A., Pazos, F., Paradela, A., Santiago, C., Heck, A. J., Marcilla, M. A Molecular Basis for the Presentation of Phosphorylated Peptides by HLA-B Antigens. Mol Cell Proteomics. 2017, 16, 181–193.

30. Malaker, S. A., Penny, S. A., Steadman, L. G., Myers, P. T., Loke, J. C., Raghavan, M., Bai, D. L., Shabanowitz, J., Hunt, D. F., Cobbold, M. Identification of Glycopeptides as Posttranslationally Modified Neoantigens in Leukemia. Cancer Immunol Res. 2017, 5, 376–384.

31. Marino, F., Mommen, G. P., Jeko, A., Meiring, H. D., van Gaans-van den Brink, J. A., Scheltema, R. A., van Els, C. A., Heck, A. J. Arginine (Di)methylated Human Leukocyte Antigen Class I Peptides Are Favorably Presented by HLA-B*07. J Proteome Res. 2017, 16, 34–44.

32. Mohammed, F., Stones, D. H., Zarling, A. L., Willcox, C. R., Shabanowitz, J., Cummings, K. L., Hunt, D. F., Cobbold, M., Engelhard, V. H., Willcox, B. E. The antigenic identity of human class I MHC phosphopeptides is critically dependent upon phosphorylation status. Oncotarget. 2017, 8, 54160–54172.

33. Sandalova, T., Sala, B. M., Achour, A. Structural aspects of chemical modifications in the MHC-restricted immunopeptidome; Implications for immune recognition. Front Chem. 2022, 10, 861609.

34. Mangalaparthi, K. K., Madugundu, A. K., Ryan, Z. C., Garapati, K., Peterson, J. A., Dey, G., Prakash, A., Pandey, A. Digging deeper into the immunopeptidome: characterization of post-translationally modified peptides presented by MHC I. J Proteins Proteom. 2021, 12, 151–160.

35. Ireland, J., Herzog, J., Unanue, E. R. Cutting edge: Unique T cells that recognize citrullinated peptides are a feature of protein immunization. J Immunol. 2006, 177, 1421–1425.

36. Huseby, E. S., Sather, B., Huseby, P. G., Goverman, J. Age-dependent T cell tolerance and autoimmunity to myelin basic protein. Immunity. 2001, 14, 471–481.

37. Malmstrom, V., Catrina, A. I., Klareskog, L. The immunopathogenesis of seropositive rheumatoid arthritis: from triggering to targeting. Nat Rev Immunol. 2017, 17, 60–75.

38. Curran, A. M., Naik, P., Giles, J. T., Darrah, E. PAD enzymes in rheumatoid arthritis: pathogenic effectors and autoimmune targets. Nat Rev Rheumatol. 2020, 16, 301–315.

39. Yang, L., Tan, D., Piao, H. Myelin Basic Protein Citrullination in Multiple Sclerosis: A Potential Therapeutic Target for the Pathology. Neurochem Res. 2016, 41, 1845–1856.

40. Ramarathinam, S. H., Croft, N. P., Illing, P. T., Faridi, P., Purcell, A. W. Employing proteomics in the study of antigen presentation: an update. Expert Rev Proteomics. 2018, 15, 637–645.

41. Jaeger, A. M., Stopfer, L. E., Ahn, R., Sanders, E. A., Sandel, D. A., Freed-Pastor, W. A., Rideout, W. M., 3rd, Naranjo, S., Fessenden, T., Nguyen, K. B., Winter, P. S., Kohn, R. E., Westcott, P. M. K., Schenkel, J. M., Shanahan, S. L., Shalek, A. K., Spranger, S., White, F. M., Jacks, T. Deciphering the immunopeptidome in vivo reveals new tumour antigens. Nature. 2022, 607, 149–155.

42. Yi, X., Liao, Y., Wen, B., Li, K., Dou, Y., Savage, S. R., Zhang, B. caAtlas: An immunopeptidome atlas of human cancer. iScience. 2021, 24, 103107.

43. Kacen, A., Javitt, A., Kramer, M. P., Morgenstern, D., Tsaban, T., Shmueli, M. D., Teo, G. C., da Veiga Leprevost, F., Barnea, E., Yu, F., Admon, A., Eisenbach, L., Samuels, Y., Schueler-Furman, O., Levin, Y., Nesvizhskii, A. I., Merbl, Y. Post-translational modifications reshape the antigenic landscape of the MHC I immunopeptidome in tumors. Nat Biotechnol. 2023, 41, 239–251.

44. Garstka, M. A., Fish, A., Celie, P. H., Joosten, R. P., Janssen, G. M., Berlin, I., Hoppes, R., Stadnik, M., Janssen, L., Ovaa, H., van Veelen, P. A., Perrakis, A., Neefjes, J. The first step of peptide selection in antigen presentation by MHC class I molecules. Proc Natl Acad Sci U S A. 2015, 112, 1505–1510.

45. Deribe, Y. L., Pawson, T., Dikic, I. Post-translational modifications in signal integration. Nat Struct Mol Biol. 2010, 17, 666–672.

46. Zhao, Y., Jensen, O. N. Modification-specific proteomics: strategies for characterization of post-translational modifications using enrichment techniques. Proteomics. 2009, 9, 4632–4641.

47. Solleder, M., Guillaume, P., Racle, J., Michaux, J., Pak, H. S., Muller, M., Coukos, G., Bassani-Sternberg, M., Gfeller, D. Mass Spectrometry Based Immunopeptidomics Leads to Robust Predictions of Phosphorylated HLA Class I Ligands. Mol Cell Proteomics. 2020, 19, 390–404.

48. Hassan, C., Kester, M. G., Oudgenoeg, G., de Ru, A. H., Janssen, G. M., Drijfhout, J. W., Spaapen, R. M., Jimenez, C. R., Heemskerk, M. H., Falkenburg, J. H., van Veelen, P. A. Accurate quantitation of MHC-bound peptides by application of isotopically labeled peptide MHC complexes. J Proteomics. 2014, 109, 240–244.

49. Colbert, J. D., Cruz, F. M., Rock, K. L. Cross-presentation of exogenous antigens on MHC I molecules. Curr Opin Immunol. 2020, 64, 1–8.

50. Lee, J. M., Hammaren, H. M., Savitski, M. M., Baek, S. H. Control of protein stability by post-translational modifications. Nat Commun. 2023, 14, 201.

51. Ljunggren, H. G., Stam, N., Ohlen, C., Neefjes, J. J., Hoglund, P., Heemels, M. T., Bastin, J., Schumacher, T., Townsend, A., Karre, K., Ploegh, H. L. Empty Mhc Class-I Molecules Come out in the Cold. Scand J Immunol. 1990, 32, 403–403.

52. Schumacher, T. N., Heemels, M. T., Neefjes, J. J., Kast, W. M., Melief, C. J., Ploegh, H. L. Direct binding of peptide to empty MHC class I molecules on intact cells and in vitro. Cell. 1990, 62, 563–567.

53. Elvin, J., Potter, C., Elliott, T., Cerundolo, V., Townsend, A. A method to quantify binding of unlabeled peptides to class I MHC molecules and detect their allele specificity. J Immunol Methods. 1993, 158, 161–171.

54. Saito, Y., Peterson, P. A., Matsumura, M. Quantitation of peptide anchor residue contributions to class I major histocompatibility complex molecule binding. J Biol Chem. 1993, 268, 21309–21317.

55. Ross, P., Holmes, J. C., Gojanovich, G. S., Hess, P. R. A cell-based MHC stabilization assay for the detection of peptide binding to the canine classical class I molecule, DLA-88. Vet Immunol Immunopathol. 2012, 150, 206–212.

56. Apostolopoulos, V., Haurum, J. S., McKenzie, I. F. MUC1 peptide epitopes associated with five different H-2 class I molecules. Eur J Immunol. 1997, 27, 2579–2587.

57. Cerundolo, V., Elliott, T., Elvin, J., Bastin, J., Rammensee, H. G., Townsend, A. The binding affinity and dissociation rates of peptides for class I major histocompatibility complex molecules. Eur J Immunol. 1991, 21, 2069–2075.

58. De Silva, A. D., Boesteanu, A., Song, R., Nagy, N., Harhaj, E., Harding, C. V., Joyce, S. Thermolabile H-2Kb molecules expressed by transporter associated with antigen processing-deficient RMA-S cells are occupied by low-affinity peptides. J Immunol. 1999, 163, 4413–4420.

59. Feltkamp, M. C., Vierboom, M. P., Kast, W. M., Melief, C. J. Efficient MHC class I-peptide binding is required but does not ensure MHC class I-restricted immunogenicity. Mol Immunol. 1994, 31, 1391–1401.

60. Dubey, P., Hendrickson, R. C., Meredith, S. C., Siegel, C. T., Shabanowitz, J., Skipper, J. C. A., Engelhard, V. H., Hunt, D. F., Schreiber, H. The immunodominant antigen of an ultraviolet-induced regressor tumor is generated by a somatic point mutation in the DEAD box helicase p68. J Exp Med. 1997, 185, 695–705.

61. Kjellen, P., Brunsberg, U., Broddefalk, J., Hansen, B., Vestberg, M., Ivarsson, I., Engstrom, A., Svejgaard, A., Kihlberg, J., Fugger, L., Holmdahl, R. The structural basis of MHC control of collagen-induced arthritis; binding of the immunodominant type II collagen 256-270 glycopeptide to H-2Aq and H-2Ap molecules. Eur J Immunol. 1998, 28, 755–767.

62. Rosloniec, E. F., Whittington, K. B., Zaller, D. M., Kang, A. H. HLA-DR1 (DRB1*0101) and DR4 (DRB1*0401) use the same anchor residues for binding an immunodominant peptide derived from human type II collagen. J Immunol. 2002, 168, 253–259.

63. Huizinga, T. W., Amos, C. I., van der Helm-van Mil, A. H., Chen, W., van Gaalen, F. A., Jawaheer, D., Schreuder, G. M., Wener, M., Breedveld, F. C., Ahmad, N., Lum, R. F., de Vries, R. R., Gregersen, P. K., Toes, R. E., Criswell, L. A. Refining the complex rheumatoid arthritis phenotype based on specificity of the HLA-DRB1 shared epitope for antibodies to citrullinated proteins. Arthritis Rheum. 2005, 52, 3433–3438.

64. Yague, J., Alvarez, I., Rognan, D., Ramos, M., Vazquez, J., de Castro, J. A. An N-acetylated natural ligand of human histocompatibility leukocyte antigen (HLA)-B39. Classical major histocompatibility complex class I proteins bind peptides with a blocked NH(2) terminus in vivo. J Exp Med. 2000, 191, 2083–2092.

65. Sun, M., Liu, J., Qi, J., Tefsen, B., Shi, Y., Yan, J., Gao, G. F. Nalpha-terminal acetylation for T cell recognition: molecular basis of MHC class I-restricted nalpha-acetylpeptide presentation. J Immunol. 2014, 192, 5509–5519.

66. de Haan, E. C., Wauben, M. H., Wagenaar-Hilbers, J. P., Grosfeld-Stulemeyer, M. C., Rijkers, D. T., Moret, E. E., Liskamp, R. M. Stabilization of peptide guinea pig myelin basic protein 72-85 by N-terminal acetylation-implications for immunological studies. Mol Immunol. 2004, 40, 943–948.

67. Shastri, N., Gonzalez, F. Endogenous generation and presentation of the ovalbumin peptide/Kb complex to T cells. J Immunol. 1993, 150, 2724–2736.

68. Koch, C. P., Perna, A. M., Pillong, M., Todoroff, N. K., Wrede, P., Folkers, G., Hiss, J. A., Schneider, G. Scrutinizing MHC-I binding peptides and their limits of variation. PLoS Comput Biol. 2013, 9, e1003088.

69. van der Gracht, A. M. F., de Geus, M. A. R., Camps, M. G. M., Ruckwardt, T. J., Sarris, A. J. C., Bremmers, J., Maurits, E., Pawlak, J. B., Posthoorn, M. M., Bonger, K. M., Filippov, D. V., Overkleeft, H. S., Robillard, M. S., Ossendorp, F., van Kasteren, S. I. Chemical Control over T cell Activation in Vivo Using Deprotection of trans-Cyclooctene-Modified Epitopes. ACS Chem Biol. 2018, 13, 1569–1576.

70. Fremont, D. H., Stura, E. A., Matsumura, M., Peterson, P. A., Wilson, I. A. Crystal structure of an H-2Kb-ovalbumin peptide complex reveals the interplay of primary and secondary anchor positions in the major histocompatibility complex binding groove. Proc Natl Acad Sci U S A. 1995, 92, 2479–2483.

71. Swee, L. K., Guimaraes, C. P., Sehrawat, S., Spooner, E., Barrasa, M. I., Ploegh, H. L. Sortase-mediated modification of alphaDEC205 affords optimization of antigen presentation and immunization against a set of viral epitopes. Proc Natl Acad Sci U S A. 2013, 110, 1428–1433.

72. Fremont, D. H., Stura, E. A., Matsumura, M., Peterson, P. A., Wilson, I. A. Crystal-Structure of an H-2k(B)-Ovalbumin Peptide Complex Reveals the Interplay of Primary and Secondary Anchor Positions in the Major Histocompatibility Complex Binding Groove. P Natl Acad Sci USA. 1995, 92, 2479–2483.

73. Deres, K., Beck, W., Faath, S., Jung, G., Rammensee, H. G. MHC/peptide binding studies indicate hierarchy of anchor residues. Cell Immunol. 1993, 151, 158–167.

74. Bremel, R. D., Homan, E. J. Frequency Patterns of T cell Exposed Amino Acid Motifs in Immunoglobulin Heavy Chain Peptides Presented by MHCs. Front Immunol. 2014, 5, 541.

75. Reddehase, M. J., Rothbard, J. B., Koszinowski, U. H. A pentapeptide as minimal antigenic determinant for MHC class I-restricted T lymphocytes. Nature. 1989, 337, 651–653.

76. Karttunen, J., Shastri, N. Measurement of ligand-induced activation in single viable T cells using the lacZ reporter gene. Proc Natl Acad Sci U S A. 1991, 88, 3972–3976.

77. Curran, A. M., Girgis, A. A., Jang, Y., Crawford, J. D., Thomas, M. A., Kawalerski, R., Coller, J., Bingham, C. O., 3rd, Na, C. H., Darrah, E. Citrullination modulates antigen processing and presentation by revealing cryptic epitopes in rheumatoid arthritis. Nat Commun. 2023, 14, 1061.

78. Martinod, K., Witsch, T., Erpenbeck, L., Savchenko, A., Hayashi, H., Cherpokova, D., Gallant, M., Mauler, M., Cifuni, S. M., Wagner, D. D. Peptidylarginine deiminase 4 promotes age-related organ fibrosis. J Exp Med. 2017, 214, 439–458.

79. Liu, T., Pan, X., Chao, L., Tan, W., Qu, S., Yang, L., Wang, B., Mei, H. Subangstrom accuracy in pHLA-I modeling by Rosetta FlexPepDock refinement protocol. J Chem Inf Model. 2014, 54, 2233–2242.

80. Raveh, B., London, N., Schueler-Furman, O. Sub-angstrom modeling of complexes between flexible peptides and globular proteins. Proteins. 2010, 78, 2029–2040.

81. Bloodworth, N., Barbaro, N. R., Moretti, R., Harrison, D. G., Meiler, J. Rosetta FlexPepDock to predict peptide-MHC binding: An approach for non-canonical amino acids. PLoS One. 2022, 17, e0275759.

82. Tivon, B., Gabizon, R., Somsen, B. A., Cossar, P. J., Ottmann, C., London, N. Covalent flexible peptide docking in Rosetta. Chem Sci. 2021, 12, 10836–10847.

83. Alam, N., Schueler-Furman, O. Modeling Peptide-Protein Structure and Binding Using Monte Carlo Sampling Approaches: Rosetta FlexPepDock and FlexPepBind. Methods Mol Biol. 2017, 1561, 139–169.

84. Marin-Acevedo, J. A., Dholaria, B., Soyano, A. E., Knutson, K. L., Chumsri, S., Lou, Y. Next generation of immune checkpoint therapy in cancer: new developments and challenges. J Hematol Oncol. 2018, 11, 39.

85. Topalian, S. L., Hodi, F. S., Brahmer, J. R., Gettinger, S. N., Smith, D. C., McDermott, D. F., Powderly, J. D., Carvajal, R. D., Sosman, J. A., Atkins, M. B., Leming, P. D., Spigel, D. R., Antonia, S. J., Horn, L., Drake, C. G., Pardoll, D. M., Chen, L., Sharfman, W. H., Anders, R. A., Taube, J. M., McMiller, T. L., Xu, H., Korman, A. J., Jure-Kunkel, M., Agrawal, S., McDonald, D., Kollia, G. D., Gupta, A., Wigginton, J. M., Sznol, M. Safety, activity, and immune correlates of anti-PD-1 antibody in cancer. N Engl J Med. 2012, 366, 2443–2454.

86. Tumeh, P. C., Harview, C. L., Yearley, J. H., Shintaku, I. P., Taylor, E. J., Robert, L., Chmielowski, B., Spasic, M., Henry, G., Ciobanu, V., West, A. N., Carmona, M., Kivork, C., Seja, E., Cherry, G., Gutierrez, A. J., Grogan, T. R., Mateus, C., Tomasic, G., Glaspy, J. A., Emerson, R. O., Robins, H., Pierce, R. H., Elashoff, D. A., Robert, C., Ribas, A. PD-1 blockade induces responses by inhibiting adaptive immune resistance. Nature. 2014, 515, 568–571.

87. Riaz, N., Morris, L., Havel, J. J., Makarov, V., Desrichard, A., Chan, T. A. The role of neoantigens in response to immune checkpoint blockade. Int Immunol. 2016, 28, 411–419.

88. Gubin, M. M., Zhang, X., Schuster, H., Caron, E., Ward, J. P., Noguchi, T., Ivanova, Y., Hundal, J., Arthur, C. D., Krebber, W. J., Mulder, G. E., Toebes, M., Vesely, M. D., Lam, S. S., Korman, A. J., Allison, J. P., Freeman, G. J., Sharpe, A. H., Pearce, E. L., Schumacher, T. N., Aebersold, R., Rammensee, H. G., Melief, C. J., Mardis, E. R., Gillanders, W. E., Artyomov, M. N., Schreiber, R. D. Checkpoint blockade cancer immunotherapy targets tumour-specific mutant antigens. Nature. 2014, 515, 577–581.

89. Woods, D. M., Sodre, A. L., Villagra, A., Sarnaik, A., Sotomayor, E. M., Weber, J. HDAC Inhibition Upregulates PD-1 Ligands in Melanoma and Augments Immunotherapy with PD-1 Blockade. Cancer Immunol Res. 2015, 3, 1375–1385.

90. Gameiro, S. R., Malamas, A. S., Tsang, K. Y., Ferrone, S., Hodge, J. W. Inhibitors of histone deacetylase 1 reverse the immune evasion phenotype to enhance T cell mediated lysis of prostate and breast carcinoma cells. Oncotarget. 2016, 7, 7390–7402.

91. Llopiz, D., Ruiz, M., Villanueva, L., Iglesias, T., Silva, L., Egea, J., Lasarte, J. J., Pivette, P., Trochon-Joseph, V., Vasseur, B., Dixon, G., Sangro, B., Sarobe, P. Enhanced anti-tumor efficacy of checkpoint inhibitors in combination with the histone deacetylase inhibitor Belinostat in a murine hepatocellular carcinoma model. Cancer Immunol Immunother. 2019, 68, 379–393.

92. Burke, B., Eden, C., Perez, C., Belshoff, A., Hart, S., Plaza-Rojas, L., Delos Reyes, M., Prajapati, K., Voelkel-Johnson, C., Henry, E., Gupta, G., Guevara-Patino, J. Inhibition of Histone Deacetylase (HDAC) Enhances Checkpoint Blockade Efficacy by Rendering Bladder Cancer Cells Visible for T Cell-Mediated Destruction. Front Oncol. 2020, 10, 699.

93. Baretti, M., Yarchoan, M. Epigenetic modifiers synergize with immune-checkpoint blockade to enhance long-lasting antitumor efficacy. J Clin Invest. 2021, 131.

94. Truong, A. S., Zhou, M., Krishnan, B., Utsumi, T., Manocha, U., Stewart, K. G., Beck, W., Rose, T. L., Milowsky, M. I., He, X., Smith, C. C., Bixby, L. M., Perou, C. M., Wobker, S. E., Bailey, S. T., Vincent, B. G., Kim, W. Y. Entinostat induces antitumor immune responses through immune editing of tumor neoantigens. J Clin Invest. 2021, 131.

95. Zhang, P., Du, Y., Bai, H., Wang, Z., Duan, J., Wang, X., Zhong, J., Wan, R., Xu, J., He, X., Wang, D., Fei, K., Yu, R., Tian, J., Wang, J. Optimized dose selective HDAC inhibitor tucidinostat overcomes anti-PD-L1 antibody resistance in experimental solid tumors. BMC Med. 2022, 20, 435.

