## Supporting Information for "Impact of Post-Translational Modification on MHC Peptide Binding and TCR Engagement"

### Table of Contents


| | |
|---|---|
| **Supporting Figures** | **S3-S6** |
| Figure S1: RMA-S Stabilization Assay Workflow | S3 |
| Figure S2: RMA-S Stabilization Assay sarsWT Concentration Scan | S4 |
| Figure S3: RMA-S Stabilization Assay 37 °C Time Scan | S5 |
| Figure S4: DC2.4 T Cell Activation | S6 |
| **Materials and Methods** | **S7** |
| Materials | S7 |
| Mammalian cell culture | S7 |
| RMA-S Stabilization Assay | S7 |
| B3Z T cell activation | S7 |
| **Synthesis and Characterization** | **S8-S33** |
| Scheme S1: SIINFEKL | S8 |
| Scheme S2: Monomethyl Lysine SIINFKEL | S10 |
| Scheme S3: Dimethyl Lysine SIINFKEL | S12 |
| Scheme S4: Trimethyl Lysine SIINFKEL | S14 |
| Scheme S5: Succinyl Lysine SIINFKEL | S16 |
| Scheme S6. Acetyl Lysine SIINFKEL | S18 |
| Scheme S7: Biotinylated Lysine SIINFKEL | S20 |
| Scheme S8: Phosphoserine SIINFEKL | S22 |
| Scheme S8: SNFVSAGI | S24 |
| Scheme S9: ESIVRFPNI | S26 |
| Scheme S10: N-Acetyl ESIVRFPNI | S28 |
| Scheme S11: Citrullinated ESIVRFPNI | S30 |
| Scheme S12: Hydroxy Proline ESIVRFPNI | S32 |

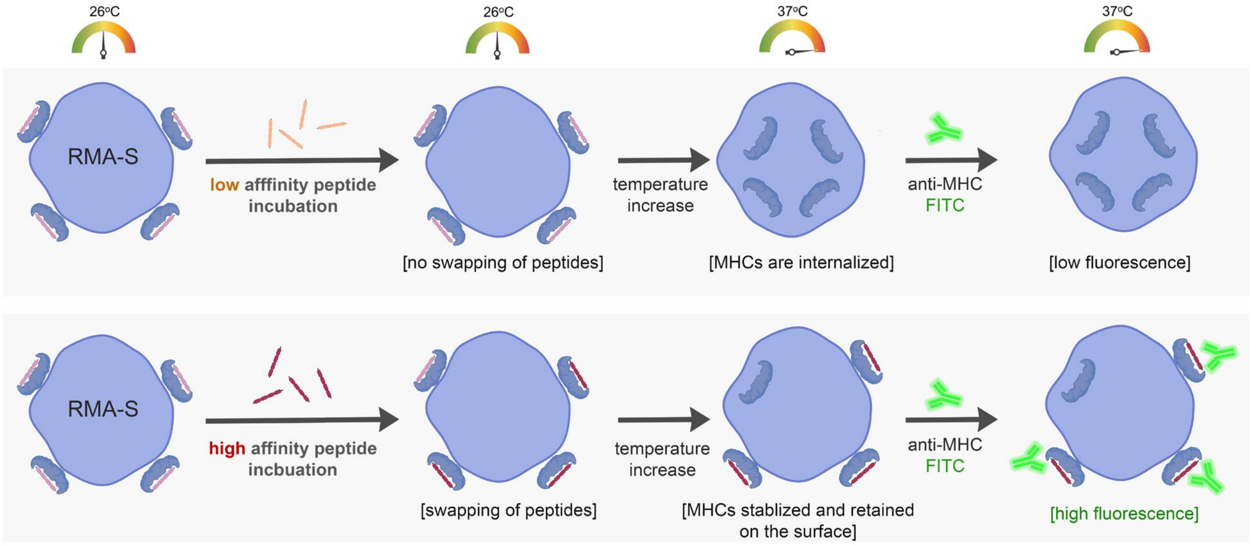


**Figure S1.**RMA-S stabilization assay workflow. RMA-S cells express low affinity peptides on its surface at 26 ºC. When the temperature is raised to 37 ºC, in the absence of a high affinity MHC binding peptide (top) the low affinity pMHC complex dissociates and the empty MHC is internalized and degraded. In the presence of a high affinity binder (bottom) the pMHC complex remains stable at 37 ºC and can be detected via fluorescently labeled anti-MHC antibodies.

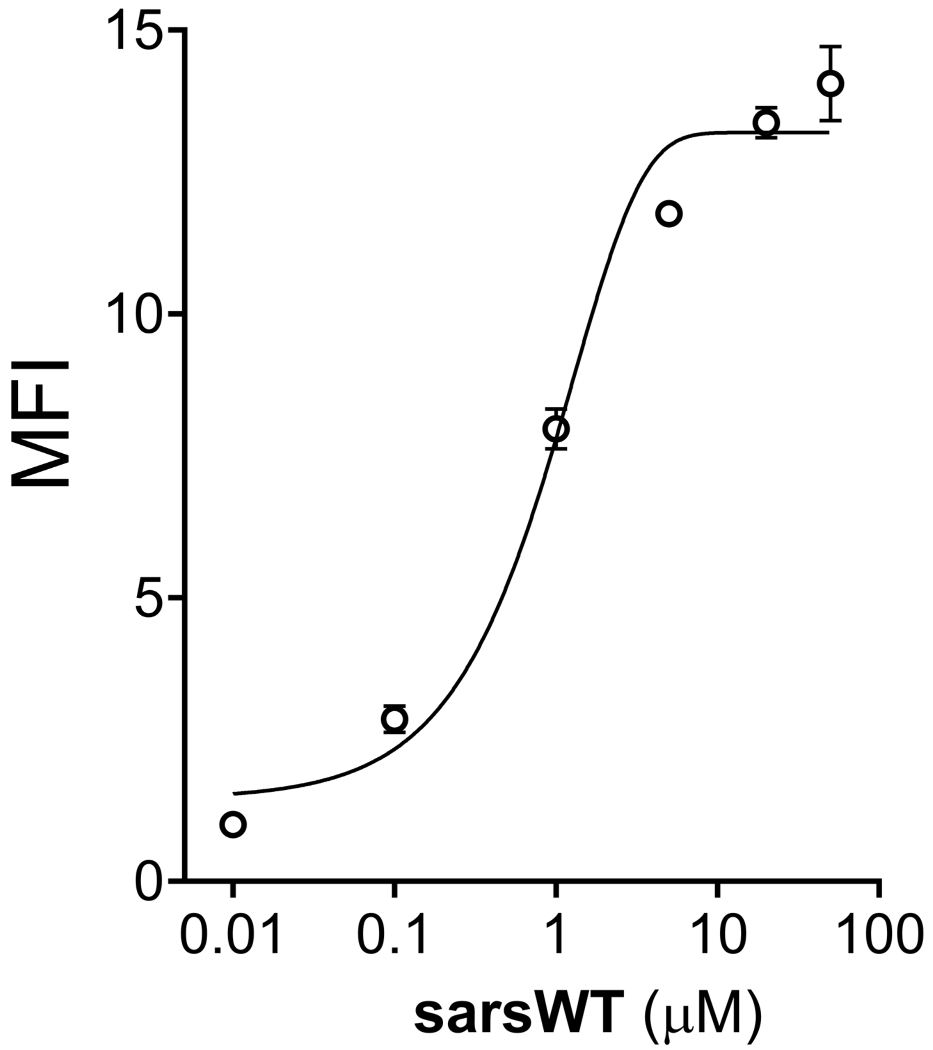


**Figure S2.** Flow cytometry analysis of RMA-S cells treated with indicated concentration of **sarsWT** detected by APC conjugated anti-mouse H-2K$^b$ antibody. Data are represented as mean ± SD (n= 3). P-values were determined by a two-tailed *t*-test (* p < 0.05, ** p < 0.01, *** p < 0.001, **** p < 0.0001, ns = not significant).

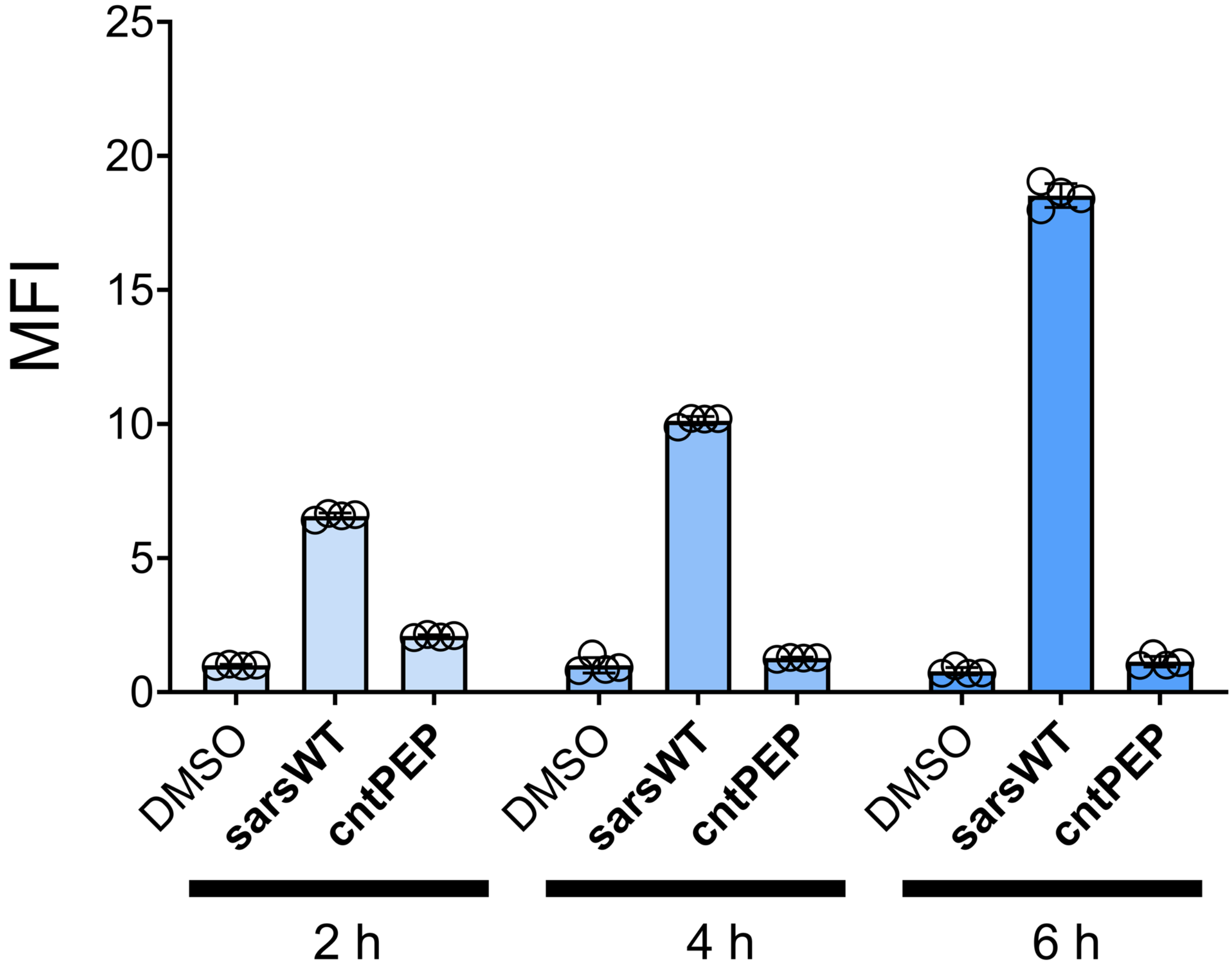


**Figure S3.** Flow cytometry analysis of RMA-S cells treated with of **sarsWT** (20 μM) for indicated time points detected by APC conjugated anti-mouse H-$2K^b$ antibody. Data are represented as mean ± SD (n= 3). P-values were determined by a two-tailed *t*-test (* $p < 0.05$, ** $p < 0.01$, *** $p < 0.001$, **** $p < 0.0001$, ns = not significant).

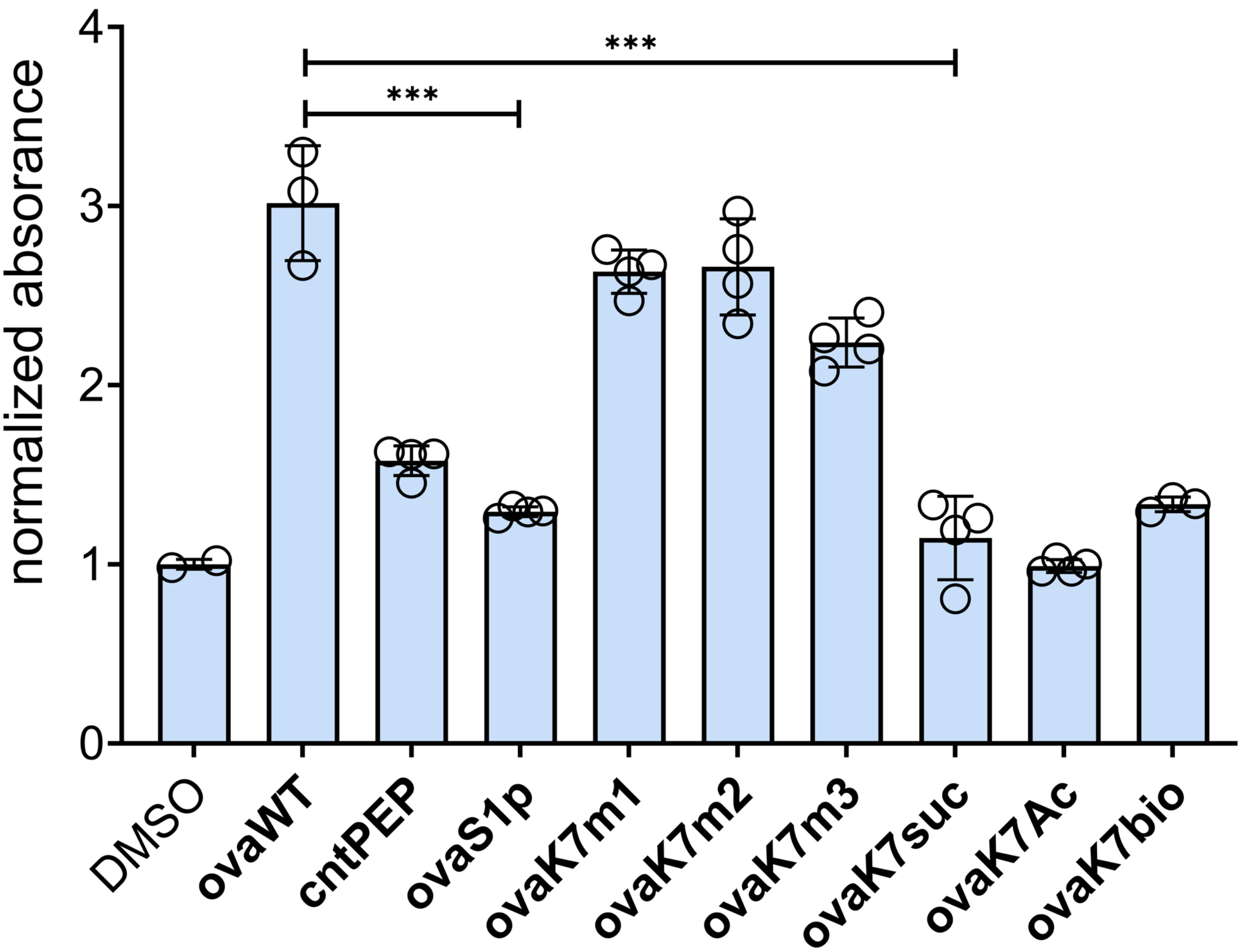


**Figures S4.** DC2.4 cells were incubated with peptide and B3Z T-cells overnight at an effector to target ratio of 1:1. β-galactosidase expression was then measured via the colorimetric reagent CPRG on a plate reader at 570 nm.

**Figure 5.** Modeling the peptide mbp with and without citrullination of the N-terminus arginine residue. (A) Arginine citrullination generates models with a higher average reweighted_sc, consistent with experimental results showing citrullination enhances binding affinity (two-tailed student's t test; individual values correspond to the top 0.5% of models generated, and bars represent mean score term values). (B) Superimposed backbones for the top model generated for the mbpWT and mbpR1cit variants show little difference in overall peptide configuration. (C) Detailed side-chain configuration for both the mbpWT and mbpR1cit variant at the N-terminal arginine residue.

**Materials.**

All peptide related reagents and protected amino acids were purchased from Chem-Impex. APC-labeled anti-mouse H-2K$^d$/H-2D$^d$ antibody was purchased from Biolegend. Pooled Human Serum was purchase from Sigma Aldrich. Dulbecco's Modified Eagle's Medium (DMEM) was purchased from VWR. Fetal Bovine Serum (FBS) was purchased from R&D Systems. Penicillin-Streptomycin was purchased from Sigma-Aldrich. All other organic chemical reagents were purchased from Fisher Scientific or Sigma Aldrich and used without further purification. ALL COMPOUNDS ARE >95% PURE BY HPLC ANALYSIS.

#### Experimental Methods.

**Mammalian Cell Culture.** RMA-S cells were a kind gift from Dr. John Sampson. RMA-S cells were maintained in RPMI 1640 media supplemented with 10% fetal bovine serum, 50 IU/mL penicillin, 50 ug/mL streptomycin,1X a MEM non-essential amino acid solution (ThermoFisher) and cultured in a humidified atmosphere of 5% $CO_2$ at 37°C. B3Z cells were a kindly provided by Dr. Aaron Esser-Kahn and maintained in RPMI 1640 media supplemented with 10% fetal bovine serum, 50 IU/mL penicillin, 50 ug/mL streptomycin and cultured in a humidified atmosphere of 5% $CO_2$ at 37°C.

**RMA-S Stabilization Assay**. $10^5$ RMA-S cells were seeded in a treated 96 well plate at 37°C overnight. The next day, RMA-S cells were moved to a 26°C incubator for 24-48 hours. Following the incubation period, cells were incubated with peptide in culture media at indicated concentrations for 1 hour at 26°C before being moved to the 37°C incubator for 6 hours. The media was then replaced with a 1:100 dilution of APC-labeled anti-mouse H-2K$^d$/H-2D$^d$ antibody in culture media for 1 hour at 4°C. Cells were removed from the well plate by vigorous pipetting, fixed with 2% formaldehyde solution, and analyzed using the Attune NxT Flow Cytometer (Thermo Fischer) equipped with a 637 nm laser with 670/14 nm bandpass filter.

**B3Z T cell activation**: $10^5$ RMA-S cells were seeded in a treated 96 well plate at 37°C overnight. The following day the culture media was replaced with media containing indicated concentration of peptide along with $10^5$ B3Z cells in culture media and co-incubated overnight. Cells were then spun down at 500xg for 5 mins and washed with 1X PBS a total of two times. Lysis buffer containing 0.02% saponin, 500 μM CPRG reagent, 100 mM MgCl2, and 100 mM beta mercaptoethanol in 1X PBS was added to each well. After 2-4 hours absorbance 570 was recorded using [insert plate reader]

#### Computational Methods

We used the FlexPepDock refinement application in combination with ROSETTA scripts to simulate peptide docking to H-2K$^b$ in ROSETTA 3.13.[1,2] FlexPepDock refinement has successfully recapitulated peptide/MHC-I complex structures with sub-angstrom accuracy,[3] and we have used this protocol to generate peptide/MHC-I structures for peptides containing post-translational modifications and other non-canonical amino acids.[4] The refinement protocol requires initial templates that approximate the final peptide configuration. We identified templates by scoring sequence alignments between the peptide to model and H2-K$^b$ bound peptides with structures available in the PDB. The H-2K$^b$ MHC-1 model was generated using AlphFold2.[5] Residues in the template peptide were sequentially mutated to match those of the peptide to model using the ROSETTA mover MutateResidue. For modeling residues with post-translational modifications, we applied either pre-existing patches with the ROSETTA mover ModifyVariantType or created *de-novo* parameter files and rotamer libraries as described (see supplemental table SX).[6] For each docking simulation 1000 models were created. Models were sorted by the reweighted_sc statistic (a modified ROSETTA energy score term that doubles contributions from interface residues and triples contributions from peptide residues) and the top 0.5% scoring models selected for further analysis. We previously validated this statistic as a metric for peptide binding affinity using a variant of the FlexPepBind protocol as described by Alam et al.[4,7] Computations were carried out with resources provided by the Vanderbilt Advanced Computing Center for Research and Education (ACCRE). Top-scoring models were rendered and visually inspected using the Pymol Molecular Graphics System v2.0 by Schrödinger, LLC.

**Scheme S1. Synthesis of SIINFKEL**

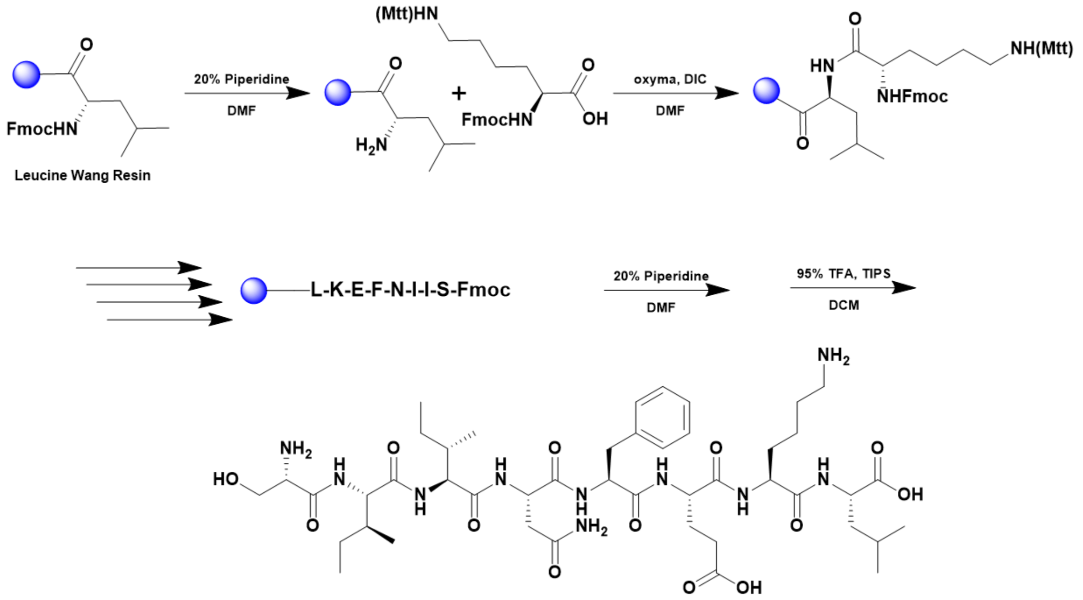


A 25 mL vessel of CEM discover bio manual peptide synthesizer was charged with 0.25 mmol of leucine wang resin. The Fmoc group was removed by using a 20% piperidine solution in DMF (10 mL). Using Synergy software, the deprotection protocol was run. The piperidine solution was drained and the resin was washed with DMF (4 x 10 mL). Fmoc-L-lysine(Mtt)-OH (5 eq, 1.25 mM) along with Oxyma (5 eq, 1.25 mM) and DIC (5 eq, 1.35 mmol) in DMF was added to the reaction vessel and the coupling protocol was run. The amino acid solution was drained, and the resin was washed with DMF (2 x 10 mL). The fmoc removal and coupling procedure was repeated as before using the same equivalencies for the remaining amino acids. To remove the peptide from resin, a TFA cocktail solution (95% TFA, 2.5% TIPS, and 2.5% DCM) was added to the resin and agitated for 2 hours. The resin was filtered, and the resulting solution was concentrated in vacuo. The peptide was trituated with cold diethyl ether and purified using reverse phase HPLC using $H_2O/CH_3CN$. The sample was analyzed for purity using a Waters 1525 Binary HPLC Pump using a Phenomenex Luna 5u C8(2) 100A (250 x 4.60 mm) column; gradient eluted with $H_20/CH_3CN$. Molecular weight was confirmed using high resolution electrospray ionization mass spectrometry (HRMS, ESI/MS) analyses obtained on an Agilent 6545B Q-TOF LC/MS equipped with 1260 infinity II LC system with auto sampler. The final peptide product was lyophilized and stored at -20°C until further use.

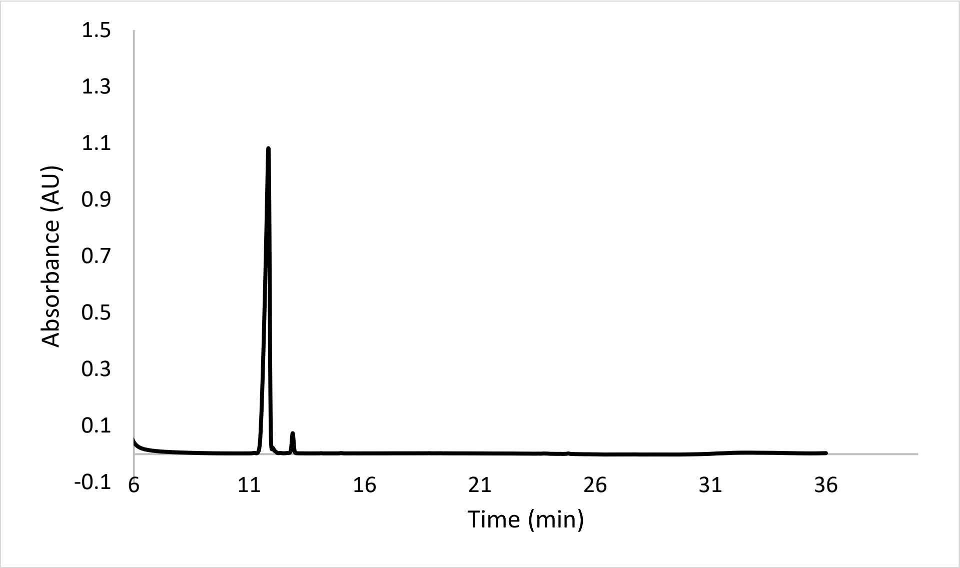


ESI-MS calculated [M+H$^+$]: 963.5515, found 963.5503.

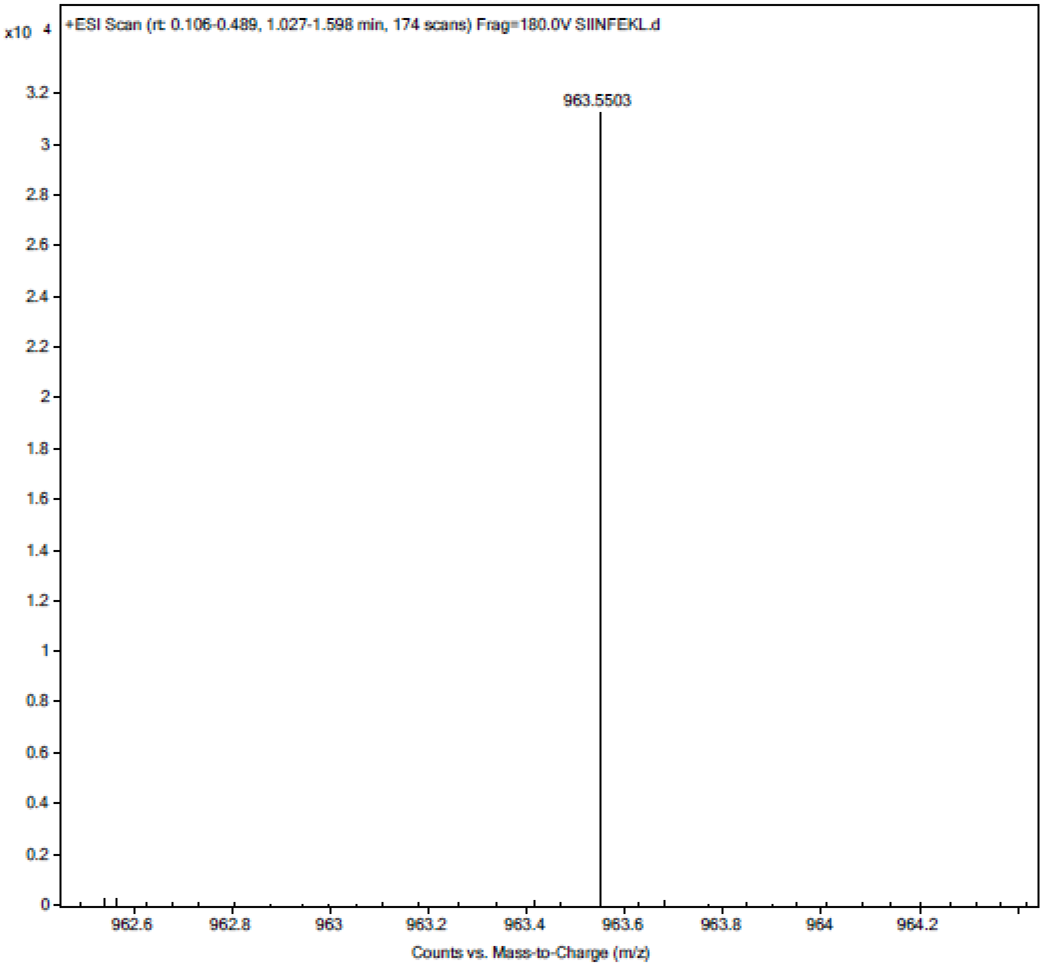


**Scheme S2. Synthesis of Monomethyl Lysine SIINFKEL**

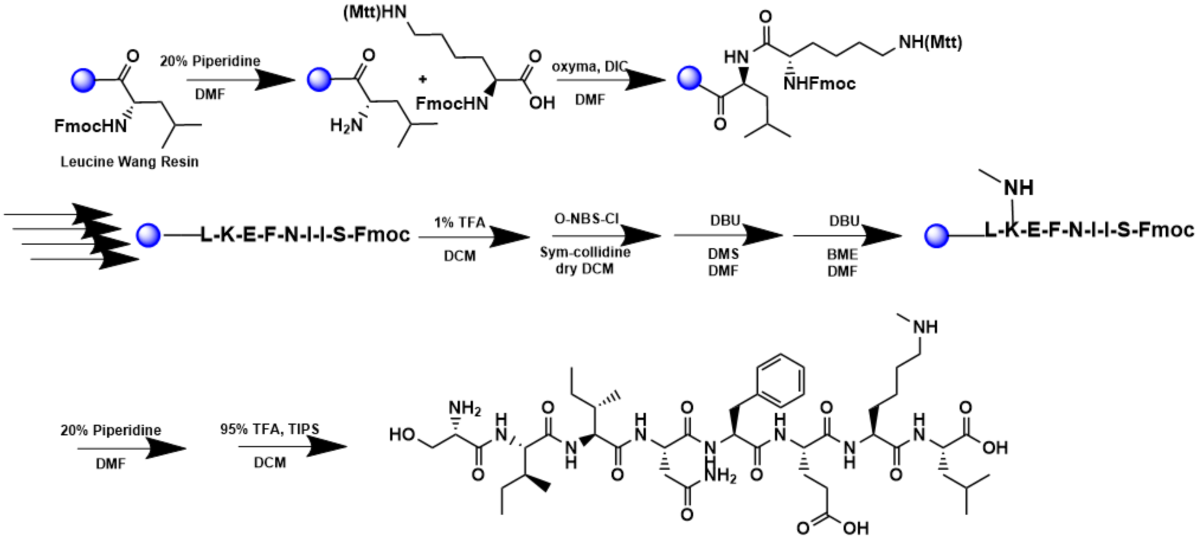


A 25 mL vessel of CEM discover bio manual peptide synthesizer was charged with 0.25 mmol of leucine wang resin. The Fmoc group was removed by using a 20% piperidine solution in DMF (10 mL). Using Synergy software, the deprotection protocol was run. The piperidine solution was drained and the resin was washed with DMF (4 x 10 mL). Fmoc-L-lysine(Mtt)-OH (5 eq, 1.25 mM) along with Oxyma (5 eq, 1.25 mM) and DIC (5 eq, 1.35 mmol) in DMF was added to the reaction vessel and the coupling protocol was run. The amino acid solution was drained, and the resin was washed with DMF (2 x 10 mL). The fmoc removal and coupling procedure was repeated as before using the same equivalencies for the remaining amino acids. The MTT protecting group was removed by the addition of 1% TFA, 2.5% TIPS, in 10 mL DCM for 15 min, washed and repeated 5 more times. O-NBS-Cl (4 eq, 1 mM) was added along with sym-collidine (10 eq, 2.5 mM) in dry DCM and treated with resin for 15 mins while agitated for a total of 2x. The solution was drained and DBU (3 eq, 0.75 mM) and DMS (10 eq, 2.5 mM) in DMF was added and agitated for 5 mins. The solution was drained and BME (10 eq, 2.5 mM) and DBU (5 eq, 1.25 mM) we added in DMF, and the solution was agitated for 30 mins.To remove the peptide from resin, a TFA cocktail solution (95% TFA, 2.5% TIPS, and 2.5% DCM) was added to the resin and agitated for 2 hours. The resin was filtered, and the resulting solution was concentrated in vacuo. The peptide was trituated with cold diethyl ether and purified using reverse phase HPLC using $H_2O/CH_3CN$. The sample was analyzed for purity using a Waters 1525 Binary HPLC Pump using a Phenomenex Luna 5u C8(2) 100A (250 x 4.60 mm) column; gradient eluted with $H_20/CH_3CN$. Molecular weight was confirmed using high resolution electrospray ionization mass spectrometry (HRMS, ESI/MS) analyses obtained on an Agilent 6545B Q-TOF LC/MS equipped with 1260 infinity II LC system with auto sampler. The final peptide product was lyophilized and stored at -20$^{o}$C until further use.

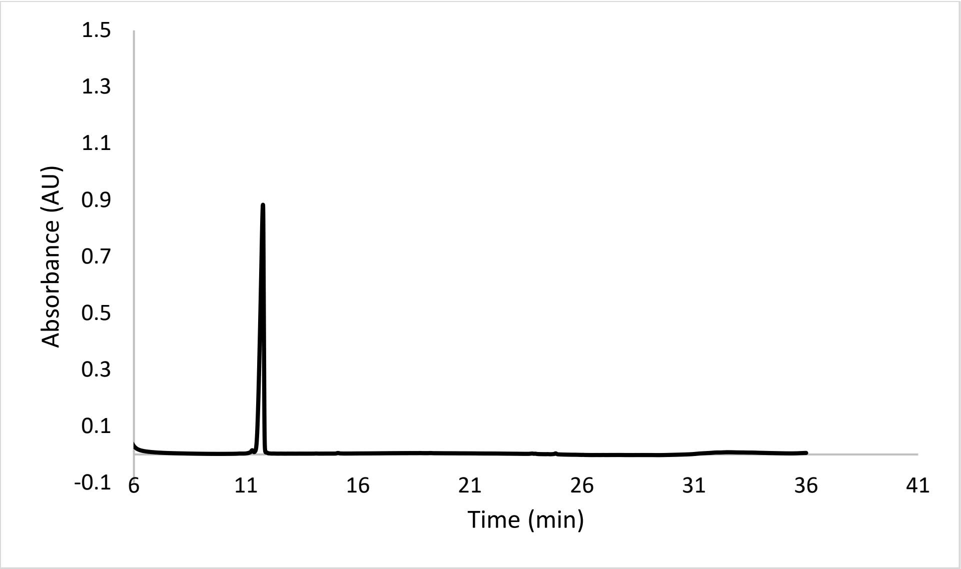


ESI-MS calculated [M+H$^+$]: 977.5671, found 977.5666.

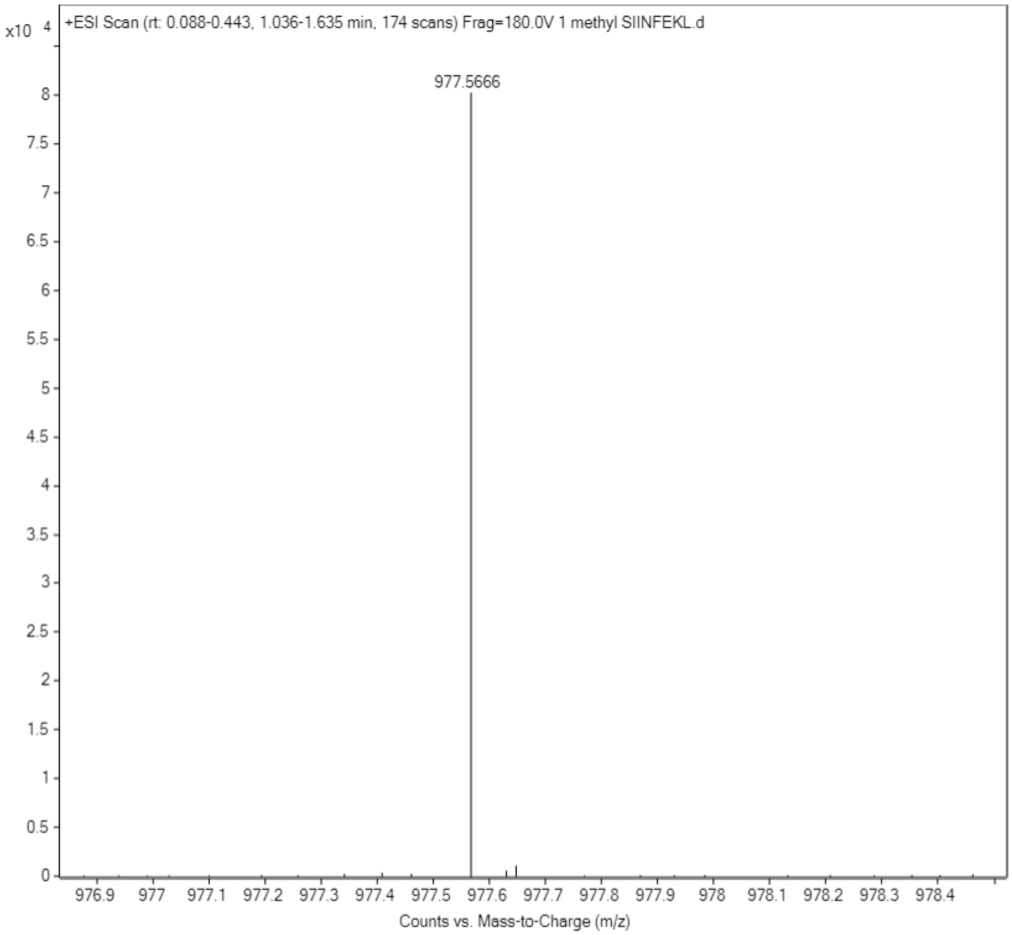

**Scheme S3. Synthesis of Dimethyl Lysine SIINFKEL**

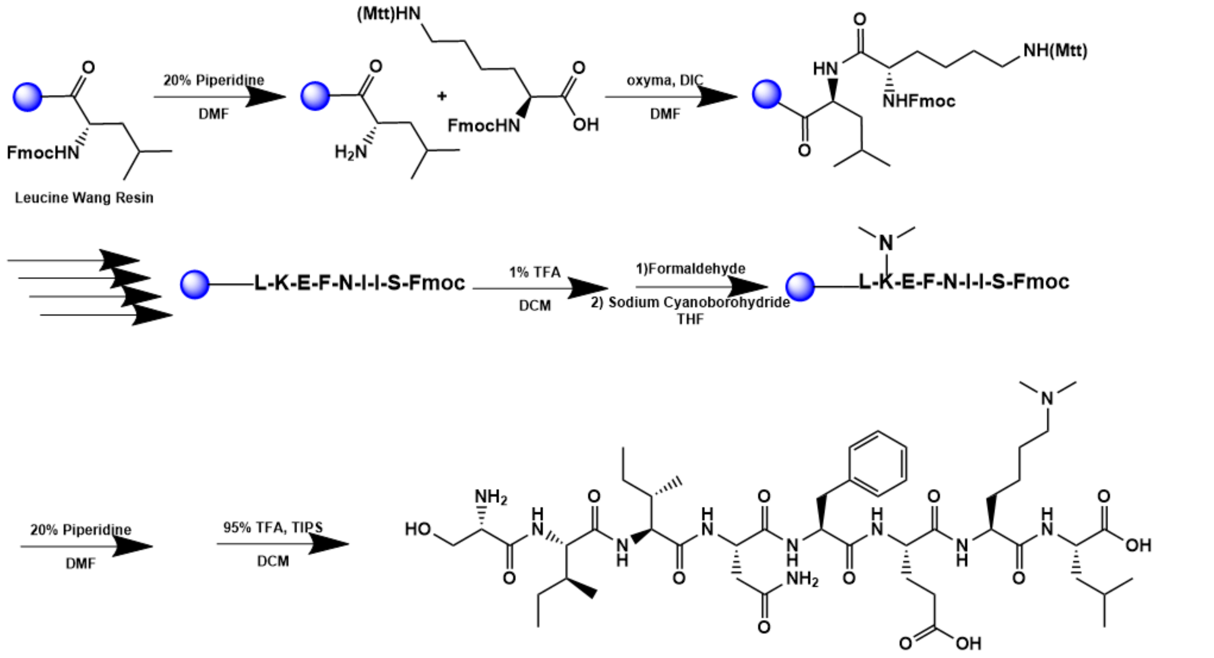


A 25 mL vessel of CEM discover bio manual peptide synthesizer was charged with 0.25 mmol of leucine wang resin. The Fmoc group was removed by using a 20% piperidine solution in DMF (10 mL). Using Synergy software, the deprotection protocol was run. The piperidine solution was drained and the resin was washed with DMF (4 x 10 mL). Fmoc-L-lysine(Mtt)-OH (5 eq, 1.25 mM) along with Oxyma (5 eq, 1.25 mM) and DIC (5 eq, 1.35 mmol) in DMF was added to the reaction vessel and the coupling protocol was run. The amino acid solution was drained, and the resin was washed with DMF (2 x 10 mL). The fmoc removal and coupling procedure was repeated as before using the same equivalencies for the remaining amino acids. The MTT protecting group was removed by the addition of 1% TFA, 2.5% TIPS, in 10 mL DCM for 15 min, washed and repeated 5 more times. Peptide solution was reacted with formaldehyde (10 eq, 12.5 mM) in THF at pH 3 for 15 mins. Sodium cyanoborohydride (20 eq, 25 mM) was added to reaction vessel and reacted for 3 hours and washed 3x with DCM and methanol. To remove the peptide from resin, a TFA cocktail solution (95% TFA, 2.5% TIPS, and 2.5% DCM) was added to the resin and agitated for 2 hours. The resin was filtered, and the resulting solution was concentrated in vacuo. The peptide was trituated with cold diethyl ether and purified using reverse phase HPLC using $H_2O/CH_3CN$. The sample was analyzed for purity using a Waters 1525 Binary HPLC Pump using a Phenomenex Luna 5u C8(2) 100A (250 x 4.60 mm) column; gradient eluted with $H_20/CH_3CN$. Molecular weight was confirmed using high resolution electrospray ionization mass spectrometry (HRMS, ESI/MS) analyses obtained on an Agilent 6545B Q-TOF LC/MS equipped with 1260 infinity II LC system with auto sampler. The final peptide product was lyophilized and stored at -20$^{o}$C until further use.

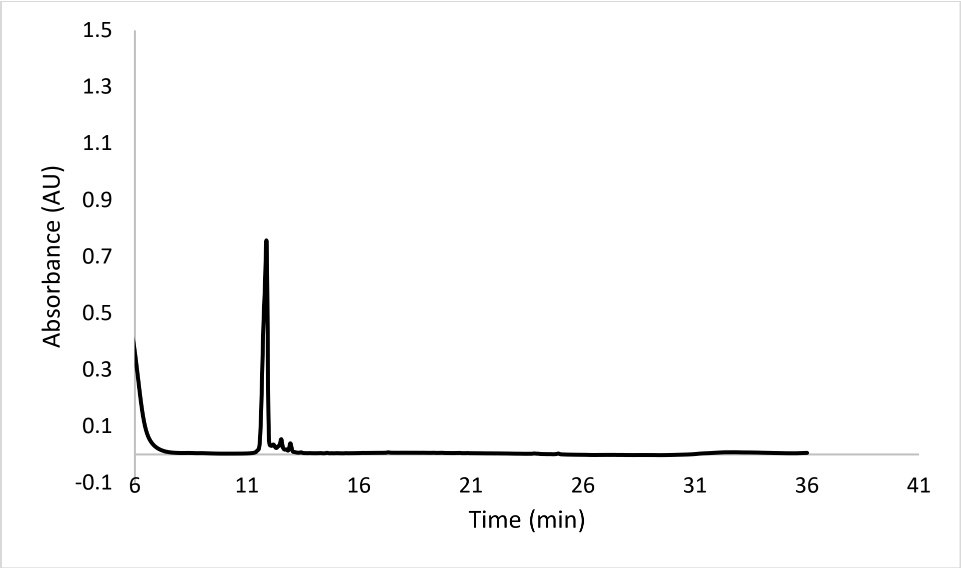


ESI-MS calculated [M+H$^+$]: 991.5828, found 991.5800.

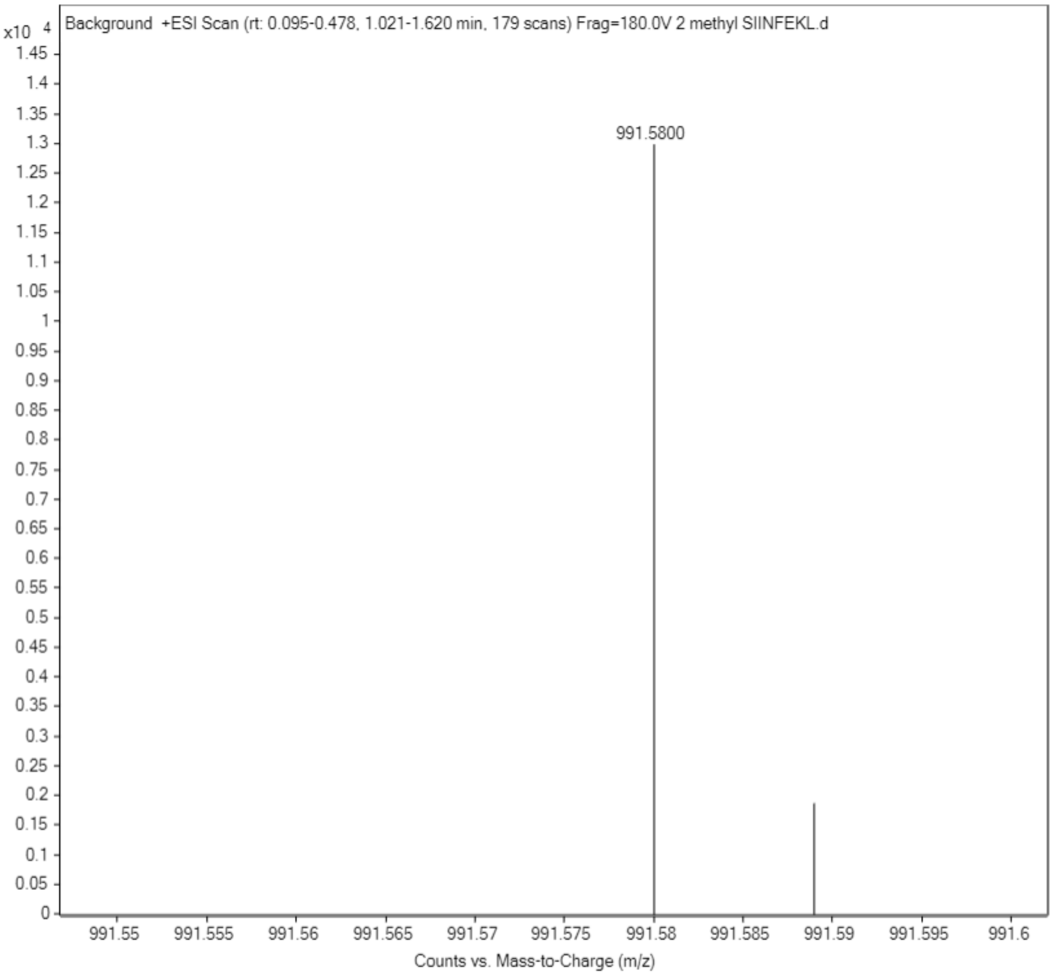

**Scheme S4. Synthesis of Trimethyl Lysine SIINFKEL**

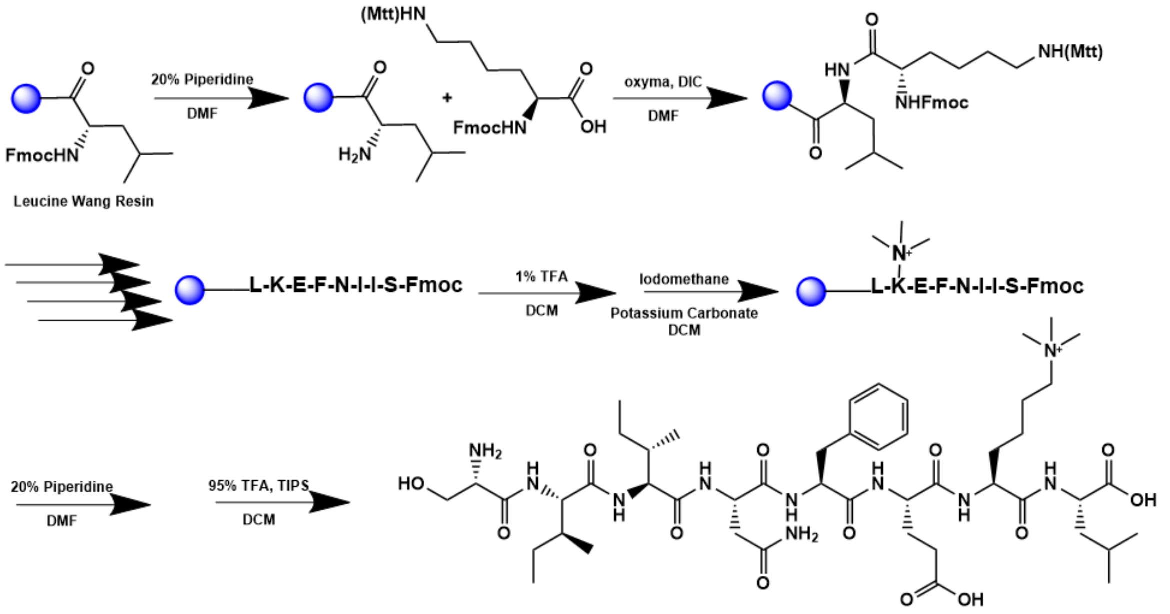


A 25 mL vessel of CEM discover bio manual peptide synthesizer was charged with 0.25 mmol of leucine wang resin. The Fmoc group was removed by using a 20% piperidine solution in DMF (10 mL). Using Synergy software, the deprotection protocol was run. The piperidine solution was drained and the resin was washed with DMF (4 x 10 mL). Fmoc-L-lysine(Mtt)-OH (5 eq, 1.25 mM) along with Oxyma (5 eq, 1.25 mM) and DIC (5 eq, 1.35 mmol) in DMF was added to the reaction vessel and the coupling protocol was run. The amino acid solution was drained, and the resin was washed with DMF (2 x 10 mL). The fmoc removal and coupling procedure was repeated as before using the same equivalencies for the remaining amino acids. The MTT protecting group was removed by the addition of 1% TFA, 2.5% TIPS, in 10 mL DCM for 15 min, washed and repeated 5 more times. Resin was transferred to a round bottom flask and stired in a solution of iodomethane (50 eq, 12.5 mM) and potassium carbonate (10 eq, 2.5 mM) in DCM and heated to 90°C overnight and repeated three times with fresh reagent. To remove the peptide from resin, a TFA cocktail solution (95% TFA, 2.5% TIPS, and 2.5% DCM) was added to the resin and agitated for 2 hours. The resin was filtered, and the resulting solution was concentrated in vacuo. The peptide was trituated with cold diethyl ether and purified using reverse phase HPLC using $H_2O/CH_3CN$. The sample was analyzed for purity using a Waters 1525 Binary HPLC Pump using a Phenomenex Luna 5u C8(2) 100A (250 x 4.60 mm) column; gradient eluted with $H_20/CH_3CN$. Molecular weight was confirmed using high resolution electrospray ionization mass spectrometry (HRMS, ESI/MS) analyses obtained on an Agilent 6545B Q-TOF LC/MS equipped with 1260 infinity II LC system with auto sampler. The final peptide product was lyophilized and stored at -20°C until further use.

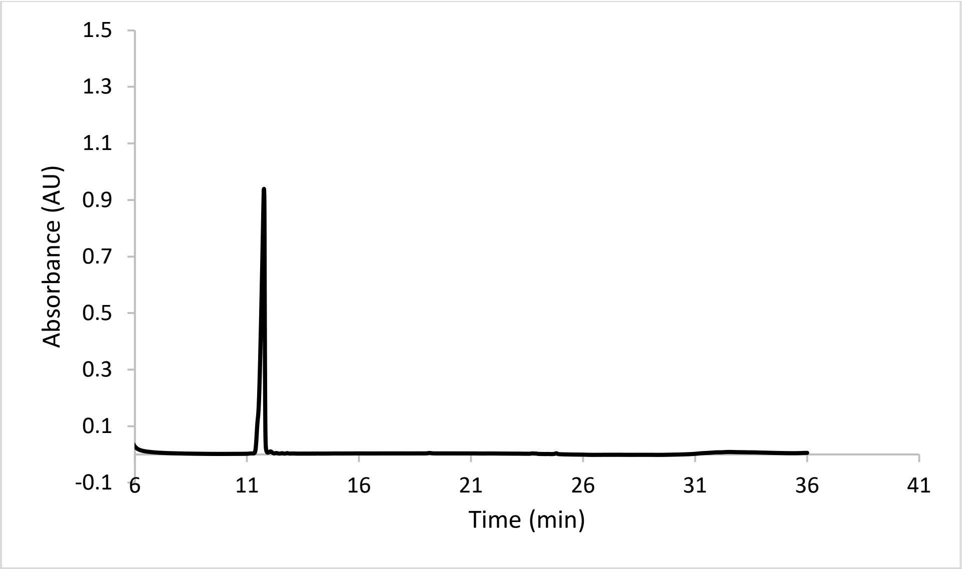


ESI-MS calculated [M+H$^+$]: 1006.6063, found 1006.6009.

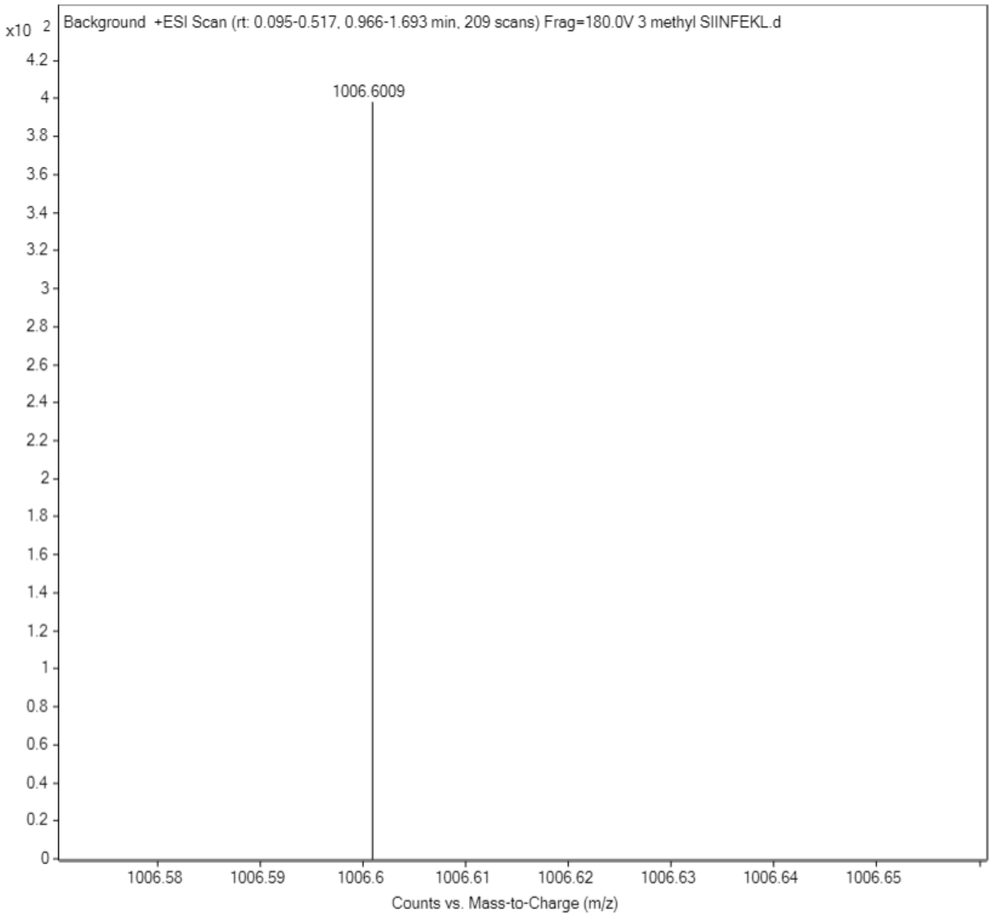

**Scheme S5. Synthesis of Succinyl Lysine SIINFKEL**

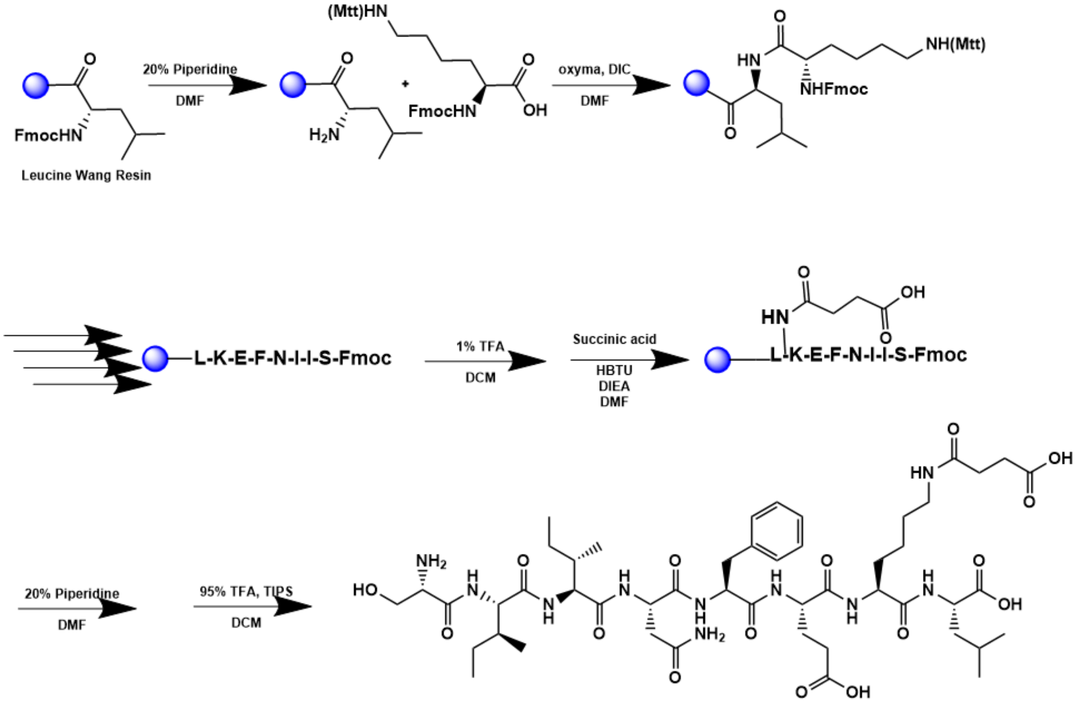


A 25 mL vessel of CEM discover bio manual peptide synthesizer was charged with 0.25 mmol of leucine wang resin. The Fmoc group was removed by using a 20% piperidine solution in DMF (10 mL). Using Synergy software, the deprotection protocol was run. The piperidine solution was drained and the resin was washed with DMF (4 x 10 mL). Fmoc-L-lysine(Mtt)-OH (5 eq, 1.25 mM) along with Oxyma (5 eq, 1.25 mM) and DIC (5 eq, 1.35 mmol) in DMF was added to the reaction vessel and the coupling protocol was run. The amino acid solution was drained, and the resin was washed with DMF (2 x 10 mL). The fmoc removal and coupling procedure was repeated as before using the same equivalencies for the remaining amino acids. The Mtt protecting group of L-Lysine(Mtt)-OH was removed by adding 10 mL of a TFA cocktail solution (1% TFA, 2% TIPS in DCM) to the resin and agitating for 10 minutes protected from light. The solution was drained, and this procedure was repeated five additional times. Succinic acid (8 eq, 2 mM) along with Oxyma (5 eq, 1.25 mM) and DIC (5 eq, 1.25 mmol) in DMF was added to the reaction vessel and agitated at room temperature for 2 hours. To remove the peptide from resin, a TFA cocktail solution (95% TFA, 2.5% TIPS, and 2.5% DCM) was added to the resin and agitated for 2 hours. The resin was filtered, and the resulting solution was concentrated in vacuo. The peptide was trituated with cold diethyl ether and purified using reverse phase HPLC using $H_2O/CH_3CN$. The sample was analyzed for purity using a Waters 1525 Binary HPLC Pump using a Phenomenex Luna 5u C8(2) 100A (250 x 4.60 mm) column; gradient eluted with $H_20/CH_3CN$. Molecular weight was confirmed using high resolution electrospray ionization mass spectrometry (HRMS, ESI/MS) analyses obtained on an Agilent 6545B Q-TOF LC/MS equipped with 1260 infinity II LC system with auto sampler. The final peptide product was lyophilized and stored at -20$^{o}$C until further use.

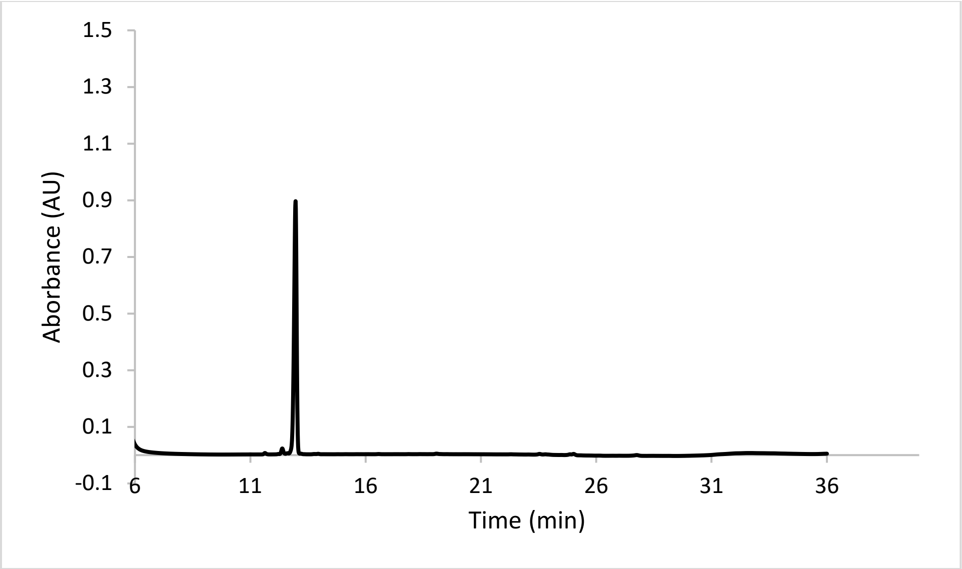


ESI-MS calculated [M+H$^+$]: 1063.5675, found 1063.5656.

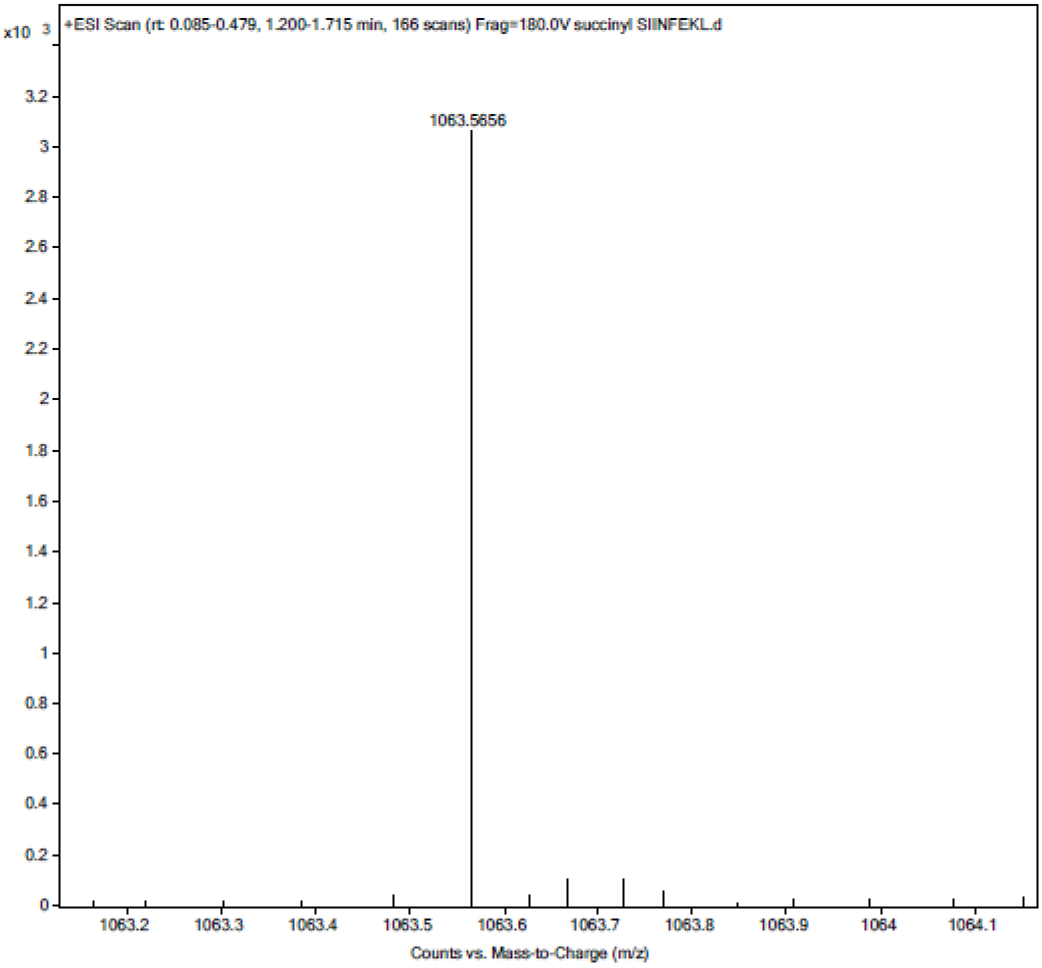

**Scheme S6. Synthesis of Acetyl Lysine SIINFKEL**

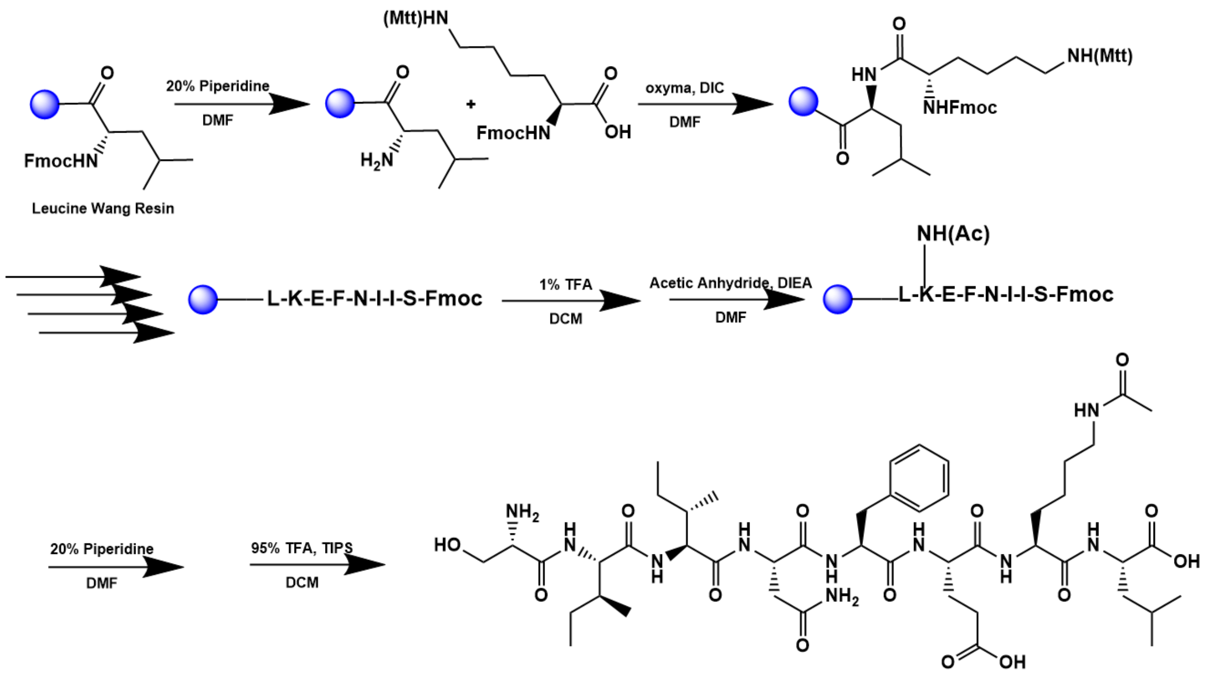


A 25 mL vessel of CEM discover bio manual peptide synthesizer was charged with 0.25 mmol of leucine wang resin. The Fmoc group was removed by using a 20% piperidine solution in DMF (10 mL). Using Synergy software, the deprotection protocol was run. The piperidine solution was drained and the resin was washed with DMF (4 x 10 mL). Fmoc-L-lysine(Mtt)-OH (5 eq, 1.25 mM) along with Oxyma (5 eq, 1.25 mM) and DIC (5 eq, 1.35 mmol) in DMF was added to the reaction vessel and the coupling protocol was run. The amino acid solution was drained, and the resin was washed with DMF (2 x 10 mL). The fmoc removal and coupling procedure was repeated as before using the same equivalencies for the remaining amino acids. The Mtt protecting group of lysine was removed by adding 10 mL of a TFA cocktail solution (1% TFA, 2% TIPS in DCM) to the resin and agitating for 10 minutes. The solution was drained, and this procedure was repeated five additional times. The resin was transferred to a 25 mL synthetic vessel and the lysine side chain of the peptide was acetylated agitating the resin for 1 hour in a solution of 5% acetic anhydride (0.5 mL), 8.5% DIEA (0.85 mL), and 86.5% DMF (8.65 mL). To remove the peptide from resin, a TFA cocktail solution (95% TFA, 2.5% TIPS, and 2.5% DCM) was added to the resin and agitated for 2 hours. The resin was filtered, and the resulting solution was concentrated in vacuo. The peptide was trituated with cold diethyl ether and purified using reverse phase HPLC using $H_2O/CH_3CN$. The sample was analyzed for purity using a Waters 1525 Binary HPLC Pump using a Phenomenex Luna 5u C8(2) 100A (250 x 4.60 mm) column; gradient eluted with $H_20/CH_3CN$. Molecular weight was confirmed using high resolution electrospray ionization mass spectrometry (HRMS, ESI/MS) analyses obtained on an Agilent 6545B Q-TOF LC/MS equipped with 1260 infinity II LC system with auto sampler. The final peptide product was lyophilized and stored at -20$^{o}$C until further use.

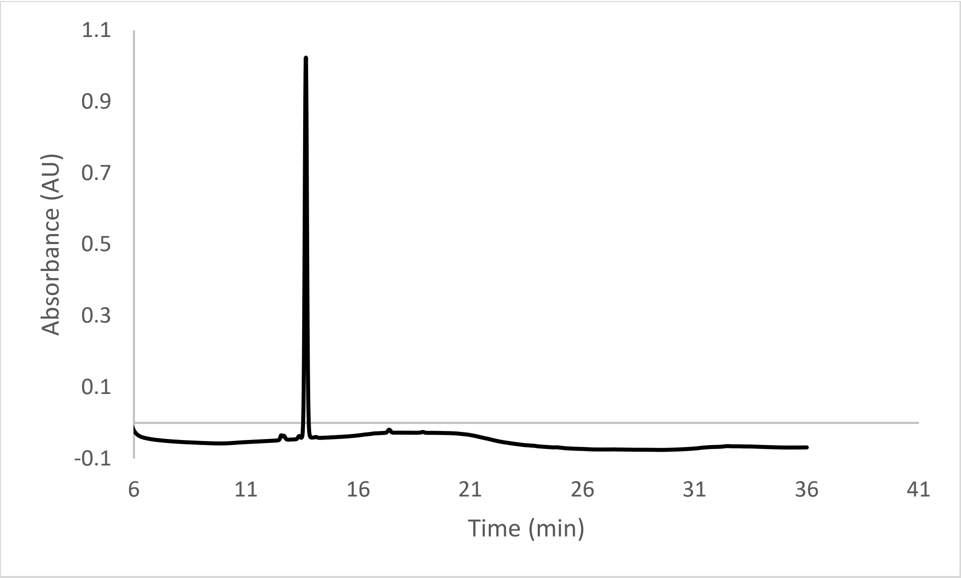


ESI-MS calculated [M+H$^+$]: 1005.5620, found 1005.5586.

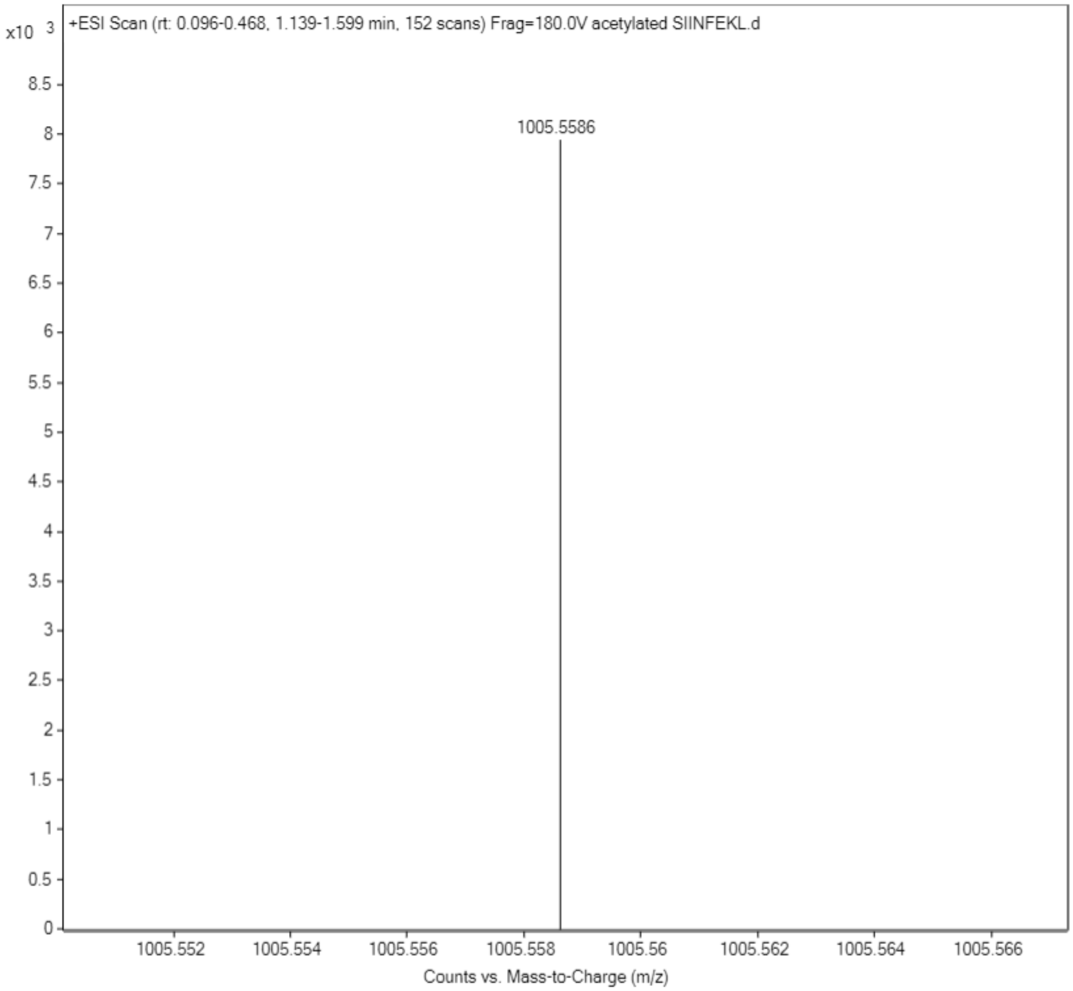

**Scheme S7. Synthesis of Biotinylated Lysine SIINFKEL**

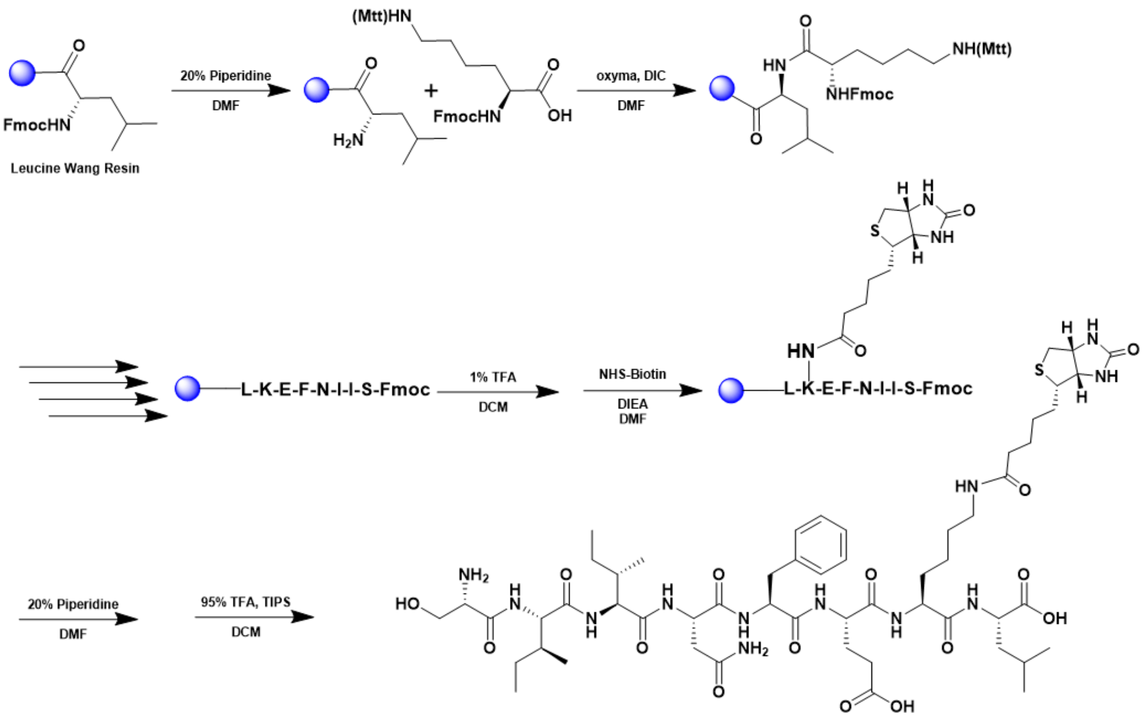


A 25 mL vessel of CEM discover bio manual peptide synthesizer was charged with 0.25 mmol of leucine wang resin. The Fmoc group was removed by using a 20% piperidine solution in DMF (10 mL). Using Synergy software, the deprotection protocol was run. The piperidine solution was drained and the resin was washed with DMF (4 x 10 mL). Fmoc-L-lysine(Mtt)-OH (5 eq, 1.25 mM) along with Oxyma (5 eq, 1.25 mM) and DIC (5 eq, 1.35 mmol) in DMF was added to the reaction vessel and the coupling protocol was run. The amino acid solution was drained, and the resin was washed with DMF (2 x 10 mL). The fmoc removal and coupling procedure was repeated as before using the same equivalencies for the remaining amino acids. The Mtt protecting group of lysine was removed by adding 10 mL of a TFA cocktail solution (1% TFA, 2% TIPS in DCM) to the resin and agitating for 10 minutes. The solution was drained, and this procedure was repeated five additional times. Biotin N-hydroxy succinimide ester (10 eq, 2.5 mM) and DIEA (4 eq, 1 mM) in DMF were reacted with peptide for 2 h while agitated. To remove the peptide from resin, a TFA cocktail solution (95% TFA, 2.5% TIPS, and 2.5% DCM) was added to the resin and agitated for 2 hours. The resin was filtered, and the resulting solution was concentrated in vacuo. The peptide was trituated with cold diethyl ether and purified using reverse phase HPLC using $H_2O/CH_3CN$. The sample was analyzed for purity using a Waters 1525 Binary HPLC Pump using a Phenomenex Luna 5u C8(2) 100A (250 x 4.60 mm) column; gradient eluted with $H_20/CH_3CN$. Molecular weight was confirmed using high resolution electrospray ionization mass spectrometry (HRMS, ESI/MS) analyses obtained on an Agilent 6545B Q-TOF LC/MS equipped with 1260 infinity II LC system with auto sampler. The final peptide product was lyophilized and stored at -20°C until further use.

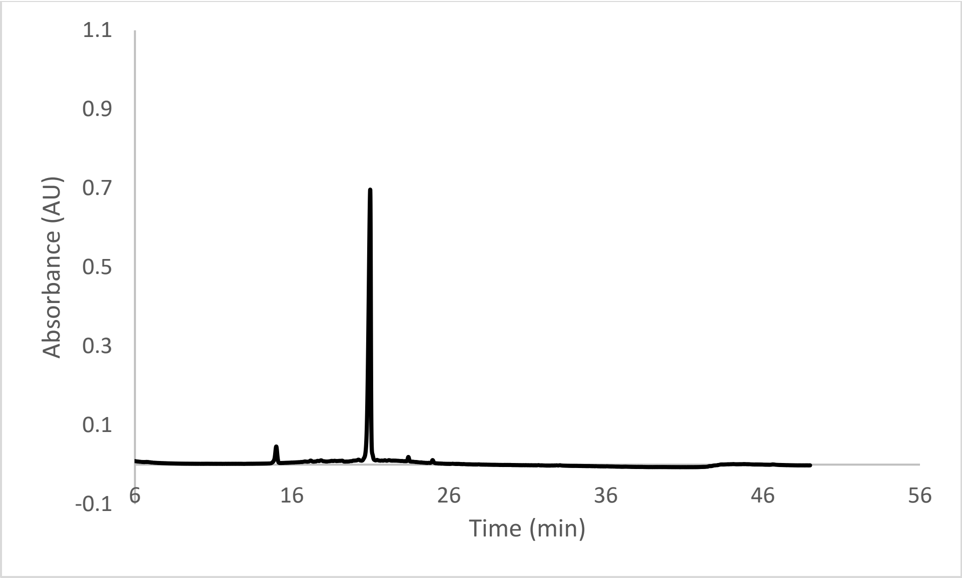


ESI-MS calculated [M+H⁺]: 1189.6291, found 1189.6272.

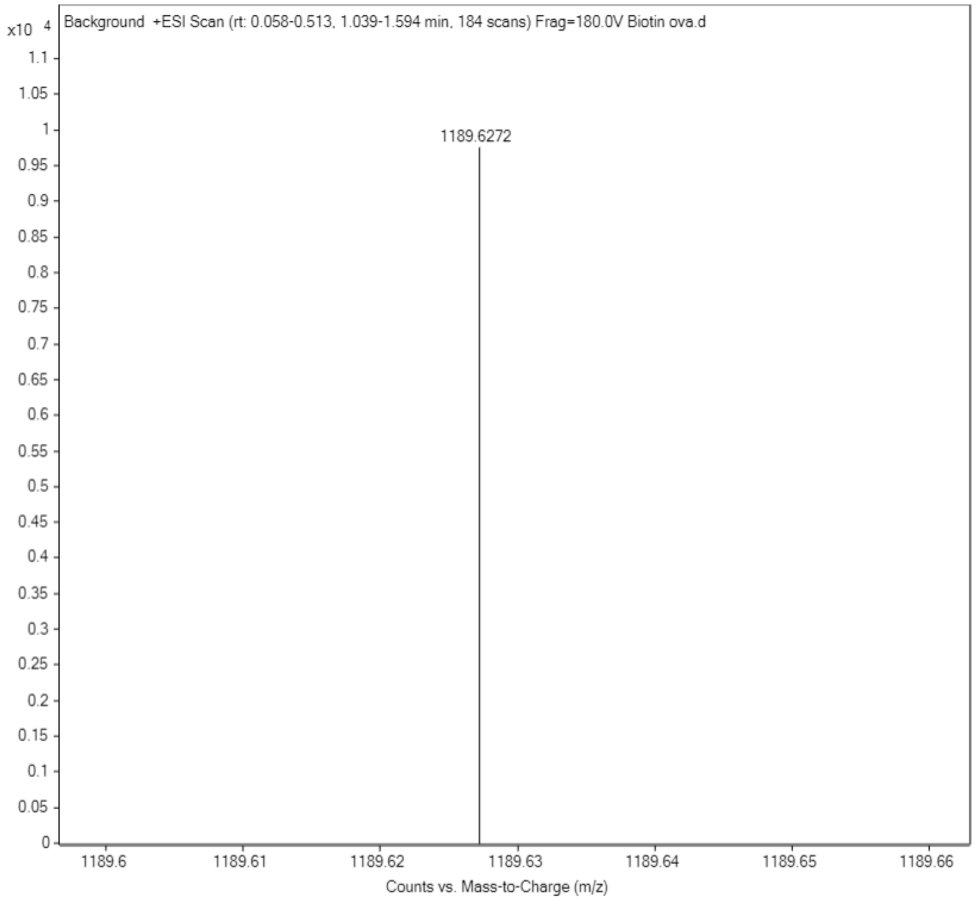

**Scheme S8. Synthesis of Phosphoserine SIINFKEL**

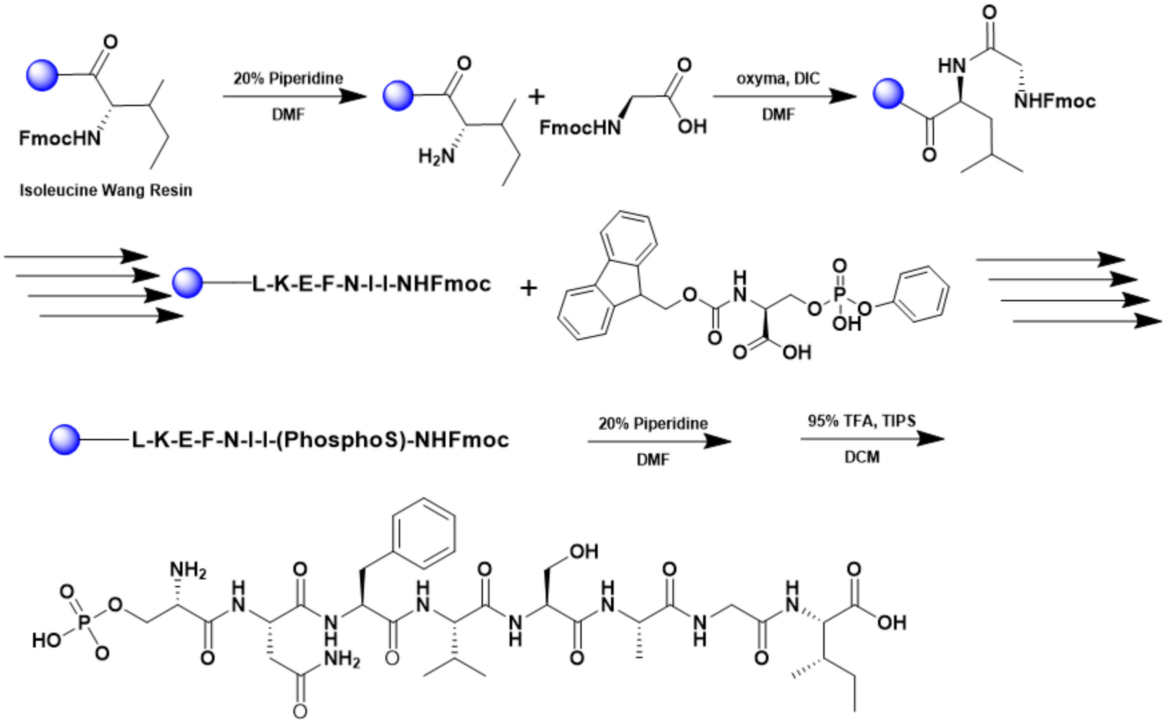


A 25 mL vessel of CEM discover bio manual peptide synthesizer was charged with 0.25 mmol of leucine wang resin. The Fmoc group was removed by using a 20% piperidine solution in DMF (10 mL). Using Synergy software, the deprotection protocol was run. The piperidine solution was drained and the resin was washed with DMF (4 x 10 mL). Fmoc-L-lysine(Mtt)-OH (5 eq, 1.25 mM) along with Oxyma (5 eq, 1.25 mM) and DIC (5 eq, 1.35 mmol) in DMF was added to the reaction vessel and the coupling protocol was run. The amino acid solution was drained, and the resin was washed with DMF (2 x 10 mL). The fmoc removal and coupling procedure was repeated as before using the same equivalencies for the remaining amino acids. To remove the peptide from resin, a TFA cocktail solution (95% TFA, 2.5% TIPS, and 2.5% DCM) was added to the resin and agitated for 2 hours. The resin was filtered, and the resulting solution was concentrated in vacuo. The peptide was triturated with cold diethyl ether and purified using reverse phase HPLC using $H_2O/CH_3CN$. The sample was analyzed for purity using a Waters 1525 Binary HPLC Pump using a Phenomenex Luna 5u C8(2) 100A (250 x 4.60 mm) column; gradient eluted with $H_20/CH_3CN$. Molecular weight was confirmed using high resolution electrospray ionization mass spectrometry (HRMS, ESI/MS) analyses obtained on an Agilent 6545B Q-TOF LC/MS equipped with 1260 infinity II LC system with auto sampler. The final peptide product was lyophilized and stored at -20°C until further use.

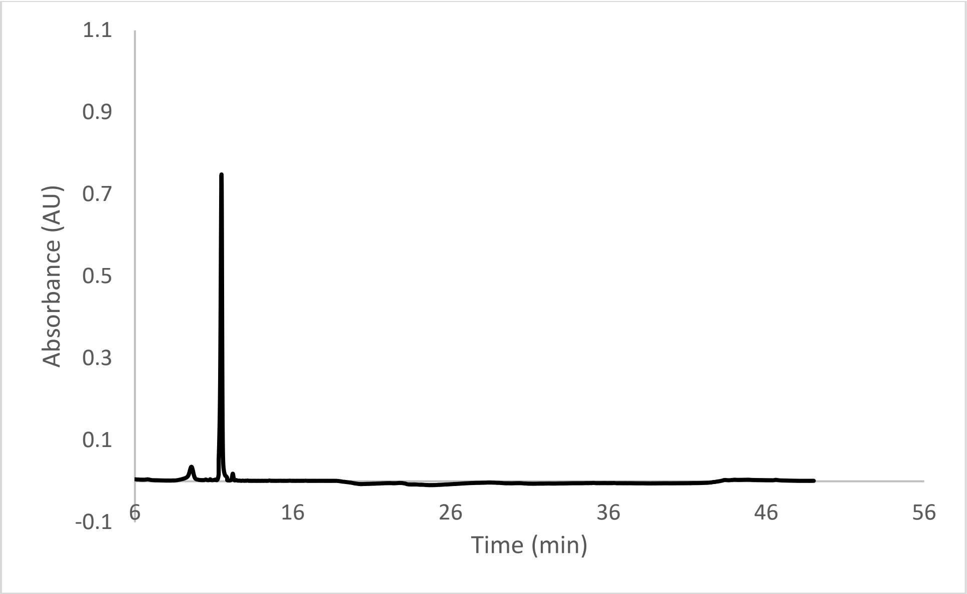


ESI-MS calculated [M+H$^+$]: 1042.5100, found 1042.5072.

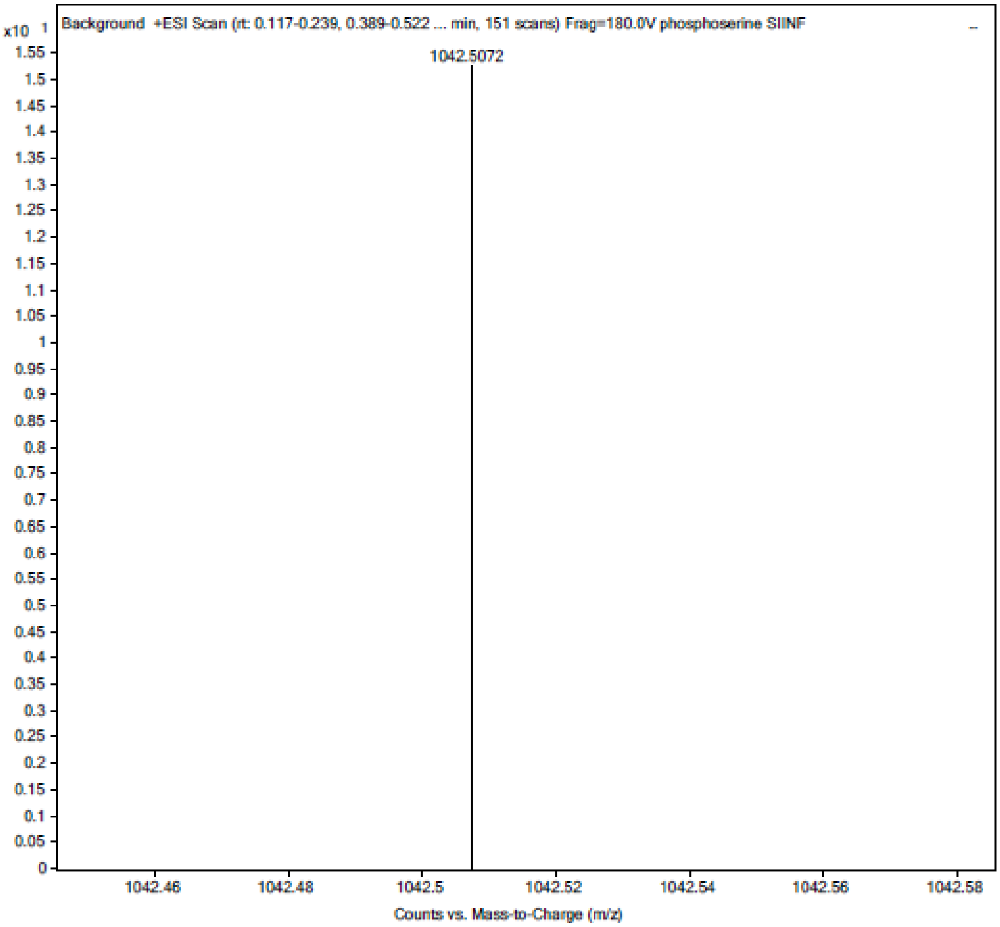

**Scheme S9. Synthesis of SNFVSAGI**

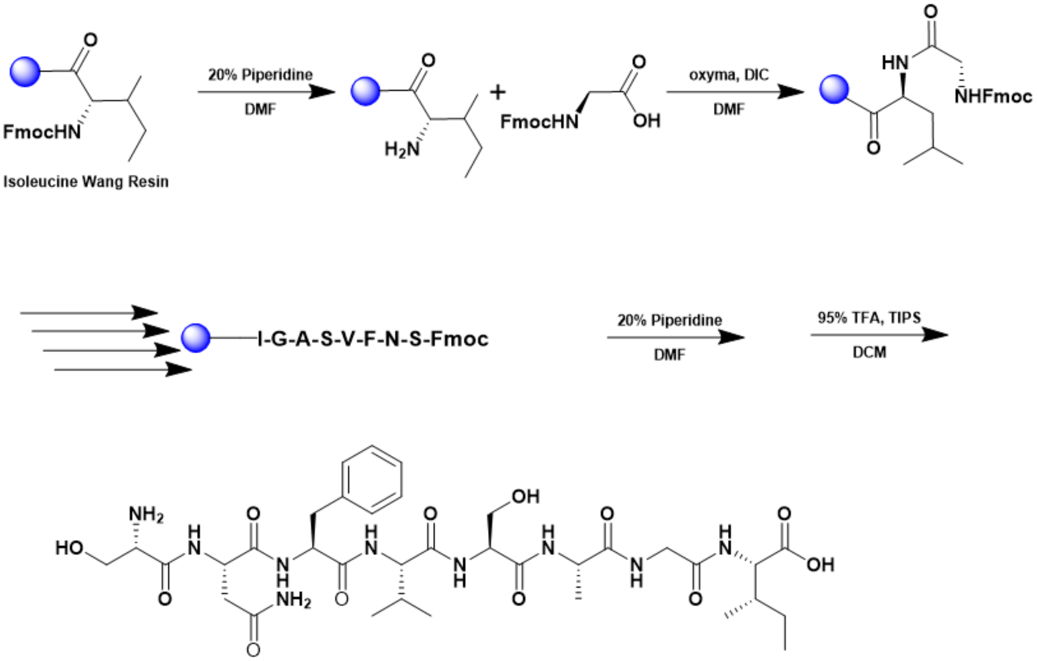


A 25 mL vessel of CEM discover bio manual peptide synthesizer was charged with 0.25 mmol of isoleucine wang resin. The Fmoc group was removed by using a 20% piperidine solution in DMF (10 mL). Using Synergy software, the deprotection protocol was run. The piperidine solution was drained and the resin was washed with DMF (4 x 10 mL). Fmoc-L-glycine (5 eq, 1.25 mM) along with Oxyma (5 eq, 1.25 mM) and DIC (5 eq, 1.35 mmol) in DMF was added to the reaction vessel and the coupling protocol was run. The amino acid solution was drained, and the resin was washed with DMF (2 x 10 mL). The fmoc removal and coupling procedure was repeated as before using the same equivalencies for the remaining amino acids. To remove the peptide from resin, a TFA cocktail solution (95% TFA, 2.5% TIPS, and 2.5% DCM) was added to the resin and agitated for 2 hours. The resin was filtered, and the resulting solution was concentrated in vacuo. The peptide was trituated with cold diethyl ether and purified using reverse phase HPLC using $H_2O/CH_3CN$. The sample was analyzed for purity using a a Waters 1525 Binary HPLC Pump using a Phenomenex Luna 5u C8(2) 100A (250 x 4.60 mm) column; gradient eluted with $H_20/CH_3CN$. Molecular weight was confirmed using high resolution electrospray ionization mass spectrometry (HRMS, ESI/MS) analyses obtained on an Agilent 6545B Q-TOF LC/MS equipped with 1260 infinity II LC system with auto sampler. The final peptide product was lyophilized and stored at -20$^o$C until further use.

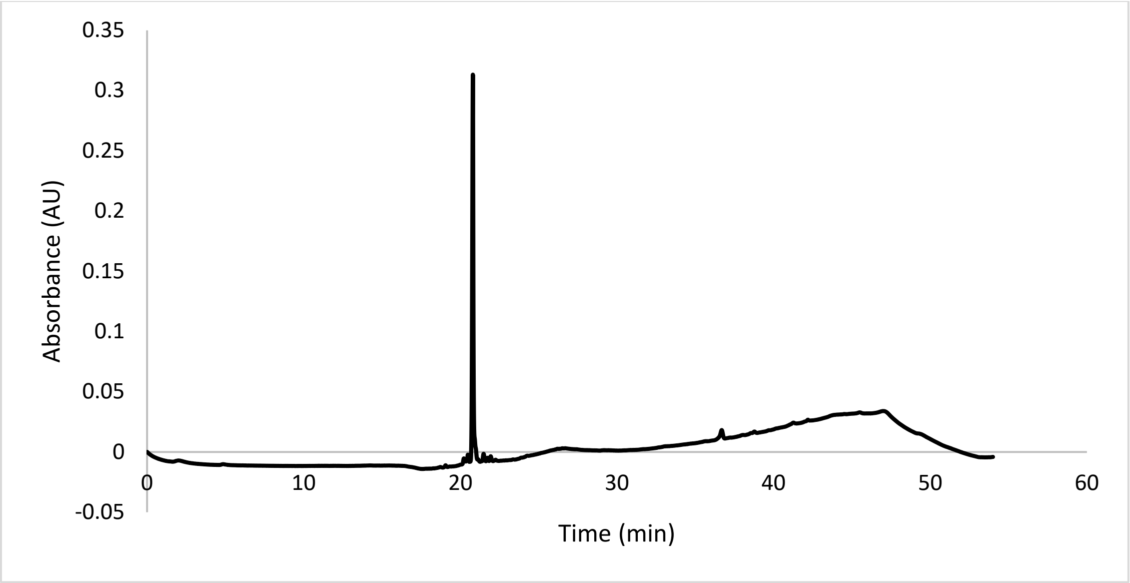


ESI-MS calculated [M+H$^+$]: 794.4048, found 794.4038

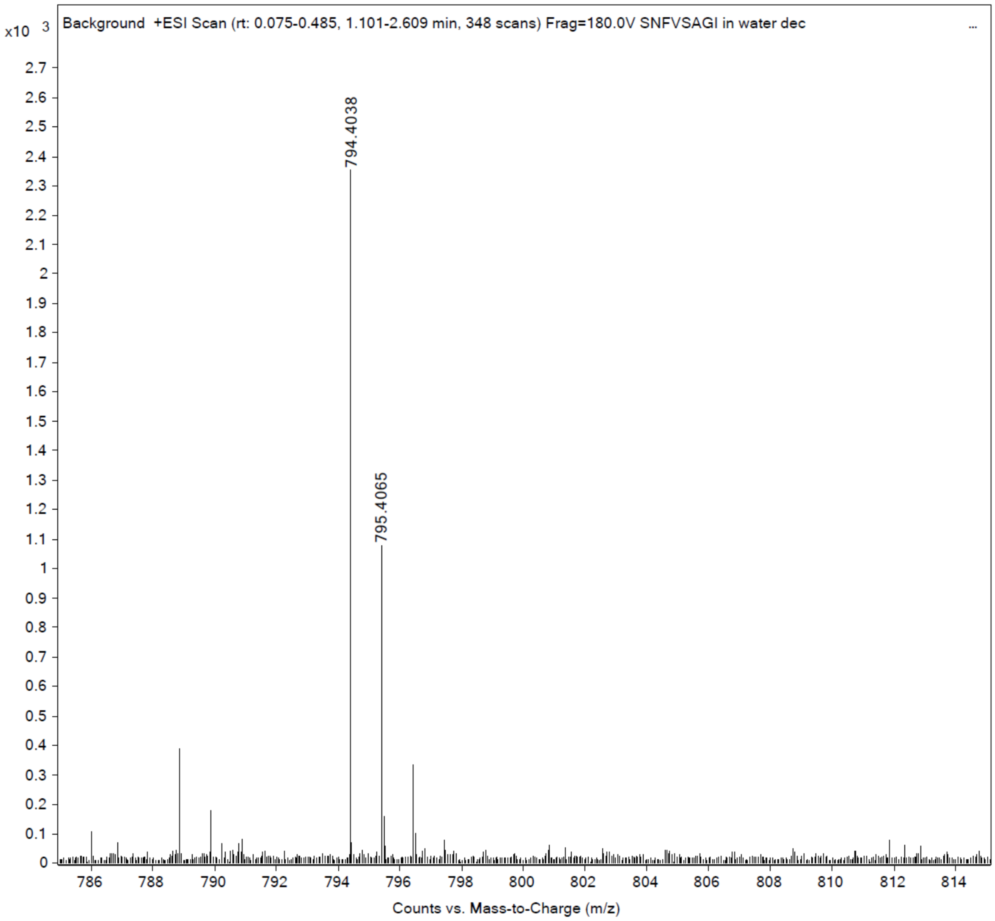

**Scheme S10. Synthesis of ESIVRFPNI**

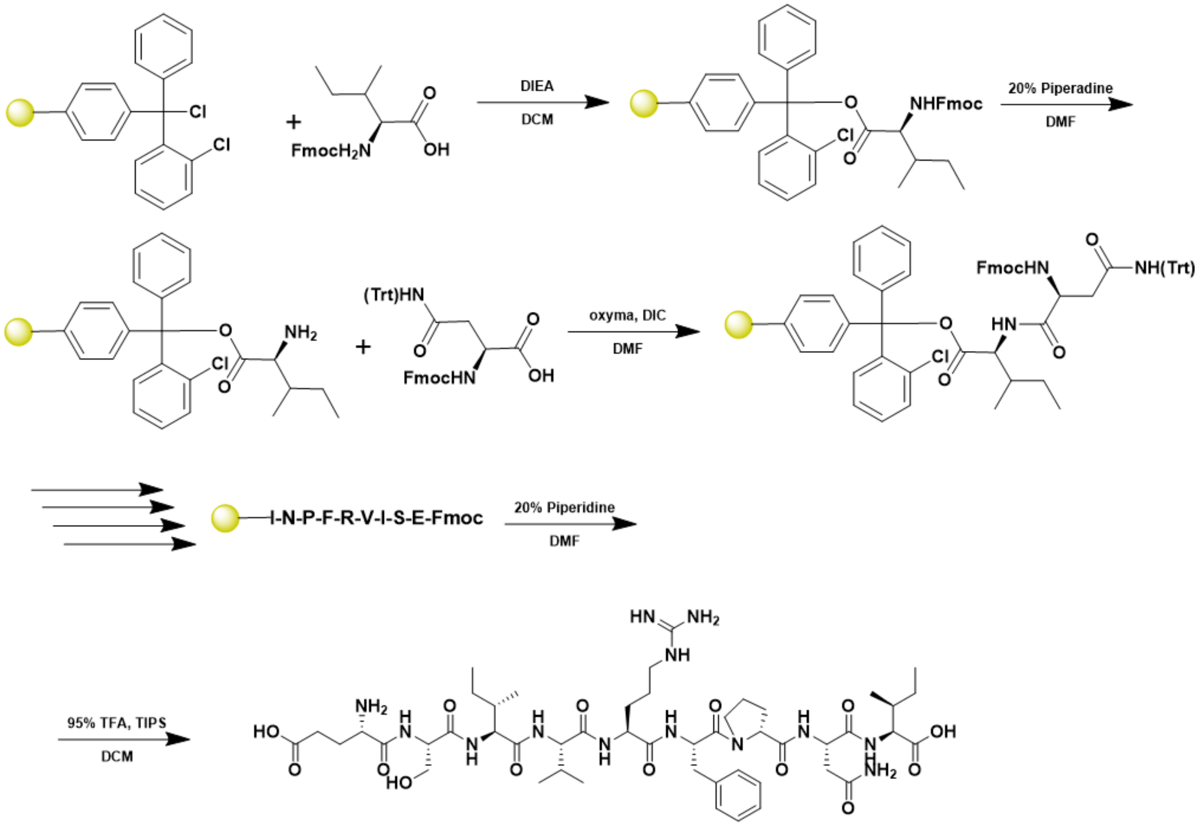


A 25 mL peptide synthesis vessel charged with 2-hlorotrityl chloride resin (0.25mmol) was added Fmoc-L-isoleucine-OH (1.1 eq, 0.275 mmol) and DIEA (3 eq, 0.75 mmol) in dry DCM. The resin was agitated for 1 h at ambient temperature and washed with MeOH and DCM (3 x each). The Fmoc group was removed by using a 20% piperidine solution in DMF for 30 min at ambient temperature, then washed as before. Fmoc-L-asparagine(Trt)-OH (5 eq, 1.25 mM) along with Oxyma (5 eq, 1.25 mM) and DIC (5 eq, 1.35 mmol) in DMF was added to the reaction vessel and agitated for 2 h at ambient temperature. The Fmoc removal and coupling procedure was repeated as before using the same equivalencies for the remaining amino acids. To remove the peptide from resin, a TFA cocktail solution (95% TFA, 2.5% TIPS, and 2.5% DCM) was added to the resin and agitated for 2 hours. The resin was filtered, and the resulting solution was concentrated in vacuo. The peptide was trituated with cold diethyl ether and purified using reverse phase HPLC using $H_2O/CH_3CN$. The sample was analyzed for purity using a Waters 1525 Binary HPLC Pump using a Phenomenex Luna 5u C8(2) 100A (250 x 4.60 mm) column; gradient eluted with $H_20/CH_3CN$. Molecular weight was confirmed using high resolution electrospray ionization mass spectrometry (HRMS, ESI/MS) analyses obtained on an Agilent 6545B Q-TOF LC/MS equipped with 1260 infinity II LC system with auto sampler. The final peptide product was lyophilized and stored at -20°C until further use.

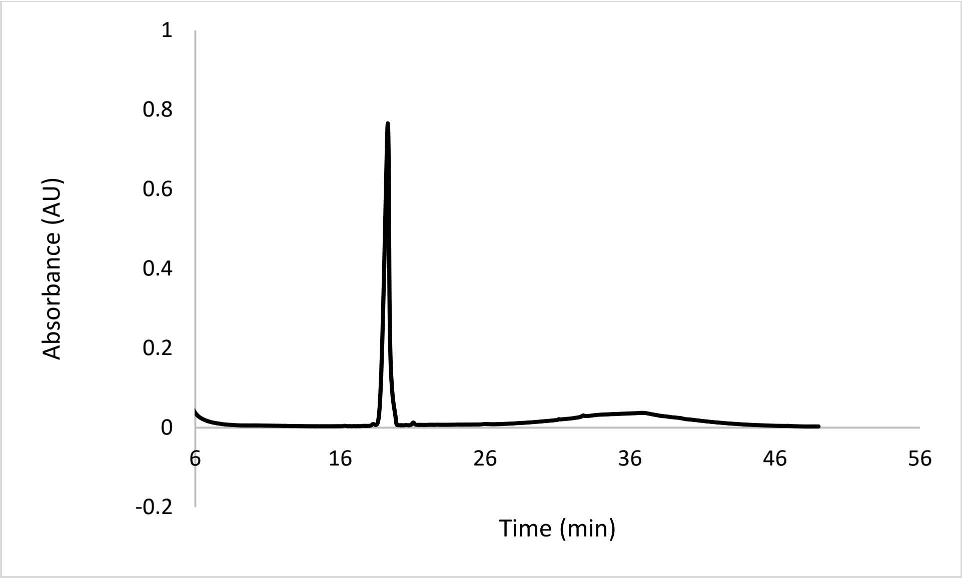


ESI-MS calculated [M]: 1073.5869, found 1073.5870

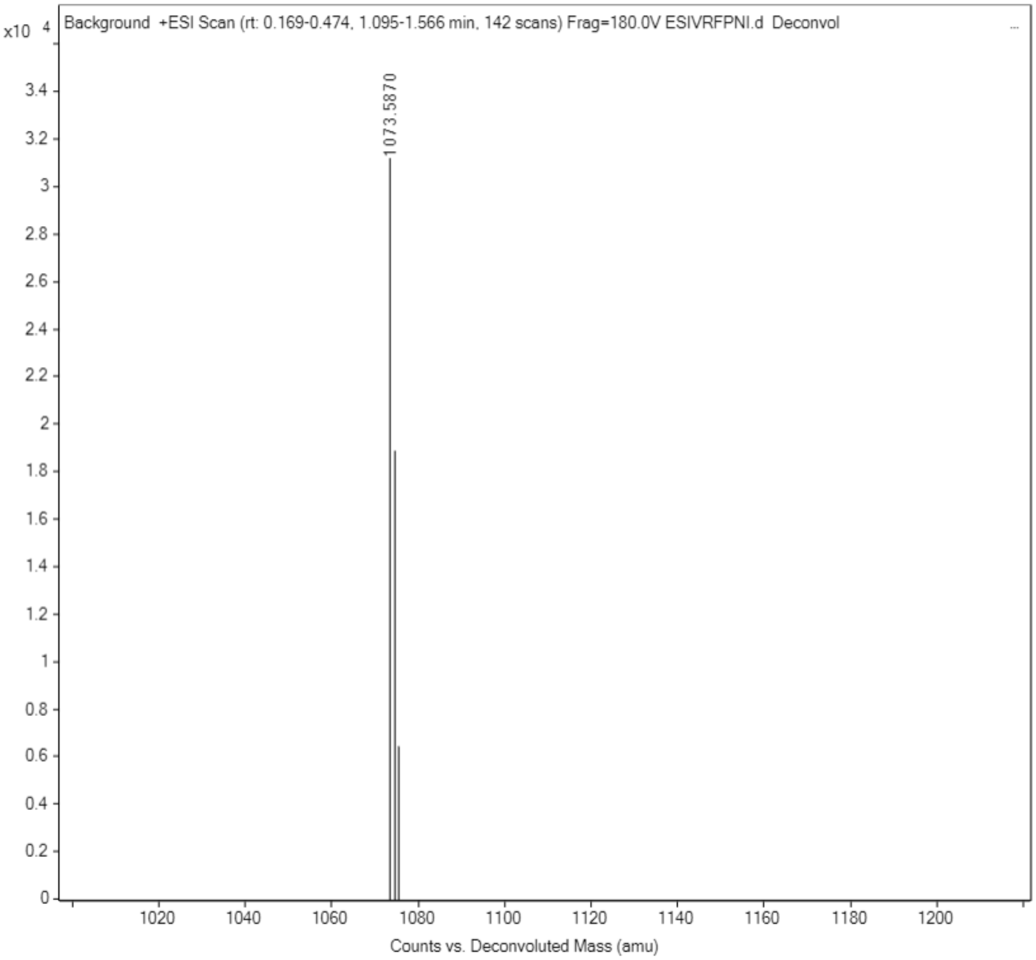

**Scheme S11. Synthesis of N-Acetyl ESIVRFPNI**

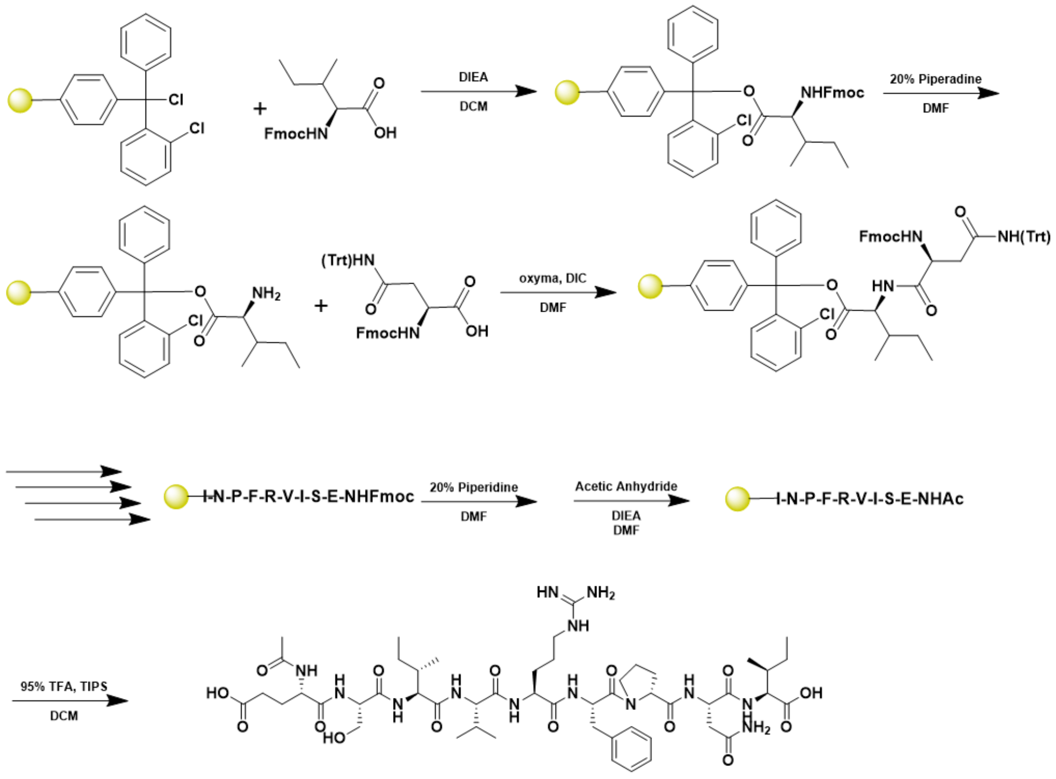


A 25 mL peptide synthesis vessel charged with 2-hlorotrityl chloride resin (0.25mmol) was added Fmoc-L-isoleucine-OH (1.1 eq, 0.275 mmol) and DIEA (3 eq, 0.75 mmol) in dry DCM. The resin was agitated for 1 h at ambient temperature and washed with MeOH and DCM (3 x each). The Fmoc group was removed by using a 20% piperidine solution in DMF for 30 min at ambient temperature, then washed as before. Fmoc-L-asparagine(Trt)-OH (5 eq, 1.25 mM) along with Oxyma (5 eq, 1.25 mM) and DIC (5 eq, 1.35 mmol) in DMF was added to the reaction vessel and agitated for 2 h at ambient temperature. The Fmoc removal and coupling procedure was repeated as before using the same equivalencies for the remaining amino acids. The final amino acid was Fmoc deprotected as described before and was acetylated by agitating the resin for 1 hour in a solution of 5% acetic anhydride (0.5 mL), 8.5% DIEA (0.85 mL), and 86.5% DMF (8.65 mL). To remove the peptide from resin, a TFA cocktail solution (95% TFA, 2.5% TIPS, and 2.5% DCM) was added to the resin and agitated for 2 hours. The resin was filtered, and the resulting solution was concentrated in vacuo. The peptide was trituated with cold diethyl ether and purified using reverse phase HPLC using $H_2O/CH_3CN$. The sample was analyzed for purity using a Waters 1525 Binary HPLC Pump using a Phenomenex Luna 5u C8(2) 100A (250 x 4.60 mm) column; gradient eluted with $H_20/CH_3CN$. Molecular weight was confirmed using high resolution electrospray ionization mass spectrometry (HRMS, ESI/MS) analyses obtained on an Agilent 6545B Q-TOF LC/MS equipped with 1260 infinity II LC system with auto sampler. The final peptide product was lyophilized and stored at -20$^o$C until further use.

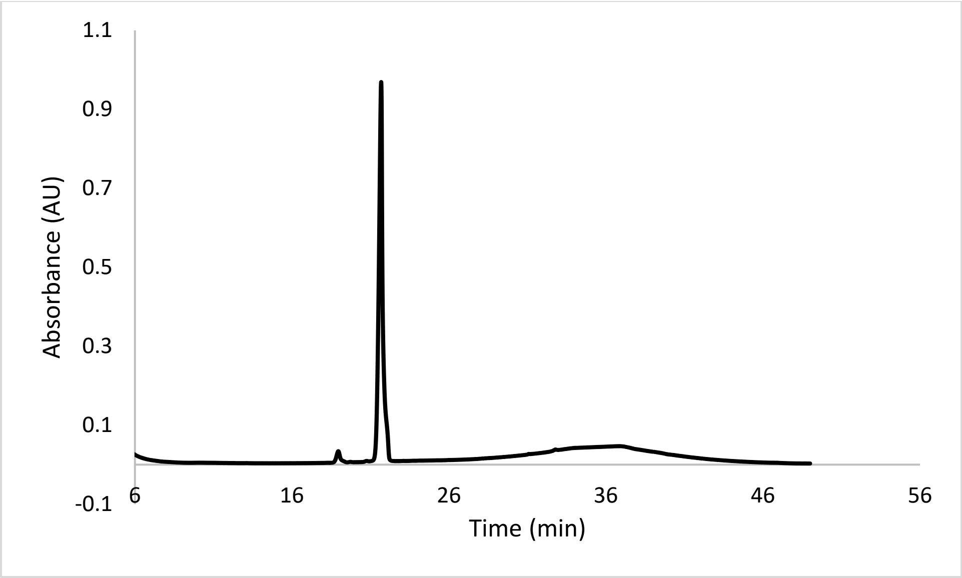


ESI-MS calculated [M]: 1116.6048, found 1116.6039

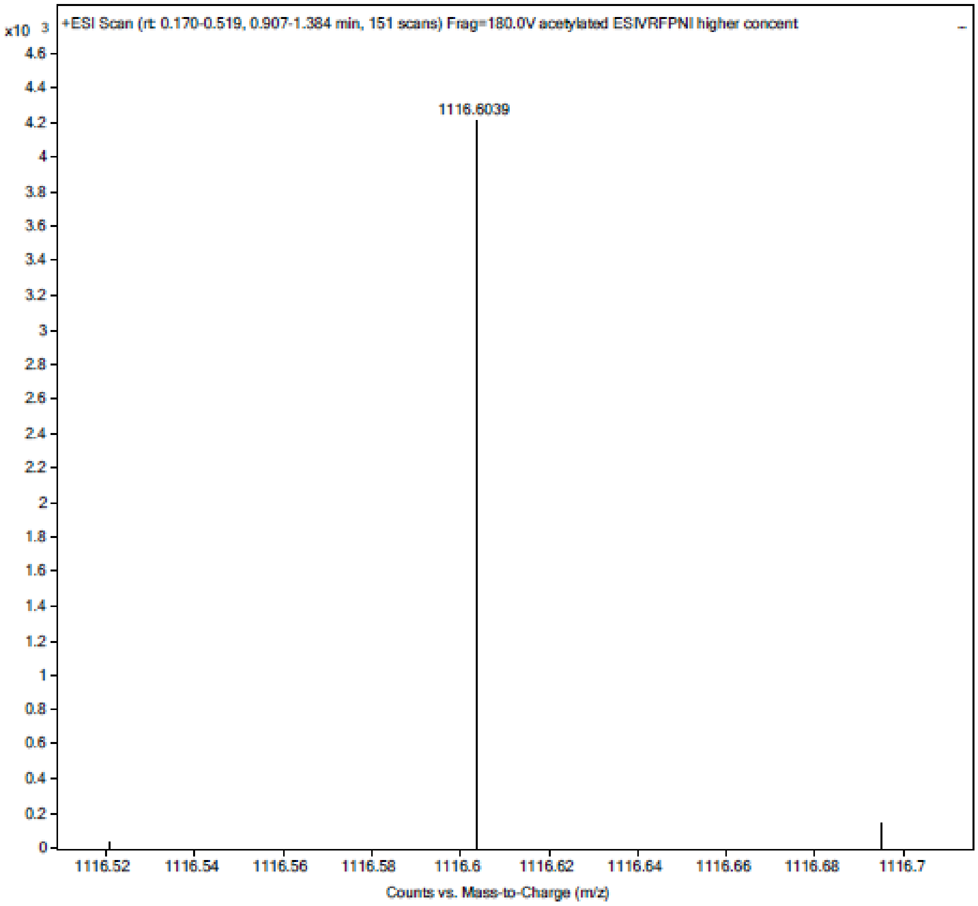


**Scheme S12. Synthesis of Citrullinated ESIVRFPNI**

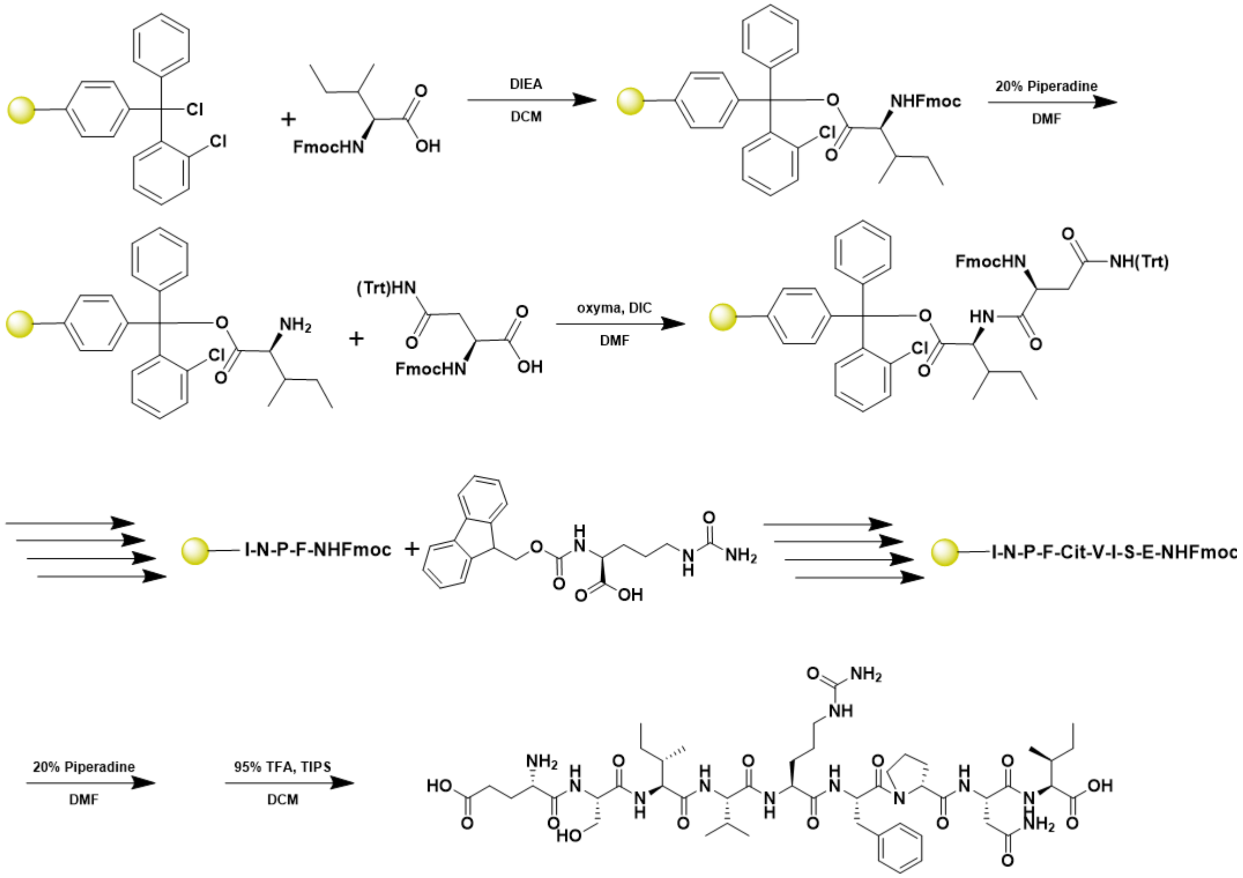


A 25 mL peptide synthesis vessel charged with 2-hlorotrityl chloride resin (0.25mmol) was added Fmoc-L-isoleucine-OH (1.1 eq, 0.275 mmol) and DIEA (3 eq, 0.75 mmol) in dry DCM. The resin was agitated for 1 h at ambient temperature and washed with MeOH and DCM (3 x each). The Fmoc group was removed by using a 20% piperidine solution in DMF for 30 min at ambient temperature, then washed as before. Fmoc-L-asparagine(Trt)-OH (5 eq, 1.25 mM) along with Oxyma (5 eq, 1.25 mM) and DIC (5 eq, 1.35 mmol) in DMF was added to the reaction vessel and agitated for 2 h at ambient temperature. The Fmoc removal and coupling procedure was repeated as before using the same equivalencies for the remaining amino acids. For the 5$^{th}$ position of the amino acid, Fmoc was deprotected as described above and Fmoc-L-citrulline (5 eq, 1.25 mM) along with Oxyma (5 eq, 1.25 mM) and DIC (5 eq, 1.35 mmol) in DMF was added to the reaction vessel. To remove the peptide from resin, a TFA cocktail solution (95% TFA, 2.5% TIPS, and 2.5% DCM) was added to the resin and agitated for 2 hours. The resin was filtered, and the resulting solution was concentrated in vacuo. The peptide was trituated with cold diethyl ether and purified using reverse phase HPLC using $H_2O/CH_3CN$. The sample was analyzed for purity using a Waters 1525 Binary HPLC Pump using a Phenomenex Luna 5u C8(2) 100A (250 x 4.60 mm) column; gradient eluted with $H_20/CH_3CN$. Molecular weight was confirmed using high resolution electrospray ionization mass spectrometry (HRMS, ESI/MS) analyses obtained on an Agilent 6545B Q-TOF LC/MS equipped with 1260 infinity II LC system with auto sampler. The final peptide product was lyophilized and stored at -20$^{o}$C until further use.

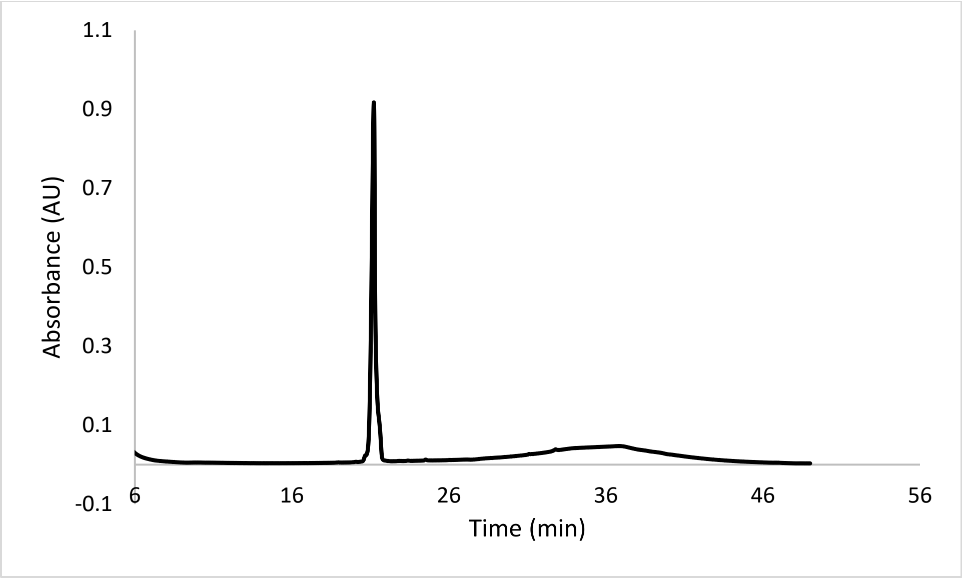


ESI-MS calculated [M]: 1074.5710, found 1074.5708

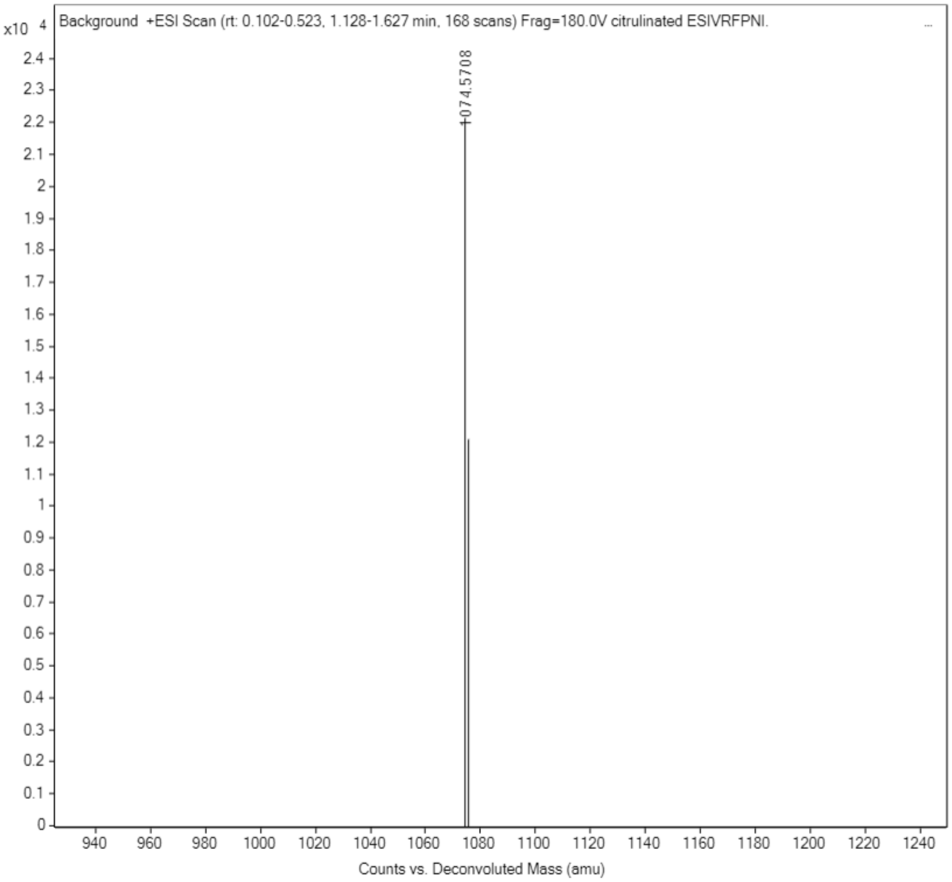

**Scheme S13. Synthesis of Hydroxy Proline ESIVRFPNI**

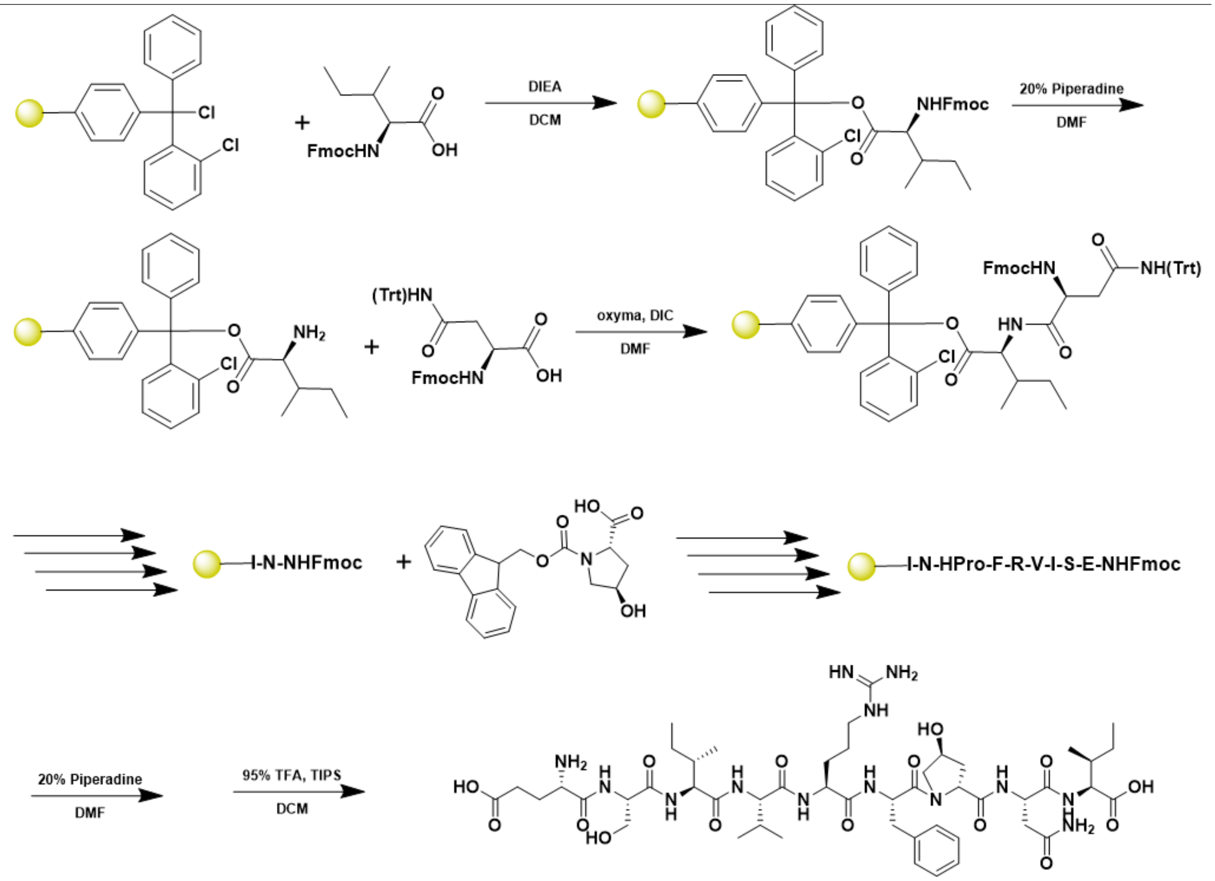


A 25 mL peptide synthesis vessel charged with 2-hlorotrityl chloride resin (0.25mmol) was added Fmoc-L-isoleucine-OH (1.1 eq, 0.275 mmol) and DIEA (3 eq, 0.75 mmol) in dry DCM. The resin was agitated for 1 h at ambient temperature and washed with MeOH and DCM (3 x each). The Fmoc group was removed by using a 20% piperidine solution in DMF for 30 min at ambient temperature, then washed as before. Fmoc-L-asparagine(Trt)-OH (5 eq, 1.25 mM) along with Oxyma (5 eq, 1.25 mM) and DIC (5 eq, 1.35 mmol) in DMF was added to the reaction vessel and agitated for 2 h at ambient temperature. The Fmoc removal and coupling procedure was repeated as before using the same equivalencies for the remaining amino acids. For the 6$^{th}$ position of the amino acid, Fmoc was deprotected as described above and Fmoc-L-trans-4-hydroxyproline (5 eq, 1.25 mM) was added along with Oxyma (5 eq, 1.25 mM) and DIC (5 eq, 1.35 mmol) in DMF the reaction vessel. To remove the peptide from resin, a TFA cocktail solution (95% TFA, 2.5% TIPS, and 2.5% DCM) was added to the resin and agitated for 2 hours. The resin was filtered, and the resulting solution was concentrated in vacuo. The peptide was trituated with cold diethyl ether and purified using reverse phase HPLC using $H_2O/CH_3CN$. The sample was analyzed for purity using a Waters 1525 Binary HPLC Pump using a Phenomenex Luna 5u C8(2) 100A (250 x 4.60 mm) column; gradient eluted with $H_20/CH_3CN$. Molecular weight was confirmed using high resolution electrospray ionization mass spectrometry (HRMS, ESI/MS) analyses obtained on an Agilent 6545B Q-TOF LC/MS equipped with 1260 infinity II LC system with auto sampler. The final peptide product was lyophilized and stored at -20$^o$C until further use.

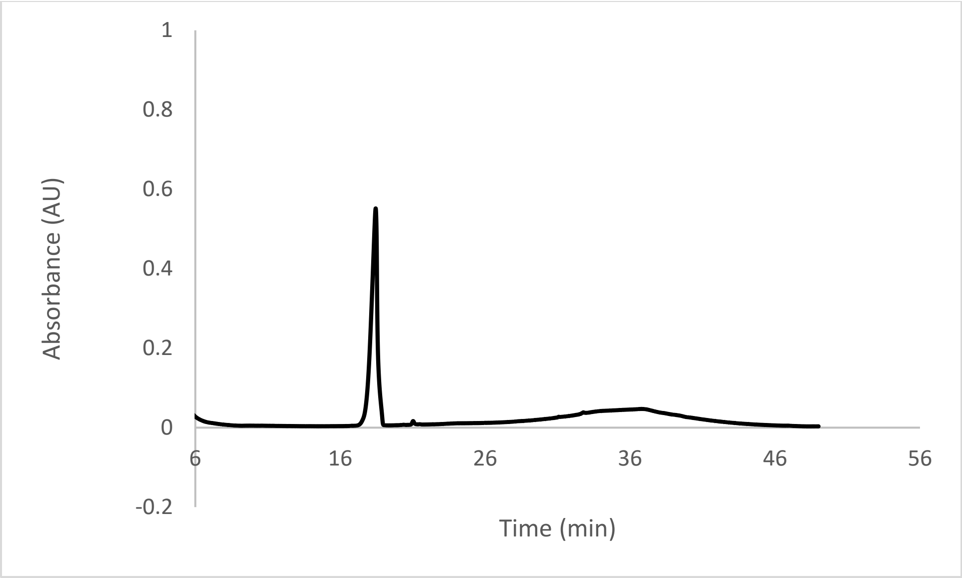


ESI-MS calculated [M]: 1089.5819, found 1089.5838

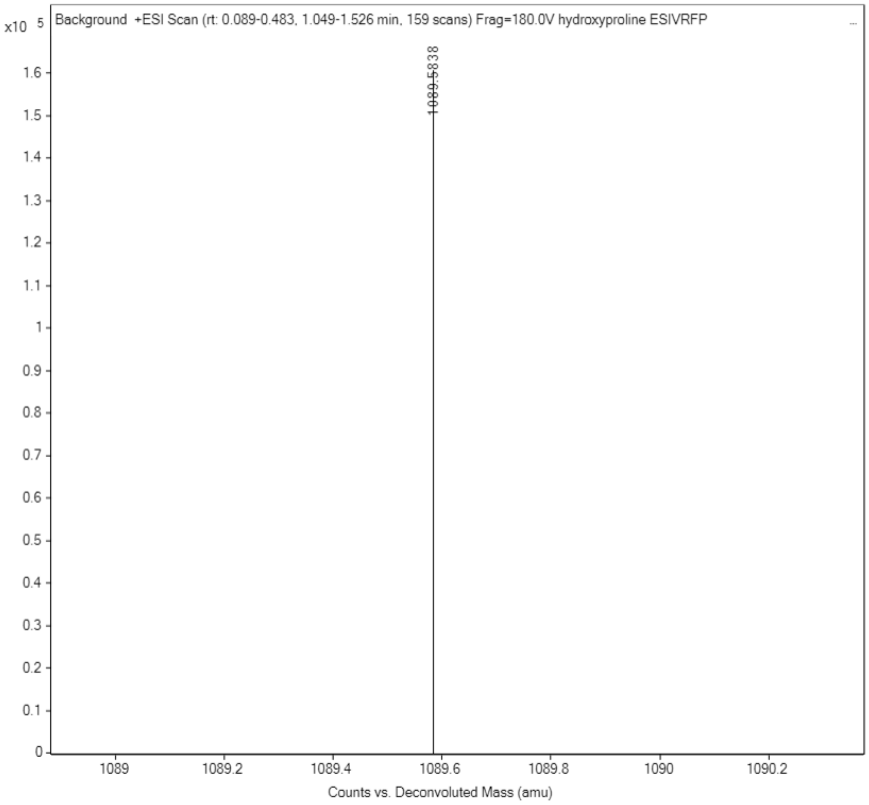
